# An integrative single-cell and spatial transcriptomics atlas highlights candidate regulatory factors in the development of gerbera capitulum

**DOI:** 10.64898/2026.07.05.736605

**Authors:** Yuan Gao, Fan Li, Chunlian Jin, Dick de Ridder, Richard Immink, Yue Sun, Pinli Hu, Yinzhu Cao, Haojing Shao, Aalt D.J. van Dijk, Jihua Wang

## Abstract

In Asteraceae species, the capitulum is a compact inflorescence, featuring a characteristic reproductive structure. Despite the identification of a few key regulatory factors, the transcriptome-level information on the developing capitulum remains limited. Here, we applied single-cell and spatial transcriptome sequencing to investigate the developing Gerbera hybrida’s capitulum during floret differentiation. We obtained a transcriptomics atlas encompassing different stages of the Gerbera capitulum and analyzed the cellular and spatial dynamics of gene expression. Using marker gene expression and GO enrichment of cluster-specific DEGs, we annotated putative cell types and described changes in gene expression across sampled stages, potentially associated with ongoing developmental processes. We detected activity of previously undescribed MADS-box genes and defined their spatial expression patterns. Notably, the MADS-box gene *GAGL12* was found to be enriched in the putative capitulum phloem cells. The *GAGL12* protein was shown in yeast two-hybrid assays to interact with several other MADS-domain proteins with hypothesized functions in vasculature development, and further detailed in silico analyses supported a candidate role in the development of capitulum vasculature. Altogether, we provide integrative and dynamic transcriptomic insight into capitulum and floret development and lay a basis for future functional studies of the control and development of this intriguing reproductive structure.

## Introduction

Flowers are essential organs in angiosperms to attract pollinating animals or other natural forces to exchange germ cells between individuals and then produce seeds to spread new plants. Various features have evolved to make flowers more appealing to pollinators, such as insects and birds, in different environments. These features include variable and vivid colors, large flower sizes, and strong fragrances. Asteraceae is one of the most evolutionarily successful flowering plant families, encompassing around 1,600 genera with more than 23,000 species(Van Huylenbroeck, 2018).

The capitulum, also known as the flowerhead or anthodium, is the defining feature of the Asteraceae family. It is a highly specialized inflorescence composed of many flowers in a tight arrangement and works as a compact inflorescence. The capitulum displays a beautiful topological pattern mimicking the shape of a single flower. The Asteraceae capitula have some commonalities, including a large and dome-shaped receptacle that holds the flowers, which are commonly known as florets. The outer florets are marginal zygomorphic ray florets with colorful, large petals, and centrally, sexually perfect and fertile disc florets develop.

The outstanding appearance of the capitulum attracts not only pollinators but also humans, which makes Asteraceae plants highly popular fresh-cut flowers in the global flower market. Within the Asteraceae, gerbera is an economically important ornamental crop because of its capitulum size, great variety of colors, and multi-styled ray floret shapes. Multiple species within the genus *Gerbera*, originating from South Africa, Asia, South America, and Tasmania, are generally called ‘gerbera’. Among them, *Gerbera hybrida*, an interspecific hybrid between *G. jamesonii* and *G. viridifolia*(Deng and Bhattarai, 2018), is the most cultivated and was introduced to Europe in the 19th century. Current breeding strategies for *G. hybrida* focus on new colors and morphological features, such as curved or spinning ray florets. In the current omics era, an integrative molecular landscape of capitulum development could be obtained to support future breeding-oriented studies of *G. hybrida* ornamental traits.

MADS-box genes play essential roles in different stages and processes of capitulum and floret development. So far, the expression patterns of six SQUA subfamily genes (*Gerbera SQUAMOSA-LIKE1-6* [*GSQUA1-6*]), three AP3/DEF subfamily genes (*Gerbera DEFICIENS-LIKE 1-3* [*GDEF1-3*]), one PI/GLO subfamily gene (*Gerbera GLOBOSA-LIKE 1* [*GGLO1*]), two AG subfamily genes (*Gerbera AGAMOUS-LIKE 1-2* [*GAGA1-2*]), eight AGL2 subfamily genes (*Gerbera REGULATOR OF CAPITULUM DEVELOPMENT 1-8* [*GRCD1-8*]), and two TM3 subfamily genes (*Gerbera SUPPRESSOR OF OVEREXPRESSION OF CO1-2* [GSOC1-2]) have been studied. Among these genes, some have well-defined functions that have been experimentally confirmed. For example, overexpression of *GSQUA2* has been proven to accelerate capitulum initiation without significant effects on floret identity(Ruokolainen, Ng, Suvi K. Broholm, Albert, Elomaa, *et al*., 2010); GGLO1 and GDEF2 proteins show strong interaction and work together to specify petal and stamen identity(Ruokolainen, Ng, Albert, *et al*., 2010); *GRCD2* controls inflorescence meristem determinacy, and the loss of its function causes indeterminate capitulum growth and extra florets(Uimari *et al*., 2004). Nevertheless, orthologs of the majority of MADS-box genes from various subfamilies are yet to be characterized in G. hybrida, leading to an incomplete understanding of the regulatory roles of MADS-box genes in capitulum and floret development.

Single-cell RNA sequencing (scRNA-seq) generates cell-type- or cell-state-associated expression signatures, which are critical to understanding the complex gene regulatory networks that control plant development. In recent years, rapid advances in scRNA-seq techniques have enabled us to investigate gene expression patterns of numerous cell types in various plant species(Dorrity *et al*., 2021; Ryu *et al*., 2019; Shaw *et al*., 2021). Complementing scRNA-seq, spatial transcriptomics has emerged as a powerful approach to retain the spatial context of gene expression, enabling us to map where specific transcripts are expressed within tissues. Although still nascent in plant research, spatial transcriptomics is proving invaluable for resolving tissue architecture and cellular interactions during development(Liu *et al*., 2023; Nobori *et al*., 2023; Xia *et al*., 2022). By combining scRNA-seq with spatial transcriptomics in the capitulum, our study provides a spatially resolved transcriptomics resource, facilitating the analysis of Gerbera capitulum development and unlocking its unique characteristics and associated gene regulatory network.

Specifically, to facilitate an integrative transcriptomic-level description of gene expression during the floret-differentiation stage of the *G. hybrida* capitulum, we applied scRNA-seq and Stereo-seq sequencing on capitulum samples spanning successive floret-differentiation stages. This allowed us to generate a transcriptomic atlas and, for various cell types, to analyze their proportions and distributions across the capitulum in different developmental stages. Subsequently, we focused on the expression patterns of MADS-box genes, which are considered essential factors controlling the orchestration of inflorescence and early flower development. Our work constructed an atlas of the floret-differentiation stages of *G. hybrida* capitulum and discovered several candidate genes that may be associated with floret differentiation, the development of the vascular system in the capitulum, and floret organ identity specification.

## Results

### Single-cell RNA sequencing

Three capitula at successive floret-differentiation stages were sampled for single-cell measurements (Fig. 1a). The measured stages (S1–S3), with the capitula width (perpendicular to the floral axis) measured inclusive of the involucral bracts prior to their removal, represent stages of capitulum patterning where individual floral meristems (FMs) are already distinct or rapidly expanding. Importantly, the involucral bracts (green leaf-like tissue surrounding the capitulum) would lead to the failure of protoplast culturing, so they were removed in all three samples in advance. After sequencing, 104.3 Gbp, 114.6 Gbp, and 121.2 Gbp of paired-end reads from three capitulum samples were obtained, respectively. Preprocessing by Cell Ranger generated three raw barcode matrices containing 32,765, 29,726, and 39,513 called barcodes for the three datasets, respectively. These numbers represent the total barcodes detected before quality control rather than recovered cells, with the large majority corresponding to low-count, near-empty droplets. After excluding these low-count barcodes, 8,241, 9,043, and 8,506 high-quality cells with a total expression count of at least 2,000 were retained from stages S1, S2, and S3 for further analysis.

**Figure 1.**
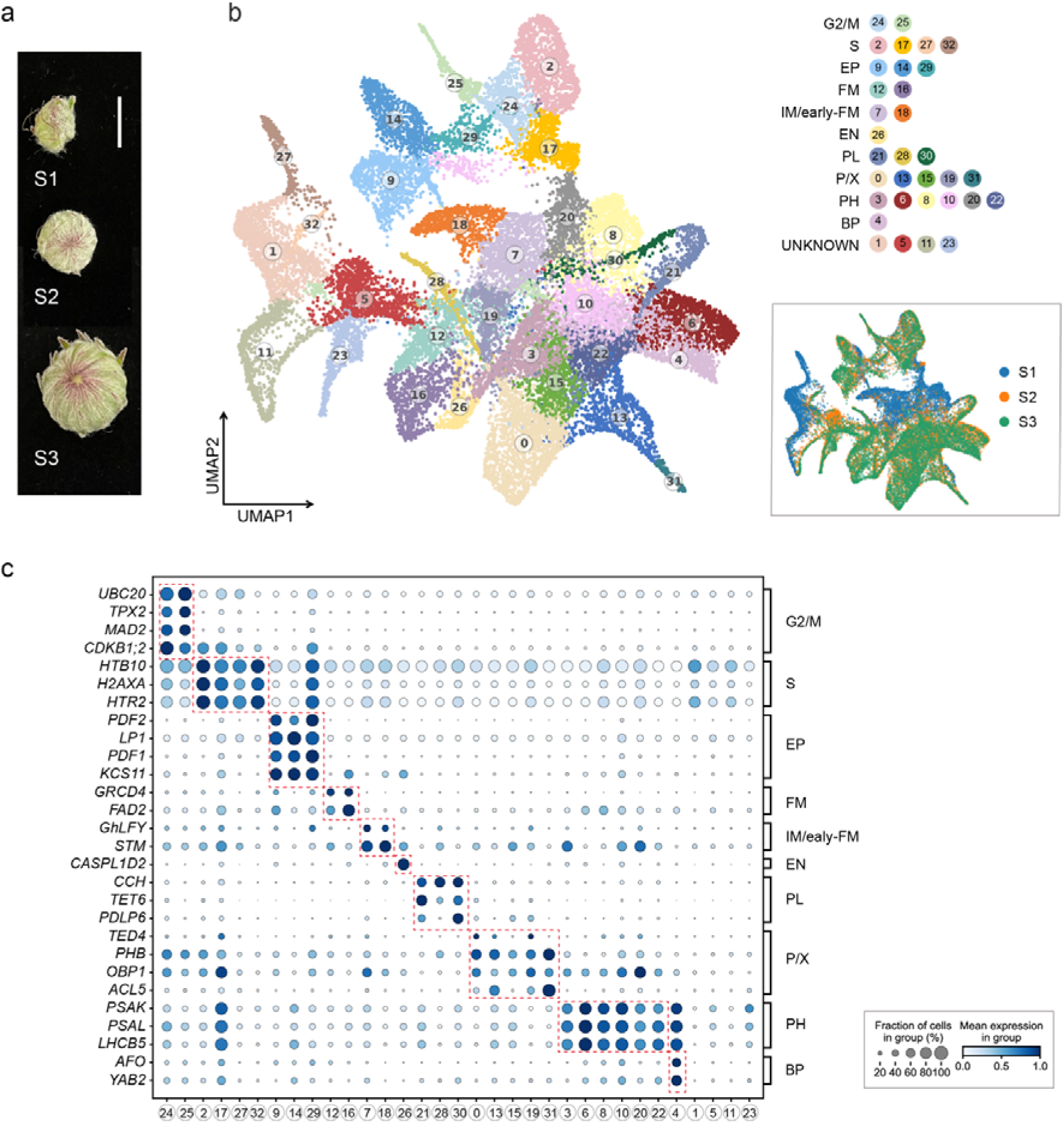
Single-cell transcriptome sequencing results. (a) Representative capitulum buds selected for protoplast isolation and scRNA-seq. The size of the capitulum increases from S1 to S3, related to its developmental stage. The scale bar is 10 mm. (b) UMAP plot of Leiden clustering result and manually annotated cell types of processed single-cell transcriptomes measured in all three samples. In total, we detected 33 clusters, which were then classified into 11 cell types (including unknown cells). The developmental stage of each cell is shown on the bottom right. (c) The expression of typical marker genes (row, named based on similarity with genes in *A. thaliana*) in each cell type (column). The color of the dot indicates the average expression level of a particular gene across different cell types (maximum count scaled to 1). The size of the dot indicates the proportion of cells within a cluster that express the gene.

We obtained 33 cell clusters at the chosen clustering resolution, a resolution that balanced cluster granularity against over-partitioning. Then we aggregated the clusters into 10 major identified putative cell types on the basis of marker-gene concordance and GO enrichment: G2/M phase cells, S phase cells, epidermis cells, inflorescence-meristem/early-floral-meristem cells, floral meristem cells, endodermis cells, phloem cells, procambium/xylem cells, photosynthetic cells, and bract primordial cells (Fig. 1b). The cell types were inferred based on the specific expression of published or homolog-derived marker genes from previous studies (Fig. 1c) and GO-term enrichment of cluster-specific differentially expressed genes (DEGs) (supplemental material), so that each annotation reflected both candidate marker expression and the broader functional program of the cluster:

#### G2/M-phase (G2/M) cells

Clusters 24 and 25 have high expression of classic G2/M markers-*UBIQUITIN-CONJUGATING ENZYME 20* (*UBC20*)(Fülöp *et al*., 2005), *MITOTIC ARREST DEFICIENT 2* (*MAD2*)(Satterlee *et al*., 2020), and *TARGETING PROTEIN FOR XKLP2* (*TPX2*)(Satterlee *et al*., 2020)-indicating that they are actively progressing through the G2/M transition of the cell cycle. Consistent with this, DEGs in these clusters were enriched for mitotic-related GO terms such as “mitotic sister chromatid segregation” (GO:0000070) and “microtubule cytoskeleton organization” (GO:0000226), which are typical biological processes related to the G2/M phase.

#### S-phase (S) cells

Genes encoding histone assembly factors (*HTB10*(Zambrano-Mila *et al*., 2019), *H2AXA*(Zhou *et al*., 2015), *HISTONE THREE RELATED* 2 (*HTR2*)(Probst *et al*., 2020)) appeared broadly across most clusters but showed peak expression in clusters 2, 17, 27, and 32. DEGs in these clusters were significantly enriched for “cell cycle” (GO:0007049), “DNA metabolic process” (GO:0006259), and “chromatin remodeling” (GO:0006338), consistent with S-phase activity and DNA replication.

#### Epidermal (EP) cells

Clusters 9, 14, and 29 showed robust expression of *PROTODERMAL FACTOR 1-2* (*PDF1-2*)(Javelle *et al*., 2011), *LIPID TRANSFER PROTEIN 1* (*LP1*)(Thoma *et al*., 1994), *3-KETOACYL-COA SYNTHASE 11* (*KCS11*)(Nobusawa *et al*., 2013), all of which are protoderm/epidermis markers and known to regulate cuticle formation and epidermal cell differentiation. Also, in all three clusters, the GO term “wax biosynthetic process” (GO:0010025) was enriched.

#### Inflorescence-meristem/early-floral-meristem (IM/early-FM) cells

Clusters 7 and 18 both exhibited high expressions of *GhLFY*(Zhao *et al*., 2016) and *SHOOT MERISTEMLESS* (*STM*)(Balkunde *et al*., 2017; Long and Barton, 2000) that are associated with inflorescence meristem identity and activity, respectively. However, GO enrichment analysis on DEGs did not yield specifically enriched terms that could be associated with inflorescence identity.

#### Floral meristem (FM) cells

In clusters 12 and 16, we saw strong upregulation of floral meristem marker *GRCD4*(Ruokolainen, Ng, Suvi K. Broholm, Albert, Elomaa, *et al*., 2010) (an AGL2-type MADS-box gene) and the non-canonical marker *FATTY ACID DESATURATION 2* (*FAD2*). GO analysis for DEGs in these clusters indicated enrichment for “specification of floral organ identity” (GO:0010093), “floral organ formation” (GO:0048449), and “floral organ morphogenesis” (GO:0048444), supporting the annotation of these two clusters as putative floral meristem cells.

#### Endodermis (EN) cells

Cluster 26 showed exclusive, high expression of *CASPL1D2*, a Casparian strip-associated protein known to be restricted to endodermal cells(Champeyroux *et al*., 2019). This one-gene signature supported a tentative annotation of cluster 26 as endodermis cells.

#### Phloem (PL) cells

Clusters 21, 28, and 30 all expressed phloem markers *COPPER CHAPERONE* (*CCH*)(Mira *et al*., 2001) and *TETRASPANIN6* (*TET6*)(Wang *et al*., 2015), which in Arabidopsis are expressed in the phloem and associated vascular tissues. However, *PLASMODESMATA-LOCATED PROTEIN* 6 (*PDLP6*)—a second phloem marker—was present only in clusters 21 and 30 (not in 28). Because *CCH* and *TET6* are closely associated with phloem identity, we still consider clusters 21, 28, and 30 as phloem cell populations, while noting that cluster 28 may represent a divergent subset.

#### Bract primordia (BP) cells

Cluster 4 uniquely co-expressed *FIL* and *YAB2*, two YABBY family genes known to mark the abaxial side of emerging organ bract primordia(Goldshmidt *et al*., 2008; Siegfried *et al*., 1999). DEGs in cluster 4 were enriched for photosynthesis-related GO terms, including “chlorophyll binding” (GO:0031409), “photosystem” (GO:0009521), and “photosynthesis” (GO:0015979), which indicates that these cells correspond to developing leaf and/or organ primordia with an active photosynthetic program on their abaxial surfaces.

#### Procambium/xylem (P/X) clusters

DEGs found in Clusters 0, 13, 15, 19, and 31 include *TRACHEARY ELEMENT DIFFERENTIATION-RELATED 4* (*TED4*) and *ACAULIS 5* (*ACL5*). *TED4* has been reported to be highly expressed in both procambium and xylem(Endo *et al*., 2018), while *ACL5* is found only in xylem(Liu *et al*., 2022); because their expression overlaps these vascular lineages, we could not resolve a single specific cell type within these clusters. They likely represent a mix of procambial and early xylem cells.

#### Photosynthetic mesophyll-like (PH) cells

Clusters 3, 6, 8, 10, 20, and 22 uniformly expressed *PHOTOSYSTEM I SUBUNIT K* (*PSAK*)(Mazor *et al*., 2017) and *PHOTOSYSTEM I SUBUNIT L* (*PSAL*)(Mazor *et al*., 2017), photosynthetic markers PSI subunits; and *LHCB5*(Ballottari *et al*., 2013), a PSII light□harvesting protein. Although these genes were also expressed in clusters 4 and 17, the magnitude and consistency of expression across clusters 3, 6, 8, 10, 20, and 22 made us designate these cells as mesophyll-type or other highly photosynthetic cell populations.

Cells in clusters 1, 5, 11, and 23, which cover 15.46% of all valid cells, have generally low total transcript counts and few cluster-specific DEGs. We could not identify markers from the literature that would allow confident assignment for these clusters, so they remain unclassified.

### Spatial RNA sequencing

The single-cell transcriptomes help to identify the cell-type composition of the capitulum in different developmental stages. However, these transcriptomes lack information about the spatial context of the measured cells, which is essential for studying such a complex biological structure. Spatial transcriptomes, on the other hand, do provide such context. Here, we chose Stereo-seq to measure the spatial transcriptomics of the capitulum. 19 capitula with sizes roughly covering S1 to S3 were selected for tissue section and Stereo-seq spatial transcriptome sequencing. Four spatial gene expression matrix files (SP1-SP4), as well as the corresponding optical images (Fig. 2a), were generated during the sequencing process. One chip (SP2) had to be discarded due to ultra-low expression count. Among all capitulum samples, SP1_d is of the best quality, i.e., it has the highest expression counts and least mRNA diffusion.

**Figure 2.**
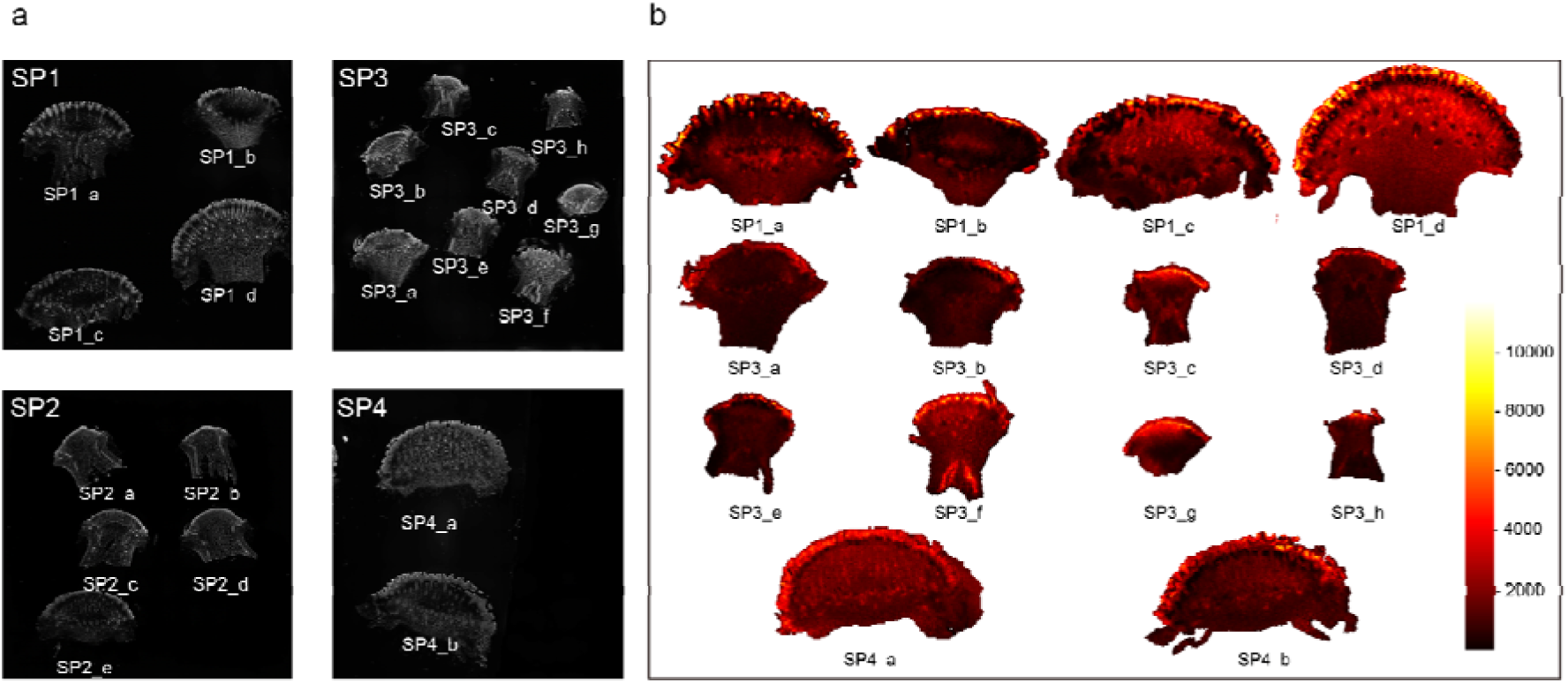
Spatial transcriptome sequencing results and integration with single-cell data. (a) Optical images of tissue sections corresponding to spatial transcriptomes. (b) Heatmap of total gene expression counts of all manually segmented spatial transcriptome samples. All samples in SP2 were discarded due to ultra-low expression counts.

We chose to work with pooled 50×50 pixel spatial domains for further analysis on the three selected chips (one chip was abandoned due to the low quality), which contained a total of 14 individual capitulum samples (Fig. 2b). For some marker genes used previously in single-cell analysis, the spatial transcriptomes showed enrichment confined to specific spatial domains. For instance, in spatial sample SP1_d, the spatial domains that highly expressed two marker genes for epidermis, *PDF1* and *LP1*(Ruokolainen *et al*., 2011), are aligned with the known location of epidermis in the capitulum. Two marker genes of procambium/xylem, *TED4* and *ACL5*, exhibit a high spatial similarity with the simulated distribution of vasculature in *G. hybrida*(Owens *et al*., 2024) (Sup. Fig. S3).

The spatial transcriptomes contain both spatial and sequence information about each read, but they cannot be segmented precisely at the cellular level, due to the inaccurate match between optical images and spatial expression profiles, non-negligible mRNA diffusion, and the data sparsity of reads. In order to investigate the spatial distribution of different cell types, we combined the advantages of both analyses by integrating the spatial and single-cell transcriptomes using cell2location(Kleshchevnikov *et al*., 2022) to infer the spatial cell-type distribution. In total, 3 integration results were generated (Sup. Fig. S4/S5/S6). These integration results were named after the combination of spatial and single-cell data (e.g. SP1_d-S3 indicates the integration of spatial data SP1_d and single-cell data S3). The primary difference between these integrations and the original spatial transcriptomes is that each pseudo-cell is no longer represented by a gene expression profile but by a cell-type distribution array

### Transcriptomic dynamics of the developing capitulum

The distribution of cell types in different samples changed significantly with developmental stages (Fig. 3a). For example, the cell count in IM/early-FM dropped to nearly zero as the capitulum grew, while that in FM increased rapidly. This matches well with previous studies indicating that, during the formation of the capitulum, the IM/early-FM cells on top of the receptacle are gradually converted to FM cells from the periphery to the central zone based on regulation by internal and external factors(Elomaa *et al*., 2018; Zhang *et al*., 2021). The count of BP cells also increased with the developmental stages, in line with the gradually enlarged bracts. Within PH cells, different clusters have different tendencies.

**Figure 3.**
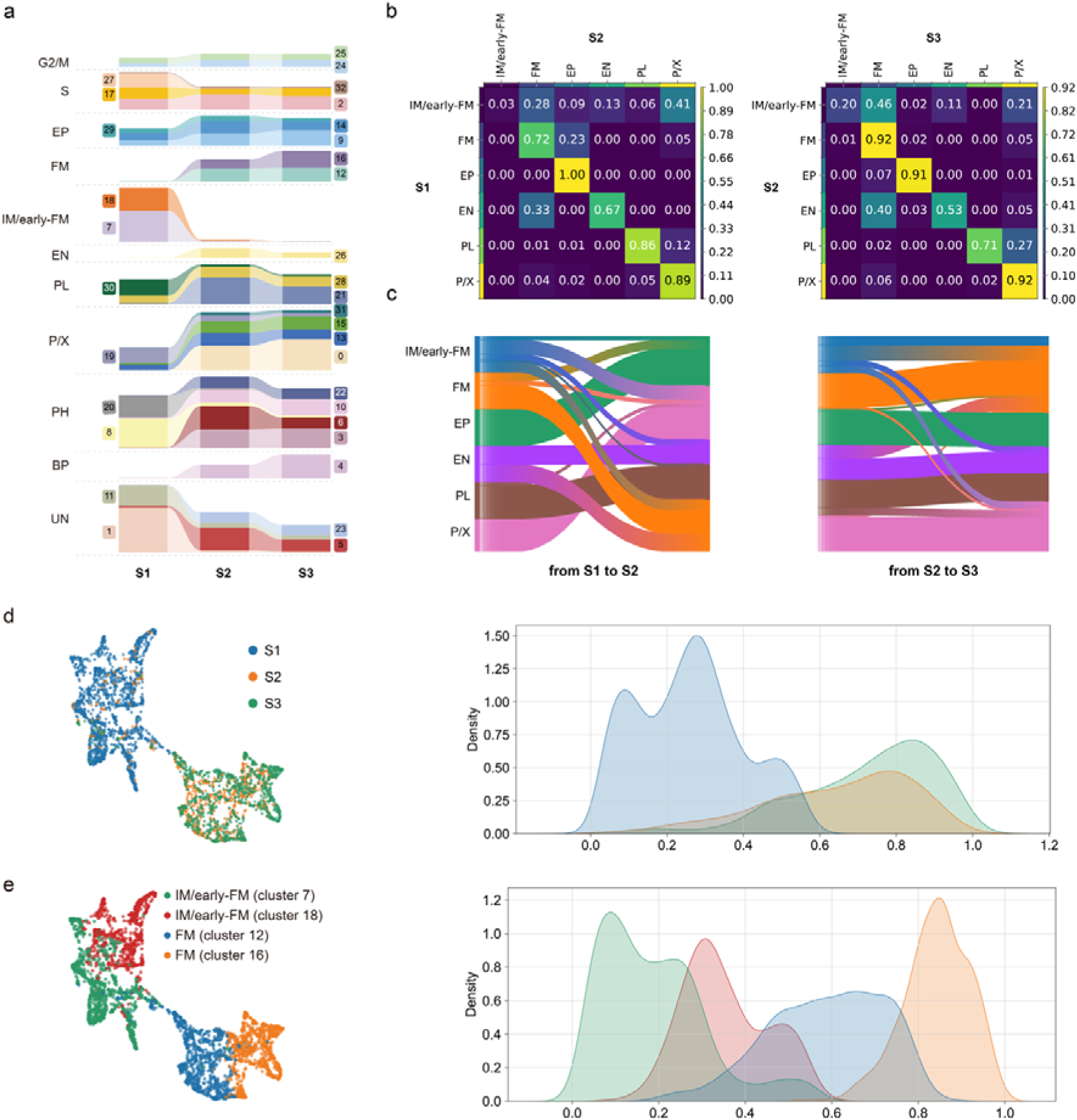
Developmental dynamics of the developing capitulum. The abbreviations of cell types are listed on the bottom right. (a) Change in counts of cell types (indicated by colors) in different developmental stages. The numbers on the right or left indicate clusters. (b) Transition matrices of descendancy probabilities between consecutive stages S1, S2, and S3. Each entry gives the probability that a cell of a given type at the earlier stage (rows) gives rise to each cell type at the later stage (columns); each row is normalized to sum to one. (c) Sankey diagrams visualizing the same cell-type transitions for (S1, S2) (left) and (S2, S3) (right). Bands connect cell types between consecutive stages, with width proportional to the transition probability and color denoting cell-type identity. (d) IM/early-FM and FM cells in different developmental stages. Left, cells in a 2D UMAP embedding; right, Cell distribution density of different stages as a function of pseudotime. (e) Subgroups of IM/early-FM and FM cells. Left, cells in a 2D UMAP embedding; right, Cell distribution density of different subgroups as a function of pseudotime.

We applied moscot(Klein *et al*., 2025) to disentangle the development relationships of different cell types in three stages. Here, we focused on the cell types that contribute the most to the formation of capitulum structure, including IM/early-FM cells, floral meristem (FM) cells, epidermis (EP) cells, endodermis (EN) cells, phloem (PL) cells, and procambium/xylem (P/X) cells. The transition matrix (Fig. 3b) from S1 to S2 shows the differentiation of IM/early-FM cells into IM/early-FM, FM, EN, and P/X cells, consistent with the expected behavior of a meristematic population. Meanwhile, the cell transition crossing different cell types occurred less from S2 to S3.

To study the transition between meristematic cell populations during floret differentiation, we zoomed in on the transition from the IM/early-FM population—defined by expression of the meristem markers *GhLFY* and *STM* without an active floral-organ-identity program—to floral meristem (FM) cells in the capitulum. During the formation of the capitulum, the IM/early-FM cells on top of the receptacle are gradually converted into FM cells from the periphery to the central zone based on regulation by internal and external factors(Elomaa *et al*., 2018; Yu *et al*., 1999; Zhang *et al*., 2021) (Fig. 3d). FM cells then start the ontogeny of flower organs in the sequence of floral whorls(Thomson and Wellmer, 2019). IM/early-FM cells were mostly found in S1, and FM cells in S2 and S3 (Fig. 3e).

### Spatial expression patterns of MADS-box genes

MADS-box transcription factors play vital roles in floral development, and previous research has shown a wide involvement of MADS-box genes in the development of the capitulum. Our single-cell and spatial transcriptomics data provide the opportunity to further detect the expression patterns of MADS-box transcription factors in the capitulum during floret differentiation. By phylogenetic analysis, we found 90 MADS-box genes in the genome of *G. hybrida* (Sup. Fig. S7). At the late development stage of the capitulum (S3), the spatial expressions of MADS-box genes can be roughly classified into four patterns: **Pattern 1**, expression throughout the whole capitulum; **Pattern 2**, at the top of the capitulum; **Pattern 3**, in the middle of the capitulum; **Pattern 4**, at the base of the capitulum (Fig. 4a). Convolution-based spatial correlation analysis and hierarchical clustering quantified the similarity among the spatial expression of MADS-box genes (Fig. 4b).

**Figure 4.**
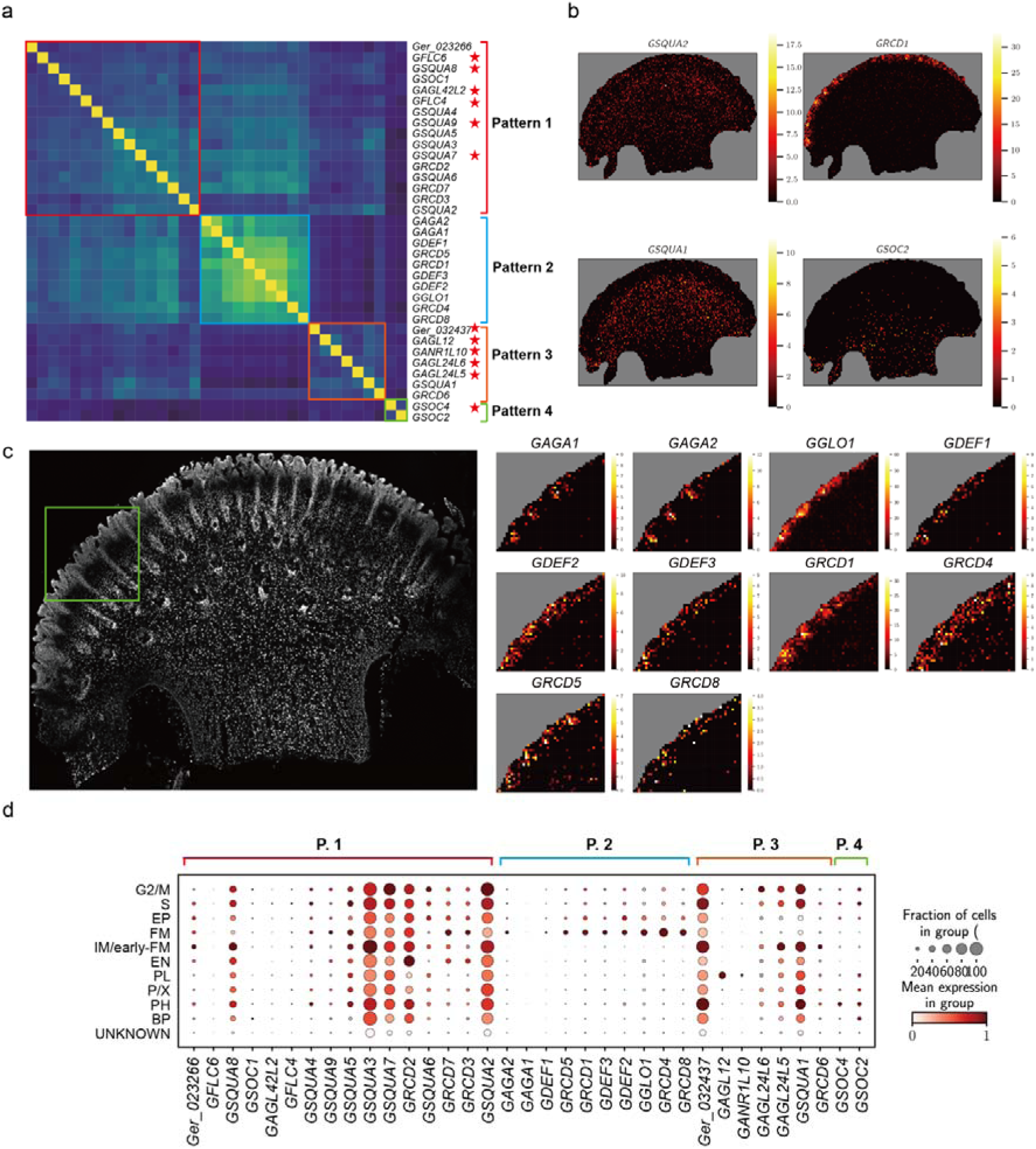
Spatial expression of MADS-box genes. (a) Heatmap of the spatial correlation matrix of MADS-box genes given a minimum expression count of 100. The color indicates the spatial correlation between genes. The genes are arranged in the same order on x-axis and y-axis. New MADS-box genes are marked by a star on the right. (b) Four typical spatial expression patterns of MADS-box genes. **Pattern 1**, whole capitulum; **pattern 2**, top; **pattern 3**, middle; **pattern 4**, bottom. (c) Detailed spatial expression of pattern 2 MADS-box genes. Left, optical image of sample SP1_d. Green box shows the range for detailed study. Right, the zoomed-in heatmaps of MADS-box genes in pattern 2. (d) The expression distribution of MADS-box genes in the single-cell transcriptome. The color of the dot indicates the average expression level of a particular gene across different cell types (scaled to [0, 1]). The size of the dot indicates the proportion of cells with a cell type that expresses the gene.

Pattern 1 represents the widest spatial distribution in the capitulum, which suggests the potential involvement of these genes in many different processes during capitulum and floret development. Most SQUA subfamily MADS-box genes, *GSQUA2/3/4/5/6,* were already proven to be widely expressed during inflorescence development, including vasculature and floral organs(Ruokolainen, Ng, Suvi K Broholm, Albert, Elomaa, *et al*., 2010), although the knowledge about their expression dynamics was still fragmented. Here, these *GSQUA* genes and three newly identified SQUA subfamily MADS-box genes, *GSQUA7/8/9*, were all detected to be expressed following pattern 1. Previously, it has been hypothesized that the high number of *GSQUA* genes facilitates the development of the complex capitulum structure(Ruokolainen, Ng, Suvi K. Broholm, Albert, Elomaa, *et al*., 2010), which is consistent with the observed extensive expression of these genes in the floret-differentiation stages (Fig. 4a/b/d) and differences in cell-type specific abundance and spatial patterns. The *SEPALLATA*-like MADS box genes *GRCD2/7* were found to share a high nucleotide sequence similarity, as well as a fully overlapping expression domain, suggesting mutual functional redundancy(Zhang *et al*., 2017); our analyses confirmed their overlapping expression domains at spatial domain level. We also detected *GFLC4/6*, two FLC-type genes that were never studied in *G. hybrida* before. In *A. thaliana*, *FLC* was first discovered to mostly act in the regulation of flowering time, as a repressor of two genes promoting flowering(Deng *et al*., 2011; Helliwell *et al*., 2006; Searle *et al*., 2006; Sheldon *et al*., 1999); further study revealed its broad involvement in the regulation of e.g., the *SEP3* MADS box gene, which fulfils key roles in subsequent flower developmental processes (Deng *et al*., 2011). The identified co-expression with the SQUA subfamily genes during capitulum development suggests similar activity for these two *G. hybrida* FLC-type genes.

Pattern 2 covers the upper surface of the capitulum, where floral meristems and floral organs are located. All the genes contained in this pattern have been studied and classified according to the ABC model for floral organ identity determination(Causier *et al*., 2010; Teeri *et al*., 2006). *GAGA1/2* were proven to be key factors of the determination of stamen and carpel identity (C-function); *GGLO1*, *GDEF2* (B-function), and *GRCD1/4/5* (E-function) were validated to play essential roles in specifying the identity of petals and stamens(Broholm *et al*., 2010; Zhang *et al*., 2017). Among the genes expressed in pattern 2, the spatial expression levels of two *AG* homologs, *GAGA1/2*, are different although overlapping with those of *GGLO1*, *GDEF*1/2/3, and *GRCD1/4/5/8* (Fig. 4c). The single-cell expression pattern of pattern 2 MADS-box genes (Fig. 4d) shows that they are highly expressed in floral meristem and epidermal cells, which distribute on top of the floral meristem according to cell2location results (Sup. Fig. S3).

The pattern 3 genes are more highly expressed in the middle zone of the capitulum, where tissues grow to support capitulum expansion and therefore support floret development. Most of the MADS-box genes in this pattern were detected for the first time, except for *GSQUA1* and *GRCD6*. *GSQUA1* is an SQUA subfamily MADS-box gene, and previous studies showed that it is highly enriched in the vasculature of the capitulum receptacle and later during floret development in petals(Ruokolainen, Ng, Suvi K. Broholm, Albert, Elomaa, *et al*., 2010). In our analyses, *GSQUA1 was* expressed widely in most parts of the capitulum except for the epidermis and floral meristems. Among the newly detected MADS-box genes, *GAGL12* and *GANR1L10* show a uniquely high expression in phloem (Fig. 4d), while the other genes are expressed in most cell types except for epidermis and floral meristems.

Compared with the other patterns, pattern 4 is particular in terms of its spatial distribution and very limited gene numbers, containing only the *TM3*-clade genes *GSOC2* and *GSOC4*. According to previous studies in e.g., *A. thaliana*(Immink *et al*., 2012), the *TM3*-clade gene *SUPPRESSOR OF OVEREXPRESSION OF CONSTANS1 (SOC1)* acts as an inducer of flowering and is crucial for the regulation of flowering time. However, recent investigations in petunia and tobacco suggest that *TM3*-clade genes activate photosynthesis and heat-shock-associated genes, resulting in improved photosynthesis and heat tolerance(Ning *et al*., 2021). The shared spatial expression pattern of *G. hybrida GSOC2* and *GSOC4* points to potential redundancy or joint activity.

### *GAGL12* as a phloem-enriched candidate regulator of capitulum vasculature

The development of phloem in the capitulum has not been well studied from a transcriptomics perspective. In single-cell data, the putative phloem cells are represented by clusters 28, 30, and 21, which specifically express two marker genes, *CCH*(Mira *et al*., 2001) and *TET6*(Wang *et al*., 2015). Here, we assign cluster 28 to phloem_a, cluster 30 to phloem_b, and cluster 21 to phloem_c. Another marker gene, *PDLP6*(Kim *et al*., 2021), is detected in phloem_b and phloem_c but not in phloem_a. A previous study showed that *PDLP6* is uniquely expressed in the phloem of mature *Arabidopsis thaliana* leaves, likely in sieve elements, a fully differentiated phloem cell type, to regulate the functioning of plasmodesmata(Li *et al*., 2024). According to RNA velocity analysis, phloem cells exhibit two separate inferred development trajectories (Fig. 5a&b). Cells in both trajectories transit from states in S1 to S2 and S3. Also, two independent terminal states were detected using cellrank2 (Fig. 5c). phloem_b is predicted to be related to phloem_c, while cells in phloem_a show stable cell states over the different developmental stages (Fig. 5d&e). Also, based on the cell2location result (Fig. 5h), the putative cell spatial distribution of phloem_a is slightly different from those of phloem_b and phloem_c.

**Figure 5.**
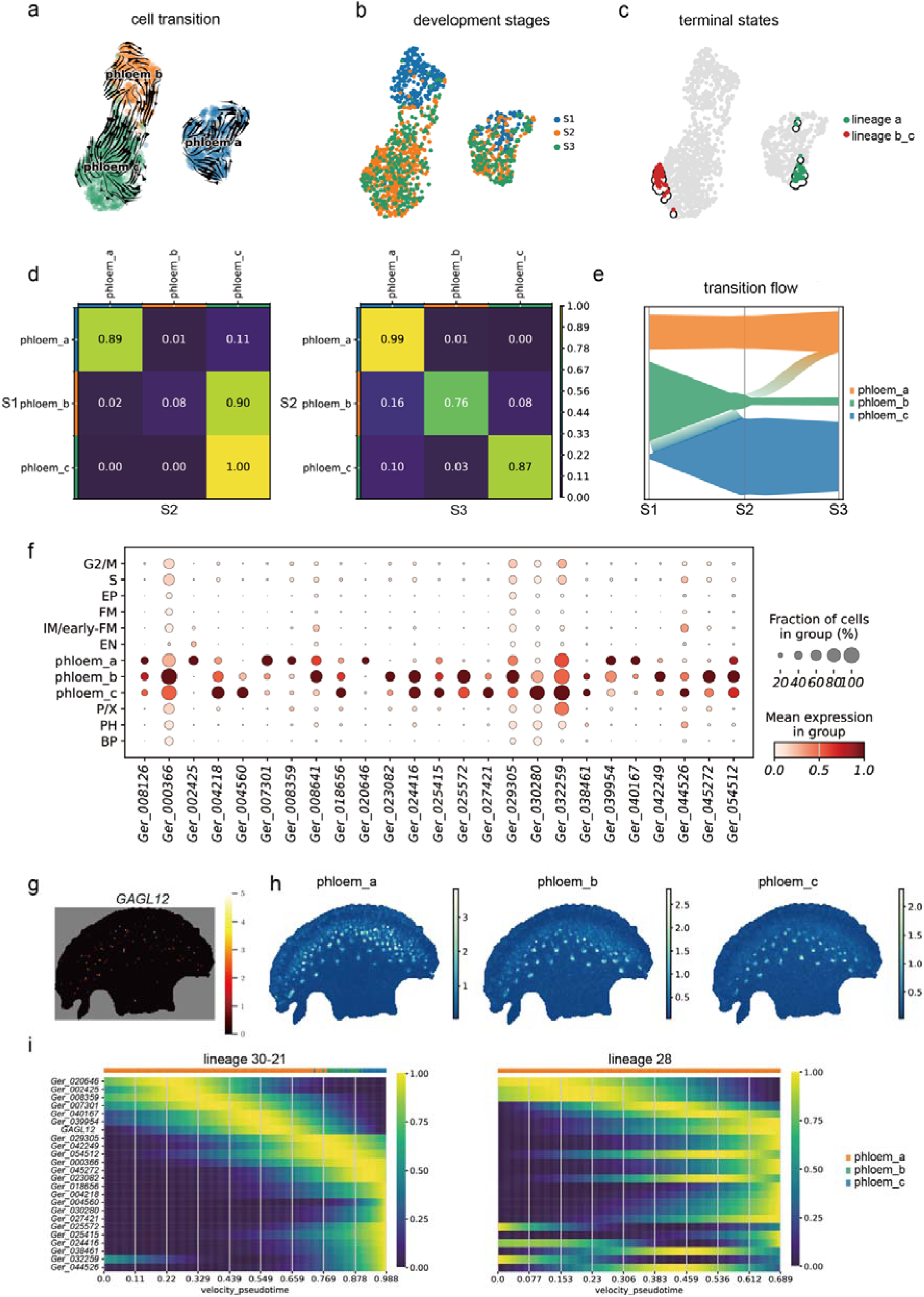
Phloem development and *GAGL12* expression in capitulum. (a) Cell transition stream of phloem cells in the single-cell atlas. The phloem cells are divided into three sub-groups: phloem_a (Leiden cluster 28), phloem_b (Leiden cluster 30), and phloem_c (Leiden cluster 21). (b) The development stages of phloem cells. (c) Terminal state of cell lineages a (containing phloem_a) and b_c (containing phloem_b and phloem_c). (d) Cell type transition matrix from S1 to S2 (left) and S2 to S3 (right). The values show the probability of transition from cell cluster in the column to the row. (e) Cluster transition flow from S1 to S3. (f) The expression of GAGL12 and its putative target genes in each cell type (column). The color of the dot indicates the average expression level of a particular gene across different cell types (maximum count scaled to 1). The size of the dot indicates the proportion of cells within a cell type that express the gene. (g) The spatial distribution of *GAGL12* in SP3_d. (h) The inferred cell count distribution in the spatial transcriptome (SP1_d) of phloem cells (clusters 21, 28, and 30). (i) The expression of *GAGL12* and its putative target genes along the pseudotime in cell lineages a and b_c.

The spatial expression of *GAGL12* is consistent with the expected spatial distribution of phloem cells (Fig. 5h). In *A. thaliana*, *AGL12* is a vital factor for the regulation of root meristem cell proliferation and flowering transition(Montiel *et al*., 2020; Tapia-López *et al*., 2008). Ectopic expression of *VvAGL12* (from *Vitis vinifera*) in *A. thaliana* promotes early flowering, plant growth, and production through changing cell wall architecture(Mao *et al*., 2023). Previous studies also suggested that *AGL12* promotes the overall root vascular system formation in both *Juglans sp.* and *A. thaliana*(Montiel *et al*., 2020). Given the specific expression of *GAGL12* in phloem cells, *GAGL12* is highlighted as a potential key gene regulating the vascular system in the *G. hybrida* capitulum.

It is well known that the proteins encoded by MADS-box genes form homo- or heterodimers and tetramers to regulate a wide range of target genes. To investigate a possible co-regulatory role of GAGL12 with other MADS-domain proteins, we zoomed in on MADS-box genes contained in patterns 1 and 3, whose spatial expression pattern overlaps with *GAGL12*. Yeast two-hybrid experiments were performed to measure the protein-protein interaction capacity of GAGL12 protein (Sup. Fig. S8). This revealed that GAGL12 protein can interact with the MIKC*-type protein Ger_023266, SEPALLATA-like proteins GRCD2/6/7, GAGL24L6, GSQUA1, and GFLC6.

Previously, high expression of *GSQUA1* in vascular bundles of the capitulum was noticed, and a function in vasculature development was proposed(Yu *et al*., 1999). *GFLC6* belongs to the *FLC* subfamily whose genes, such as *MAF1*, *MAF4*, and *FLC*, in *A. thaliana* were found to be expressed preferentially in vasculature, including leaf veins, shoot apical vasculature, and root stele(Gu *et al*., 2013). However, based on *flc* mutant analysis and FLC target gene identification, this is not associated with a role in vasculature development but in the repression of key flowering time regulators expressed in the vasculature(Gu *et al*., 2013).

To further study a possible regulatory function of *GAGL12* in the development of the capitulum vasculature, we first extracted the initial putative GRNs of *GAGL12*. Then we scanned for the occurrence of a CArG-box, the consensus binding motif of MADS-domain proteins, in upstream regions (3kb) of all genes in the genome of *G. hybrida*. The intersection of the genes in the extracted GRN, the genes with a clear CArG-box, and the DEGs in phloem cells were then taken as the final putative target genes of *GAGL12*.

Within the set of inferred target genes, we focused on those with a specifically high expression in phloem cells (Fig. 5f). Based on sequence alignment to genes in TAIR and NCBI, *Ger_000366* is identified as *plasmodesmata callose-binding protein 3* (*PDCB3*), a major member of the PDCB gene family. Though the biological function of *PDCB3* is not yet known, its paralogs *PDCB1* and *PDCB2* were identified as callose-binding proteins and potentially modify cell-to-cell trafficking on plasmodesmata(Simpson *et al*., 2009). As a key feature of plant cells, plasmodesmata are vital for cell-to-cell transportation of various cargos(Sager and Lee, 2018). Interestingly, *PDLP6* (*Ger_045272*) is another putative target gene closely related to plasmodesmata.

## Discussion

In this study, we aimed to disentangle gene expression dynamics during the development of the capitulum of *G. hybrida*. Single-cell and spatial transcriptome sequencing were adopted to construct an atlas of the unbiased gene expression patterns in different developmental stages. Based on the single-cell transcriptome results, we annotated 10 putative cell types in the capitula samples spanning floret-differentiation stages, ranging from younger capitula enriched in putative inflorescence-meristem/early floral-meristem cells to more mature ones with floral meristems covering the surface. We recognize that most cell-type markers have not been experimentally validated as cell-type-specific markers in gerbera capitula. Therefore, our annotations should be interpreted as putative cell-type assignments. However, these assignments were not derived from marker genes alone; they were further supported by GO enrichment of cluster-specific DEGs, which provided transcriptome-wide evidence for the dominant biological programs represented by each cluster. The distribution of different cell types in the single-cell data changes significantly across development, suggesting stage-associated changes in cellular composition and transcriptional state.

The single-cell and spatial transcriptomes provide a wealth of information, making it possible to study developmental dynamics at the single-cell and spatial scale. The single-cell transcriptomes provide more detailed gene expression profiles - higher gene expression counts, gene numbers, and resolution - than currently possible using spatial transcriptomics. The spatial transcriptomes, on the other hand, enable us to study the morphological heterogeneity of gene expression. Our study provides a useful resource for exploring the spatial features and temporal dynamics of the wide range of cell types, genes, and pathways of *G. hybrida*’s capitulum.

In our study, we focused most on the spatial expression of MADS-box genes, which play vital roles in the development of reproductive structures, such as inflorescence and flower organs. We observed four obvious spatial expression patterns of these genes and quantified the patterns by calculating the spatial correlation. Some of the MADS-box genes analyzed in our study have been studied before, and our analyses were consistent with and extended existing information on their specific expression dynamics and patterns. Additionally, numerous unstudied MADS-box genes with highly specific expression patterns were identified, the exact biological functions of which are still to be revealed. The spatial analysis provides useful clues for inferring their biological functions, as exemplified for the so far unstudied MADS-box genes. *GAGL12,* the ortholog gene of *AGL12* in *A. thaliana*, was found to be specifically expressed in supposed phloem cells. We also found several indications that this MADS-domain protein in complex with specific other MADS proteins is important for vascular tissue development in the floret-differentiation stages.

## Materials and Methods

### Plant materials

A double haploid cultivar, *G. hybrida* SH6, was chosen for all experiments in this study to avoid the complexity in sequencing and assembly of the highly heterozygous genomes of ordinary commercial cultivars. Here, we used the previously published *G. hybrida* SH6 genome(Wen *et al*., 2026) as the reference genome. All samples were collected from Yunnan Academy of Agricultural Sciences in Kunming, Yunnan province, China.

Based on previous studies(Zhang *et al*., 2021; Zhao *et al*., 2020), the capitula with the quickly expanding FMs, which are going through a quick floret-differentiation, were chosen for sequencing to generate a single-cell-level and spatial transcriptomic atlas. For cultivar SH6, the diameters of the capitula at these stages range from around 2 mm to 1 cm, and we selected three successive developmental stages in this range for further experiments, which correspond to the floret-initiation and differentiation phases of capitulum patterning.

### Single-cell transcriptome sequencing

The samples collected for scRNA sequencing were measured immediately after gathering. For single-cell RNA sequencing, we chose 10x Genomics scRNA-seq 3’ v3 Gene Expression as the basic platform. First, we removed all bracts from all three samples, followed by protoplast culturing. Enzyme solution, elution buffer, and wash buffer were prepared. Every capitulum was cut into thin slices with a blade, and the tissue slices were cleaned twice using the wash buffer. The tissue slices were placed in a centrifuge tube containing enzyme solution, and the centrifuge tube was incubated in a shaking table at 30°C, 75rpm. During incubation, the state of protoplasts (including morphology, quantity, size, number of fragments, and cell viability) was observed by microscope to determine the enzymatic hydrolysis time. After enzymatic hydrolysis, the centrifuge tube was briefly shaken to release the protoplasts completely. The enzyme solution was transferred and filtered into a new 15ml centrifuge tube using a 40μm cell sieve. 2ml wash buffer was added to the remaining tissue and shaken well, and the wash buffer was filtered into the tube using a 40μm cell sieve. The tube was centrifuged at 150g for 5 minutes. Then, the supernatant was discarded, 2ml wash buffer was added with a wide muzzle, and the protoplasts were gently reinserted. The tube was centrifuged at 150g for 3 minutes twice. After discarding the supernatant, an appropriate amount of wash buffer was added to the tube and mixed gently. 5μl protoplast suspension was mixed with 5μl 0.4% Trypan Blue dye. Cell concentration and cell activity were detected on a cell counting plate. Wash buffer was used to adjust the protoplast suspension concentration to 1,000∼2,000 cells/μl. The cells placed on the ice were used to build a 10x single-cell library within 30 minutes.

### Single-cell transcriptome analysis

10x paired-end read files (in FASTQ format) were generated for three samples, respectively. Using 10x Genomics Cell Ranger (ver. 7.1.0), the raw sequencing data were then mapped to the genome of *G. hybrida,* and single-cell gene expression profiles were generated. We chose Scanpy(Wolf *et al*., 2018) (ver. 1.9.6) as the basic method for further analyses. The expression profiles of the three samples were analyzed together. Based on the initial analysis, cells with a total expression count of less than 2,000 were considered of low sequencing quality and were filtered out (Sup. Fig. S1). After normalizing the expression profile to a total count of 100,000 for each cell, the detection of highly variable genes (default setting), and principal component analysis (PCA) (svd_solver=’arpack’, use_highly_variable=True), we calculated the neighbor distances of cells (method=’gauss’, n_neighbors=70). Following the advice given by the developers of Scanpy, we obtained the diffusion map of cells by diffmap(Coifman *et al*., 2005; Haghverdi *et al*., 2015) in Scanpy, and then we recalculated the neighbor distances based on the diffusion map. The recalculated neighbor distances were then used for subsequent analyses. We applied UMAP(Becht *et al*., 2019) (ver. 0.5.3, default parameters) as a dimensionality reduction method and leidenalg (Traag et al., 2019) (ver. 0.9.1, resolution=2) for cell clustering. After this, for each cluster, we used the logistic regression method(Ntranos *et al*., 2019) (sc.tl.rank_genes_groups (adata, groupby=‘leiden’, method=’logreg’)) provided in Scanpy to detect differentially expressed genes (DEGs) in each cluster. These DEGs were taken as putative marker genes of each cluster.

### Spatial transcriptome sequencing and optical image capture

We used Stereo-seq(Chen *et al*., 2022) to measure high-resolution spatial expression profiles of the capitulum. After collecting the capitula from plants, we first removed the bracts to reduce possible noise. Then, the samples were embedded in the Tissue-Tek® O.C.T. Compound and frozen at -70 □ immediately. Sections with 10μm thickness were dissected from the radial symmetry axis of the capitula and transferred to the Stereo-seq microarrays. The chips were fixed in methanol at -20°C for 30 minutes, followed by nucleic acid staining (CFW and Invitrogen Qubit ssDNA HS Reagent) and imaging (Ti-7 Nikon Eclipse microscope). Imaging was performed using the FITC channel at 10× magnification, capturing the entire 10×10mm chip. To release mRNAs from our tissue, each chip was treated with 100μl of 0.01N HCl (containing Permeabilization Enzyme) for permeabilization, followed by reverse transcription (SuperScript II, Invitrogen). After PCR amplification, purification was performed using VAHTSTM DNA Clean Beads (VAZYME). In the end, the chips were sequenced on the MGI DNBSEQ Tx sequencer.

### Spatial transcriptome preprocessing

After we obtained the Stereo-seq data, we first aligned the reads to the UMI library to retrieve the spatial information of each read and then to *G. hybrida’s* genome to generate expression profiles. After combining the spatial transcriptomes and optical images (obtained after ssDNA staining), we manually segmented 19 samples and obtained 19 gene expression matrices from 4 Stereo-seq results. The low-quality chip SP2 were abandoned in downstream analysis. On a 10×10 mm Stereo-seq chip, 26,460×26,460 barcode pools are arranged tightly to capture mRNA sequences. The distance between each pool is about 0.7 microns, which is much smaller than the average cell diameter. In our study, we need to compress the data first to study the spatial transcriptome at the cell or sub-tissue level. Considering the highly different cell sizes in our tissue, we chose to pool all cells in a 50×50 window. This compressed basic unit will be called a pseudo-cell.

### Integration of scRNA-seq and spatial transcriptome data

To integrate the single-cell and spatial transcriptomes, we adopted cell2location(Kleshchevnikov *et al*., 2022) (ver. 0.1.3), a neural network-based method, to infer the distribution of cell types in the single-cell data. Each cell type inferred from the single-cell transcriptome was represented by a typical gene expression profile or “signature”. These signatures were used to infer the cell type distribution in spatial transcriptomes, based on which cell types were then clustered (Leiden clustering) to study tissue-level heterogeneity. The spatial transcriptomes were then integrated with matched single-cell transcriptomes, respectively, generating putative spatial cell-type distributions.

### Inference of single-cell gene regulatory networks

We took advantage of a hybrid approach to discover TF genes in the *G. hybrida* genome. Protein sequences of well-classified and well-studied TFs, such as genes in the WRKY, GRF, and MIKC-MADS families in *Helianthus annuus* (sunflower) and *Chrysanthemum seticuspe*, two other species in Asteraceae, and *A. thaliana*, were obtained from PlantTFDB. Then we used BlastP(Camacho *et al*., 2009) (ver. 2.14.1+) to map the obtained TF sequences to the *G. hybrida* proteome for the identification of TFs. As an addition, we also applied InterProScan(Paysan-Lafosse *et al*., 2023) (ver. 5.64-96.0) for overall annotation and selected genes annotated as TFs. Finally, 3,790 proteins were recognized as TFs, classified into TF gene families, and used for GRN inference.

After obtaining the TF gene list, we used pySCENIC(Aibar *et al*., 2017; Van de Sande *et al*., 2020) (ver. 0.12.1) as the basic pipeline to infer gene regulatory networks. We chose the single-cell data as input for the inference of GRNs, as these are less sparse than the spatial transcriptomes. The GRNs generated for the three single-cell samples consist of predicted regulatory relationships that contain a TF, a target gene, and a variable importance measure (VIM), which indicates how important variation in the TF is for predicting the expression level of the target gene. To focus on the most confidently predicted regulatory relationships, we chose a VIM threshold of 1.0 for each of the three GRNs. Three filtered GRNs were reconstructed with the regulatory relationships selected from the three single-cell data, respectively (Supplementary Materials).

### Calculation of spatial correlation

To quantify the spatial expression similarity of genes in spatial transcriptomes, we adopted a convolution-based method to decrease the effects of data sparsity in spatial transcriptomes. Genes with total expression count less than 100 in each spatial transcriptome were filtered out to reduce noise. A 5× 5 pseudo-cell isotropic Gaussian kernel (sigma=3) was applied to diffuse the expression profiles from neighboring spatial domains. Next, we calculated the Pearson correlation between the resulting profiles, which we call pseudo-cell spatial correlation. We analyzed each MADS-box gene’s pseudo-cell spatial correlation with all the other MADS-box genes in all samples.

### Genome function annotation

We chose Gene Ontology (GO) and Kyoto Encyclopedia of Genes and Genomes (KEGG) for further study of gene function. GO annotation was performed by eggNOG-mapper online, and KEGG annotation by KofamKOALA with KOfam database (ver. 2024-03-25)(Aramaki *et al*., 2020). All GO enrichment analyses were performed by GOATOOLS(Klopfenstein *et al*., 2018) with the GO database goslim_plant. KEGG Mapper was chosen for the reconstruction of KEGG pathways(Kanehisa *et al*., 2022).

### Phylogenetic analysis of MADS-box genes

MADS-domain protein sequences in *A. thaliana*, including M-type, MIKC, and MIKC*-types were obtained from PlantTFDB. We chose Clustal Omega(Sievers *et al*., 2011) (ver. 1.2.4) for multiple sequence alignment (MSA) of all MADS-box protein sequences. The resulting alignment was then used to construct a maximum-likelihood tree by IQ-TREE(Minh *et al*., 2020) (ver. 2.2.2.6). The newly annotated MADS-box genes were then named by the sub-clade name, the same naming patterns of genes already studied in the same sub-clade, or the best alignment based on Blast results.

### CArG-box occurrence scanning

All motifs representing putative consensus binding-sites of MADS-domain proteins (CArG-boxes) were downloaded from JASPAR (10th release, 2024) (Rauluseviciute et al., 2024). The 3kb upstream sequences of genes in G. hybrida were extracted from the genome assembly by BEDTools (ver. 2.31.0) (Quinlan and Hall, 2010). The upstream sequences were then scanned using FIMO (Grant et al., 2011) (ver. 5.5.3) for the occurrence of the MADS binding site motifs. Sequences with FIMO q-value less than 0.1 were taken as valid binding sites.

### Inference of putative target genes of *GAGL12*

In order to infer a list of genes potentially regulated by *GAGL12*, we selected target genes in GRNs of *GAGL12* inferred by pySCENIC, the genes with significant CArG-box occurrence on the 3kb upstream, and the DEGs of phloem cells (including clusters 21, 28, and 30). The intersection of these three lists was taken as a set of putative target genes of *GAGL12*.

### Cell type transition analysis

We used moscot (Klein *et al*., 2025), an optimal-transportation-based method, to generate the cell-type transition probability matrix and to perform downstream analyses of cells in single-cell data. Cells in IM/early-FM, FM, EP, EN, PL, and P/X were selected for further analysis by TemporalProblem package in moscot.problems.time using default parameters.

### Experimental validation of protein-to-protein interaction

The yeast two-hybrid approach(Brückner *et al*., 2009; de Folter and Immink, 2011) was applied to test for protein-protein interactions between GAGL12 protein and a selection of other *G. hybrida* MADS-domain proteins. We amplified the coding sequence (CDS) of genes *GAGL12*, *GRCD2*, *GRCD6*, *GRCD7*, *GFLC6, GSQUA1*, *GSQUA3*, *GSQUA6*, *GSQUA8*, *Ger_023266, GAGL24L5*, *GAGL24L6*, *GANR1L10*, and *Ger_032437*, and inserted them into the *pGADT7* or *pGBKT7* vectors to build recombinant constructs, which were transferred into the yeast strain Y2H Gold. The *pGADT7*-T and *pGBKT7*-53 co-expression system serves as the positive control, while *pGADT7*-T and *pGBKT7*-lam are used as the negative control.

Yeast suspensions were serially diluted at gradients of 100, 10□¹, 10□², 10□³ (OD600 = 0.2, 0.02, 0.002, 0.0002), and 5 μL of each dilution was spot-inoculated onto (1) SD/-Leu/-Trp agar plates and (2) SD/-Leu/-Trp/-His/-Ade/X-α-gal agar plates. Yeast growth on these plates was recorded to detect potential protein-protein interactions.

## Acknowledgements

This work was supported by Yunnan Fundamental Research Projects (202401AW070007), Yunnan Xingdian Talents—Special Selection Project for High-level Scientific and Technological Talents and Innovation Teams-Team Specific Project (202505AS350021), Yunnan Xingdian Talents—Youth Special Project (XDYC-QNRC-2022-0731), and Major Science and Technology Project of Yunnan Provincial Department of Science and Technology (202602AE090010).

## Author Contributions

J.W., A.D.J.v.D., and H.S. conceived and supervised the study; Y.G., F.L. and C.J. designed and performed the experiments and computational analyses; Y.G., D.d.R., R.I., Y.S., P.H. and Y.C. contributed to data interpretation and methodology; Y.G. wrote the manuscript with input from all authors. All authors read and approved the final manuscript.

## Data Availability

The single-cell RNA sequencing and spatial transcriptome data generated in this study have been deposited in CNGBdb with accession number CNP0007743 (available when published). All other data supporting the findings of this study are available within the article and its supplementary materials, or from the corresponding author upon reasonable request.

## Supplementary Information

**Figure S1:** The overview of unfiltered single-cell transcriptomes.

**Figure S2:** The distribution of the gene expression counts and expressed gene number of all clusters for cells after filtering on the UMAP embedding.

**Figure S3:** The spatial expression of single-cell transcriptomics marker genes in the stereo-seq sample SP1_d.

**Figure S4:** The cell2location result of mapping the cell cluster in single-cell transcriptome S1 onto spatial transcriptome SP3_g.

**Figure S5:** The cell2location result of mapping cell clusters in single-cell transcriptome S2 onto spatial transcriptome SP3_f.

**Figure S6:** The cell2location result of mapping the cell cluster in single-cell transcriptome S3 onto the spatial transcriptome SP1_d.

**Figure S7:** Phylogenetic tree of MADS-box genes in *A. thaliana* and *G. hybrida*.

**Figure S8:** The yeast-two hybrid experiment result for testing the interaction between GAGL12 protein and the selected MADS-domain proteins.

**Figure S9:** Expressions of putative target genes of *GAGL12*.

**Table S1:** The conversion of gene locus tags in the G. hybrida genome to reference marker gene symbols in previous studies.

**Table S2:** The inferred top 200 differentially expressed genes (DEGs) in each cluster of single-cell transcriptomes. The rows with the name ending with ‘name’ are the gene names, and the rows with the name ending with ‘score’ are the scores.

**Table S3:** The conversion of locus tags of all MADS-box genes to gene symbols.

**Table S4:** The gene regulatory networks (GRNs) of all transcription factors (TFs) in three single-cell samples inferred by pyscenic.

**Table S5:** The functional annotation of putative target genes regulated by *GAGL12*.

**File S1:** The gene ontology (GO) enrichment results for the top 200 DEGs in each cluster of the single-cell transcriptome. The color indicates the GO class; the length indicates -log(p-value); the number on the left side indicates the gene number annotated to this GO term.

