## Supplementary File S1 for "An integrative single-cell and spatial transcriptomics atlas highlights candidate regulatory factors in the development of gerbera capitulum"

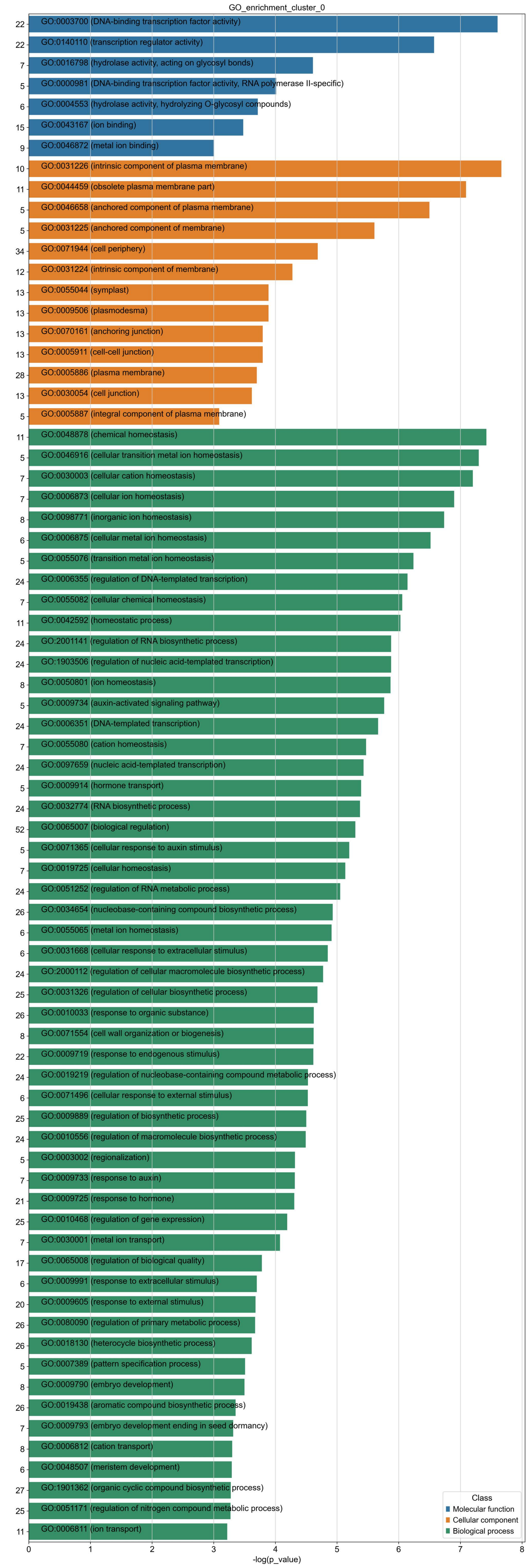
GO_enrichment_cluster_0
22 GO:0003700 (DNA-binding transcription factor activity)
22 GO:0140110 (transcription regulator activity)
7 GO:0016798 (hydrolase activity, acting on glycosyl bonds)
5 GO:0000981 (DNA-binding transcription factor activity, RNA polymerase II-specific)
6 GO:0004553 (hydrolase activity, hydrolyzing O-glycosyl compounds)
15 GO:0043167 (ion binding)
9 GO:0046872 (metal ion binding)
10 GO:0031226 (intrinsic component of plasma membrane)
11 GO:0044459 (obsolete plasma membrane part)
5 GO:0046658 (anchored component of plasma membrane)
5 GO:0031225 (anchored component of membrane)
34 GO:0071944 (cell periphery)
12 GO:0031224 (intrinsic component of membrane)
13 GO:0055044 (symplast)
13 GO:0009506 (plasmodesma)
13 GO:0070161 (anchoring junction)
13 GO:0005911 (cell-cell junction)
28 GO:0005886 (plasma membrane)
13 GO:0030054 (cell junction)
5 GO:0005887 (integral component of plasma membrane)
11 GO:0048878 (chemical homeostasis)
5 GO:0046916 (cellular transition metal ion homeostasis)
7 GO:0030003 (cellular cation homeostasis)
7 GO:0006873 (cellular ion homeostasis)
8 GO:0098771 (inorganic ion homeostasis)
6 GO:0006875 (cellular metal ion homeostasis)
5 GO:0055076 (transition metal ion homeostasis)
24 GO:0006355 (regulation of DNA-templated transcription)
7 GO:0055082 (cellular chemical homeostasis)
11 GO:0042592 (homeostatic process)
24 GO:2001141 (regulation of RNA biosynthetic process)
24 GO:1903506 (regulation of nucleic acid-templated transcription)
8 GO:0050801 (ion homeostasis)
5 GO:0009734 (auxin-activated signaling pathway)
24 GO:0006351 (DNA-templated transcription)
7 GO:0055080 (cation homeostasis)
24 GO:0097659 (nucleic acid-templated transcription)
5 GO:0009914 (hormone transport)
24 GO:0032774 (RNA biosynthetic process)
52 GO:0065007 (biological regulation)
5 GO:0071365 (cellular response to auxin stimulus)
7 GO:0019725 (cellular homeostasis)
24 GO:0051252 (regulation of RNA metabolic process)
26 GO:0034654 (nucleobase-containing compound biosynthetic process)
6 GO:0055065 (metal ion homeostasis)
6 GO:0031668 (cellular response to extracellular stimulus)
24 GO:2000112 (regulation of cellular macromolecule biosynthetic process)
25 GO:0031326 (regulation of cellular biosynthetic process)
26 GO:0010033 (response to organic substance)
8 GO:0071554 (cell wall organization or biogenesis)
22 GO:0009719 (response to endogenous stimulus)
24 GO:0019219 (regulation of nucleobase-containing compound metabolic process)
6 GO:0071496 (cellular response to external stimulus)
25 GO:0009889 (regulation of biosynthetic process)
24 GO:0010556 (regulation of macromolecule biosynthetic process)
5 GO:0003002 (regionalization)
7 GO:0009733 (response to auxin)
21 GO:0009725 (response to hormone)
25 GO:0010468 (regulation of gene expression)
7 GO:0030001 (metal ion transport)
17 GO:0065008 (regulation of biological quality)
6 GO:0009991 (response to extracellular stimulus)
20 GO:0009605 (response to external stimulus)
26 GO:0080090 (regulation of primary metabolic process)
26 GO:0018130 (heterocycle biosynthetic process)
5 GO:0007389 (pattern specification process)
8 GO:0009790 (embryo development)
26 GO:0019438 (aromatic compound biosynthetic process)
7 GO:0009793 (embryo development ending in seed dormancy)
8 GO:0006812 (cation transport)
6 GO:0048507 (meristem development)
27 GO:1901362 (organic cyclic compound biosynthetic process)
25 GO:0051171 (regulation of nitrogen compound metabolic process)
11 GO:0006811 (ion transport)
0 1 2 3 4 5 6 7 8
-log(p_value)
Class
Molecular function
Cellular component
Biological process

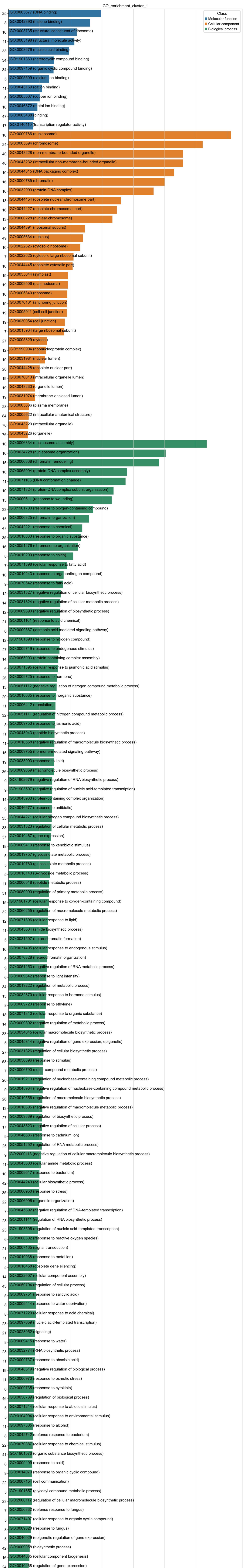
GO_enrichment_cluster_1
Class
Molecular function
Cellular component
Biological process
-log(p_value)

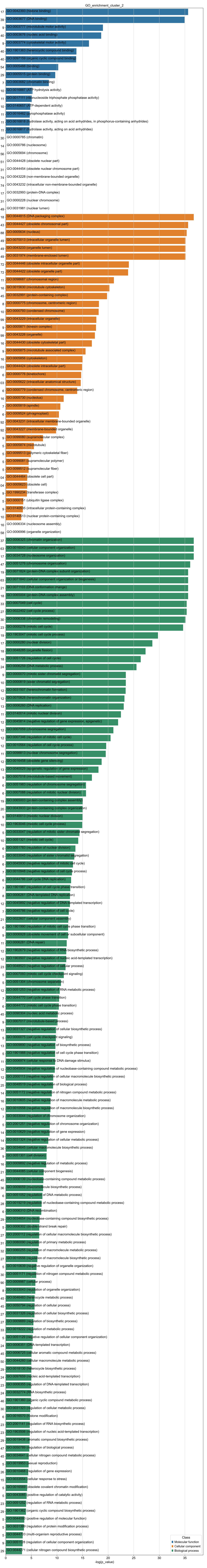
GO_enrichment_cluster_2
17 GO:0042393 (histone binding)
39 GO:0003677 (DNA binding)
9 GO:0003777 (microtubule motor activity)
40 GO:0003676 (nucleic acid binding)
9 GO:0003774 (cytoskeletal motor activity)
40 GO:1901363 (heterocyclic compound binding)
40 GO:0097159 (organic cyclic compound binding)
54 GO:0005488 (binding)
29 GO:0005515 (protein binding)
7 GO:0003682 (chromatin binding)
10 GO:0016887 (ATP hydrolysis activity)
11 GO:0017111 (ribonucleoside triphosphate phosphatase activity)
10 GO:0140657 (ATP-dependent activity)
11 GO:0016462 (pyrophosphatase activity)
11 GO:0016818 (hydrolase activity, acting on acid anhydrides, in phosphorus-containing anhydrides)
11 GO:0016817 (hydrolase activity, acting on acid anhydrides)
30 GO:0000785 (chromatin)
14 GO:0000786 (nucleosome)
56 GO:0005694 (chromosome)
51 GO:0044428 (obsolete nuclear part)
31 GO:0044454 (obsolete nuclear chromosome part)
74 GO:0043228 (non-membrane-bounded organelle)
74 GO:0043232 (intracellular non-membrane-bounded organelle)
17 GO:0032993 (protein-DNA complex)
31 GO:0000228 (nuclear chromosome)
49 GO:0031981 (nuclear lumen)
18 GO:0044815 (DNA packaging complex)
43 GO:0044427 (obsolete chromosomal part)
88 GO:0005634 (nucleus)
49 GO:0070013 (intracellular organelle lumen)
49 GO:0043233 (organelle lumen)
49 GO:0031974 (membrane-enclosed lumen)
72 GO:0044446 (obsolete intracellular organelle part)
72 GO:0044422 (obsolete organelle part)
11 GO:0098687 (chromosomal region)
16 GO:0015630 (microtubule cytoskeleton)
45 GO:0032991 (protein-containing complex)
8 GO:0000775 (chromosome, centromeric region)
10 GO:0000793 (condensed chromosome)
99 GO:0043229 (intracellular organelle)
9 GO:0005871 (kinesin complex)
99 GO:0043226 (organelle)
16 GO:0044430 (obsolete cytoskeletal part)
9 GO:0005875 (microtubule associated complex)
16 GO:0005856 (cytoskeleton)
103 GO:0044424 (obsolete intracellular part)
6 GO:0000776 (kinetochore)
103 GO:0005622 (intracellular anatomical structure)
6 GO:0000779 (condensed chromosome, centromeric region)
16 GO:0005730 (nucleolus)
7 GO:0005819 (spindle)
6 GO:0009524 (phragmoplast)
92 GO:0043231 (intracellular membrane-bounded organelle)
92 GO:0043227 (membrane-bounded organelle)
9 GO:0099080 (supramolecular complex)
5 GO:0005874 (microtubule)
5 GO:0099513 (polymeric cytoskeletal fiber)
5 GO:0099081 (supramolecular polymer)
5 GO:0099512 (supramolecular fiber)
104 GO:0044464 (obsolete cell part)
104 GO:0005623 (obsolete cell)
9 GO:1990234 (transferase complex)
5 GO:0000151 (ubiquitin ligase complex)
9 GO:0140535 (intracellular protein-containing complex)
10 GO:0140513 (nuclear protein-containing complex)
16 GO:0006334 (nucleosome assembly)
58 GO:0006996 (organelle organization)
37 GO:0006325 (chromatin organization)
63 GO:0016043 (cellular component organization)
17 GO:0034728 (nucleosome organization)
47 GO:0051276 (chromosome organization)
19 GO:0071824 (protein-DNA complex subunit organization)
63 GO:0071840 (cellular component organization or biogenesis)
21 GO:0071103 (DNA conformation change)
18 GO:0065004 (protein-DNA complex assembly)
33 GO:0007049 (cell cycle)
27 GO:0022402 (cell cycle process)
30 GO:0006338 (chromatin remodeling)
23 GO:0000278 (mitotic cell cycle)
19 GO:1903047 (mitotic cell cycle process)
17 GO:0000280 (nuclear division)
18 GO:0048285 (organelle fission)
18 GO:0051726 (regulation of cell cycle)
24 GO:0006259 (DNA metabolic process)
9 GO:0000070 (mitotic sister chromatid segregation)
10 GO:0000819 (sister chromatid segregation)
12 GO:0031507 (heterochromatin formation)
12 GO:0070828 (heterochromatin organization)
13 GO:0006260 (DNA replication)
11 GO:0140014 (mitotic nuclear division)
12 GO:0045814 (negative regulation of gene expression, epigenetic)
12 GO:0007059 (chromosome segregation)
12 GO:0007346 (regulation of mitotic cell cycle)
12 GO:0010564 (regulation of cell cycle process)
11 GO:0098813 (nuclear chromosome segregation)
12 GO:0016458 (obsolete gene silencing)
13 GO:0040029 (epigenetic regulation of gene expression)
9 GO:0007018 (microtubule-based movement)
6 GO:0051983 (regulation of chromosome segregation)
6 GO:0007088 (regulation of mitotic nuclear division)
19 GO:0065003 (protein-containing complex assembly)
20 GO:0043933 (protein-containing complex organization)
9 GO:0140013 (meiotic nuclear division)
10 GO:1903046 (meiotic cell cycle process)
5 GO:0033047 (regulation of mitotic sister chromatid segregation)
10 GO:0051321 (meiotic cell cycle)
6 GO:0051783 (regulation of nuclear division)
5 GO:0033045 (regulation of sister chromatid segregation)
6 GO:0045930 (negative regulation of mitotic cell cycle)
7 GO:0010948 (negative regulation of cell cycle process)
6 GO:0044786 (cell cycle DNA replication)
7 GO:1901987 (regulation of cell cycle phase transition)
8 GO:0006261 (DNA-templated DNA replication)
12 GO:0045892 (negative regulation of DNA-templated transcription)
7 GO:0045786 (negative regulation of cell cycle)
21 GO:0022607 (cellular component assembly)
6 GO:1901990 (regulation of mitotic cell cycle phase transition)
9 GO:0006928 (obsolete movement of cell or subcellular component)
11 GO:0006281 (DNA repair)
12 GO:1902679 (negative regulation of RNA biosynthetic process)
12 GO:1903507 (negative regulation of nucleic acid-templated transcription)
23 GO:0048523 (negative regulation of cellular process)
5 GO:0007093 (mitotic cell cycle checkpoint signaling)
5 GO:0051304 (chromosome separation)
12 GO:0051253 (negative regulation of RNA metabolic process)
7 GO:0044770 (cell cycle phase transition)
6 GO:0044772 (mitotic cell cycle phase transition)
45 GO:0090304 (nucleic acid metabolic process)
9 GO:0007017 (microtubule-based process)
13 GO:0031327 (negative regulation of cellular biosynthetic process)
5 GO:0000075 (cell cycle checkpoint signaling)
13 GO:0009890 (negative regulation of biosynthetic process)
5 GO:1901988 (negative regulation of cell cycle phase transition)
11 GO:0006974 (cellular response to DNA damage stimulus)
12 GO:0045934 (negative regulation of nucleobase-containing compound metabolic process)
12 GO:2000113 (negative regulation of cellular macromolecule biosynthetic process)
25 GO:0048519 (negative regulation of biological process)
14 GO:0051172 (negative regulation of nitrogen compound metabolic process)
16 GO:0010605 (negative regulation of macromolecule metabolic process)
12 GO:0010558 (negative regulation of macromolecule biosynthetic process)
7 GO:0033044 (regulation of chromosome organization)
5 GO:2001251 (negative regulation of chromosome organization)
14 GO:0010629 (negative regulation of gene expression)
14 GO:0031324 (negative regulation of cellular metabolic process)
36 GO:0034645 (cellular macromolecule biosynthetic process)
9 GO:0051301 (cell division)
16 GO:0009892 (negative regulation of metabolic process)
21 GO:0044085 (cellular component biogenesis)
45 GO:0006139 (nucleobase-containing compound metabolic process)
36 GO:0009059 (macromolecule biosynthetic process)
6 GO:0051052 (regulation of DNA metabolic process)
28 GO:0019219 (regulation of nucleobase-containing compound metabolic process)
6 GO:0006310 (DNA recombination)
29 GO:0034654 (nucleobase-containing compound biosynthetic process)
5 GO:0006302 (double-strand break repair)
27 GO:2000112 (regulation of cellular macromolecule biosynthetic process)
31 GO:0080090 (regulation of primary metabolic process)
32 GO:0060255 (regulation of macromolecule metabolic process)
27 GO:0010556 (regulation of macromolecule biosynthetic process)
5 GO:0010639 (negative regulation of organelle organization)
30 GO:0051171 (regulation of nitrogen compound metabolic process)
90 GO:0009987 (cellular process)
8 GO:0033043 (regulation of organelle organization)
45 GO:0046483 (heterocycle metabolic process)
45 GO:0050794 (regulation of cellular process)
27 GO:0031326 (regulation of cellular biosynthetic process)
27 GO:0009889 (regulation of biosynthetic process)
33 GO:0019222 (regulation of metabolic process)
5 GO:0051129 (negative regulation of cellular component organization)
24 GO:0006351 (DNA-templated transcription)
45 GO:0006725 (cellular aromatic compound metabolic process)
50 GO:0044260 (cellular macromolecule metabolic process)
29 GO:0018130 (heterocycle biosynthetic process)
24 GO:0097659 (nucleic acid-templated transcription)
23 GO:0006355 (regulation of DNA-templated transcription)
24 GO:0032774 (RNA biosynthetic process)
46 GO:1901360 (organic cyclic compound metabolic process)
30 GO:0031323 (regulation of cellular metabolic process)
6 GO:0016570 (histone modification)
23 GO:2001141 (regulation of RNA biosynthetic process)
23 GO:1903506 (regulation of nucleic acid-templated transcription)
29 GO:0019438 (aromatic compound biosynthetic process)
47 GO:0050789 (regulation of biological process)
45 GO:0034641 (cellular nitrogen compound metabolic process)
5 GO:0019953 (sexual reproduction)
26 GO:0010468 (regulation of gene expression)
15 GO:0033554 (cellular response to stress)
5 GO:0016569 (obsolete covalent chromatin modification)
5 GO:0043085 (positive regulation of catalytic activity)
23 GO:0051252 (regulation of RNA metabolic process)
29 GO:1901362 (organic cyclic compound biosynthetic process)
5 GO:0044093 (positive regulation of molecular function)
5 GO:0031399 (regulation of protein modification process)
5 GO:0044703 (multi-organism reproductive process)
9 GO:0051128 (regulation of cellular component organization)
29 GO:0044271 (cellular nitrogen compound biosynthetic process)
0
5
10
15
20
25
30
35
-log(p_value)
Class
Molecular function
Cellular component
Biological process

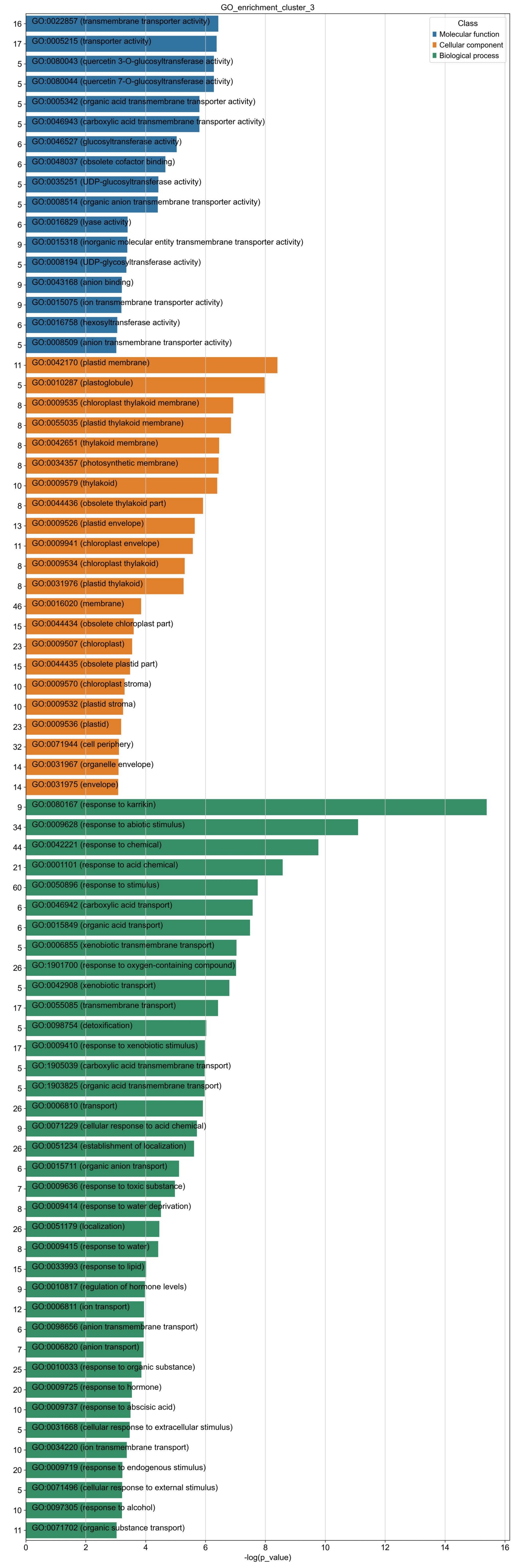

GO_enrichment_cluster_3
Class
Molecular function
Cellular component
Biological process
16 GO:0022857 (transmembrane transporter activity)
17 GO:0005215 (transporter activity)
5 GO:0080043 (quercetin 3-O-glucosyltransferase activity)
5 GO:0080044 (quercetin 7-O-glucosyltransferase activity)
5 GO:0005342 (organic acid transmembrane transporter activity)
5 GO:0046943 (carboxylic acid transmembrane transporter activity)
6 GO:0046527 (glucosyltransferase activity)
6 GO:0048037 (obsolete cofactor binding)
5 GO:0035251 (UDP-glucosyltransferase activity)
5 GO:0008514 (organic anion transmembrane transporter activity)
6 GO:0016829 (lyase activity)
9 GO:0015318 (inorganic molecular entity transmembrane transporter activity)
5 GO:0008194 (UDP-glycosyltransferase activity)
9 GO:0043168 (anion binding)
9 GO:0015075 (ion transmembrane transporter activity)
6 GO:0016758 (hexosyltransferase activity)
5 GO:0008509 (anion transmembrane transporter activity)
11 GO:0042170 (plastid membrane)
5 GO:0010287 (plastoglobule)
8 GO:0009535 (chloroplast thylakoid membrane)
8 GO:0055035 (plastid thylakoid membrane)
8 GO:0042651 (thylakoid membrane)
8 GO:0034357 (photosynthetic membrane)
10 GO:0009579 (thylakoid)
8 GO:0044436 (obsolete thylakoid part)
13 GO:0009526 (plastid envelope)
11 GO:0009941 (chloroplast envelope)
8 GO:0009534 (chloroplast thylakoid)
8 GO:0031976 (plastid thylakoid)
46 GO:0016020 (membrane)
15 GO:0044434 (obsolete chloroplast part)
23 GO:0009507 (chloroplast)
15 GO:0044435 (obsolete plastid part)
10 GO:0009570 (chloroplast stroma)
10 GO:0009532 (plastid stroma)
23 GO:0009536 (plastid)
32 GO:0071944 (cell periphery)
14 GO:0031967 (organelle envelope)
14 GO:0031975 (envelope)
9 GO:0080167 (response to karrikin)
34 GO:0009628 (response to abiotic stimulus)
44 GO:0042221 (response to chemical)
21 GO:0001101 (response to acid chemical)
60 GO:0050896 (response to stimulus)
6 GO:0046942 (carboxylic acid transport)
6 GO:0015849 (organic acid transport)
5 GO:0006855 (xenobiotic transmembrane transport)
26 GO:1901700 (response to oxygen-containing compound)
5 GO:0042908 (xenobiotic transport)
17 GO:0055085 (transmembrane transport)
5 GO:0098754 (detoxification)
17 GO:0009410 (response to xenobiotic stimulus)
5 GO:1905039 (carboxylic acid transmembrane transport)
5 GO:1903825 (organic acid transmembrane transport)
26 GO:0006810 (transport)
9 GO:0071229 (cellular response to acid chemical)
26 GO:0051234 (establishment of localization)
6 GO:0015711 (organic anion transport)
7 GO:0009636 (response to toxic substance)
8 GO:0009414 (response to water deprivation)
26 GO:0051179 (localization)
8 GO:0009415 (response to water)
15 GO:0033993 (response to lipid)
9 GO:0010817 (regulation of hormone levels)
12 GO:0006811 (ion transport)
6 GO:0098656 (anion transmembrane transport)
7 GO:0006820 (anion transport)
25 GO:0010033 (response to organic substance)
20 GO:0009725 (response to hormone)
10 GO:0009737 (response to abscisic acid)
5 GO:0031668 (cellular response to extracellular stimulus)
10 GO:0034220 (ion transmembrane transport)
20 GO:0009719 (response to endogenous stimulus)
5 GO:0071496 (cellular response to external stimulus)
10 GO:0097305 (response to alcohol)
11 GO:0071702 (organic substance transport)
0 2 4 6 8 10 12 14 16
-log(p_value)

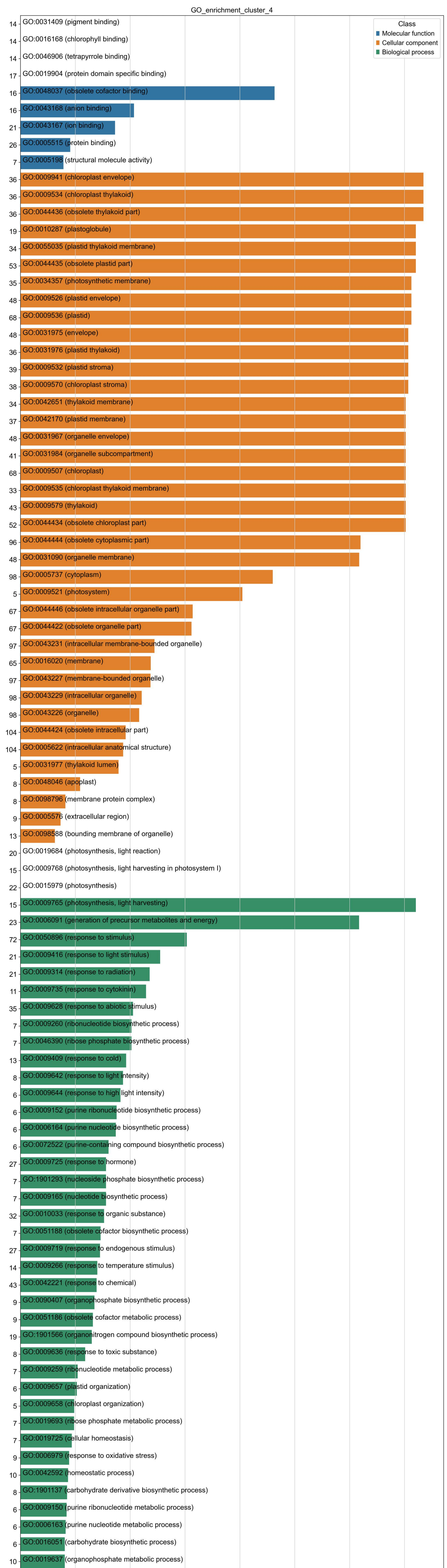
GO_enrichment_cluster_4
Class
Molecular function
Cellular component
Biological process
14 GO:0031409 (pigment binding)
14 GO:0016168 (chlorophyll binding)
14 GO:0046906 (tetrapyrrole binding)
17 GO:0019904 (protein domain specific binding)
16 GO:0048037 (obsolete cofactor binding)
16 GO:0043168 (anion binding)
21 GO:0043167 (ion binding)
26 GO:0005515 (protein binding)
7 GO:0005198 (structural molecule activity)
36 GO:0009941 (chloroplast envelope)
36 GO:0009534 (chloroplast thylakoid)
36 GO:0044436 (obsolete thylakoid part)
19 GO:0010287 (plastoglobule)
34 GO:0055035 (plastid thylakoid membrane)
53 GO:0044435 (obsolete plastid part)
35 GO:0034357 (photosynthetic membrane)
48 GO:0009526 (plastid envelope)
68 GO:0009536 (plastid)
48 GO:0031975 (envelope)
36 GO:0031976 (plastid thylakoid)
39 GO:0009532 (plastid stroma)
38 GO:0009570 (chloroplast stroma)
34 GO:0042651 (thylakoid membrane)
37 GO:0042170 (plastid membrane)
48 GO:0031967 (organelle envelope)
41 GO:0031984 (organelle subcompartment)
68 GO:0009507 (chloroplast)
33 GO:0009535 (chloroplast thylakoid membrane)
43 GO:0009579 (thylakoid)
52 GO:0044434 (obsolete chloroplast part)
96 GO:0044444 (obsolete cytoplasmic part)
48 GO:0031090 (organelle membrane)
98 GO:0005737 (cytoplasm)
5 GO:0009521 (photosystem)
67 GO:0044446 (obsolete intracellular organelle part)
67 GO:0044422 (obsolete organelle part)
97 GO:0043231 (intracellular membrane-bounded organelle)
65 GO:0016020 (membrane)
97 GO:0043227 (membrane-bounded organelle)
98 GO:0043229 (intracellular organelle)
98 GO:0043226 (organelle)
104 GO:0044424 (obsolete intracellular part)
104 GO:0005622 (intracellular anatomical structure)
5 GO:0031977 (thylakoid lumen)
8 GO:0048046 (apoplast)
8 GO:0098796 (membrane protein complex)
9 GO:0005576 (extracellular region)
13 GO:0098588 (bounding membrane of organelle)
20 GO:0019684 (photosynthesis, light reaction)
15 GO:0009768 (photosynthesis, light harvesting in photosystem I)
22 GO:0015979 (photosynthesis)
15 GO:0009765 (photosynthesis, light harvesting)
23 GO:0006091 (generation of precursor metabolites and energy)
72 GO:0050896 (response to stimulus)
21 GO:0009416 (response to light stimulus)
21 GO:0009314 (response to radiation)
11 GO:0009735 (response to cytokinin)
35 GO:0009628 (response to abiotic stimulus)
7 GO:0009260 (ribonucleotide biosynthetic process)
7 GO:0046390 (ribose phosphate biosynthetic process)
13 GO:0009409 (response to cold)
8 GO:0009642 (response to light intensity)
6 GO:0009644 (response to high light intensity)
6 GO:0009152 (purine ribonucleotide biosynthetic process)
6 GO:0006164 (purine nucleotide biosynthetic process)
6 GO:0072522 (purine-containing compound biosynthetic process)
27 GO:0009725 (response to hormone)
7 GO:1901293 (nucleoside phosphate biosynthetic process)
7 GO:0009165 (nucleotide biosynthetic process)
32 GO:0010033 (response to organic substance)
7 GO:0051188 (obsolete cofactor biosynthetic process)
27 GO:0009719 (response to endogenous stimulus)
14 GO:0009266 (response to temperature stimulus)
43 GO:0042221 (response to chemical)
9 GO:0090407 (organophosphate biosynthetic process)
9 GO:0051186 (obsolete cofactor metabolic process)
19 GO:1901566 (organonitrogen compound biosynthetic process)
8 GO:0009636 (response to toxic substance)
7 GO:0009259 (ribonucleotide metabolic process)
6 GO:0009657 (plastid organization)
5 GO:0009658 (chloroplast organization)
7 GO:0019693 (ribose phosphate metabolic process)
7 GO:0019725 (cellular homeostasis)
9 GO:0006979 (response to oxidative stress)
10 GO:0042592 (homeostatic process)
8 GO:1901137 (carbohydrate derivative biosynthetic process)
6 GO:0009150 (purine ribonucleotide metabolic process)
6 GO:0006163 (purine nucleotide metabolic process)
6 GO:0016051 (carbohydrate biosynthetic process)
10 GO:0019637 (organophosphate metabolic process)
11 GO:0044283 (small molecule biosynthetic process)
5 GO:0022900 (electron transport chain)
5 GO:0009199 (ribonucleoside triphosphate metabolic process)
5 GO:0009141 (nucleoside triphosphate metabolic process)
7 GO:0009117 (nucleotide metabolic process)
7 GO:0006753 (nucleoside phosphate metabolic process)
15 GO:0009410 (response to xenobiotic stimulus)
6 GO:0072521 (purine-containing compound metabolic process)
5 GO:1901657 (glycosyl compound metabolic process)
8 GO:0048878 (chemical homeostasis)
5 GO:0009743 (response to carbohydrate)
11 GO:1901135 (carbohydrate derivative metabolic process)
5 GO:0055082 (cellular chemical homeostasis)
0
5
10
15
20
25
30
35
-log(p_value)

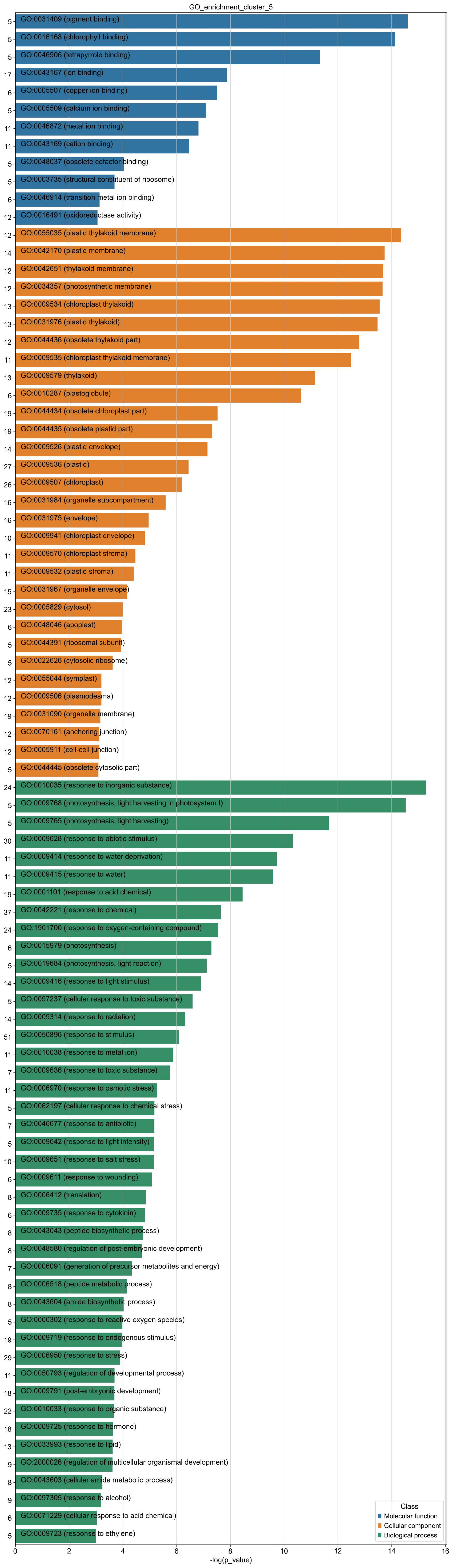

GO_enrichment_cluster_5
GO:0031409 (pigment binding)
GO:0016168 (chlorophyll binding)
GO:0046906 (tetrapyrrole binding)
GO:0043167 (ion binding)
GO:0005507 (copper ion binding)
GO:0005509 (calcium ion binding)
GO:0046872 (metal ion binding)
GO:0043169 (cation binding)
GO:0048037 (obsolete cofactor binding)
GO:0003735 (structural constituent of ribosome)
GO:0046914 (transition metal ion binding)
GO:0016491 (oxidoreductase activity)
GO:0055035 (plastid thylakoid membrane)
GO:0042170 (plastid membrane)
GO:0042651 (thylakoid membrane)
GO:0034357 (photosynthetic membrane)
GO:0009534 (chloroplast thylakoid)
GO:0031976 (plastid thylakoid)
GO:0044436 (obsolete thylakoid part)
GO:0009535 (chloroplast thylakoid membrane)
GO:0009579 (thylakoid)
GO:0010287 (plastoglobule)
GO:0044434 (obsolete chloroplast part)
GO:0044435 (obsolete plastid part)
GO:0009526 (plastid envelope)
GO:0009536 (plastid)
GO:0009507 (chloroplast)
GO:0031984 (organelle subcompartment)
GO:0031975 (envelope)
GO:0009941 (chloroplast envelope)
GO:0009570 (chloroplast stroma)
GO:0009532 (plastid stroma)
GO:0031967 (organelle envelope)
GO:0005829 (cytosol)
GO:0048046 (apoplast)
GO:0044391 (ribosomal subunit)
GO:0022626 (cytosolic ribosome)
GO:0055044 (symplast)
GO:0009506 (plasmodesma)
GO:0031090 (organelle membrane)
GO:0070161 (anchoring junction)
GO:0005911 (cell-cell junction)
GO:0044445 (obsolete cytosolic part)
GO:0010035 (response to inorganic substance)
GO:0009768 (photosynthesis, light harvesting in photosystem I)
GO:0009765 (photosynthesis, light harvesting)
GO:0009628 (response to abiotic stimulus)
GO:0009414 (response to water deprivation)
GO:0009415 (response to water)
GO:0001101 (response to acid chemical)
GO:0042221 (response to chemical)
GO:1901700 (response to oxygen-containing compound)
GO:0015979 (photosynthesis)
GO:0019684 (photosynthesis, light reaction)
GO:0009416 (response to light stimulus)
GO:0097237 (cellular response to toxic substance)
GO:0009314 (response to radiation)
GO:0050896 (response to stimulus)
GO:0010038 (response to metal ion)
GO:0009636 (response to toxic substance)
GO:0006970 (response to osmotic stress)
GO:0062197 (cellular response to chemical stress)
GO:0046677 (response to antibiotic)
GO:0009642 (response to light intensity)
GO:0009651 (response to salt stress)
GO:0009611 (response to wounding)
GO:0006412 (translation)
GO:0009735 (response to cytokinin)
GO:0043043 (peptide biosynthetic process)
GO:0048580 (regulation of post-embryonic development)
GO:0006091 (generation of precursor metabolites and energy)
GO:0006518 (peptide metabolic process)
GO:0043604 (amide biosynthetic process)
GO:0000302 (response to reactive oxygen species)
GO:0009719 (response to endogenous stimulus)
GO:0006950 (response to stress)
GO:0050793 (regulation of developmental process)
GO:0009791 (post-embryonic development)
GO:0010033 (response to organic substance)
GO:0009725 (response to hormone)
GO:0033993 (response to lipid)
GO:2000026 (regulation of multicellular organismal development)
GO:0043603 (cellular amide metabolic process)
GO:0097305 (response to alcohol)
GO:0071229 (cellular response to acid chemical)
GO:0009723 (response to ethylene)
5
5
5
17
6
5
11
11
5
5
6
12
12
14
12
12
13
13
12
11
13
6
19
19
14
27
26
16
16
10
11
11
15
23
6
5
5
12
12
19
12
12
5
24
5
5
30
11
11
19
37
24
6
5
14
5
14
51
11
7
11
5
7
5
10
6
8
6
8
8
7
8
8
5
19
29
11
18
22
18
13
9
8
9
6
5
0
2
4
6
8
10
12
14
16
-log(p_value)
Class
Molecular function
Cellular component
Biological process

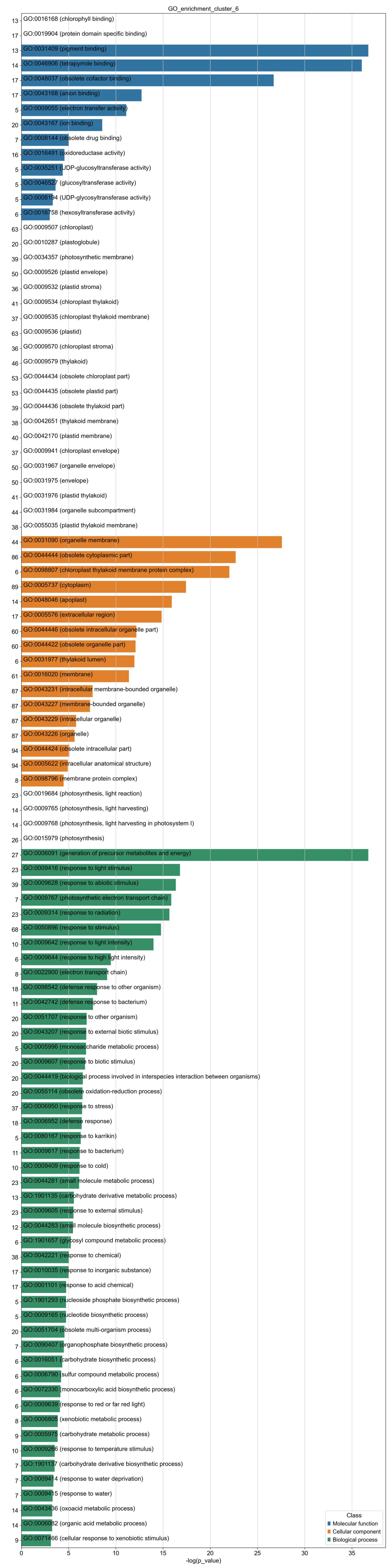
GO_enrichment_cluster_6
13 GO:0016168 (chlorophyll binding)
17 GO:0019904 (protein domain specific binding)
13 GO:0031409 (pigment binding)
14 GO:0046906 (tetrapyrrole binding)
17 GO:0048037 (obsolete cofactor binding)
17 GO:0043168 (anion binding)
5 GO:0009055 (electron transfer activity)
20 GO:0043167 (ion binding)
7 GO:0008144 (obsolete drug binding)
16 GO:0016491 (oxidoreductase activity)
5 GO:0035251 (UDP-glucosyltransferase activity)
5 GO:0046527 (glucosyltransferase activity)
5 GO:0008194 (UDP-glycosyltransferase activity)
6 GO:0016758 (hexosyltransferase activity)
63 GO:0009507 (chloroplast)
20 GO:0010287 (plastoglobule)
39 GO:0034357 (photosynthetic membrane)
50 GO:0009526 (plastid envelope)
36 GO:0009532 (plastid stroma)
41 GO:0009534 (chloroplast thylakoid)
37 GO:0009535 (chloroplast thylakoid membrane)
63 GO:0009536 (plastid)
36 GO:0009570 (chloroplast stroma)
46 GO:0009579 (thylakoid)
53 GO:0044434 (obsolete chloroplast part)
53 GO:0044435 (obsolete plastid part)
39 GO:0044436 (obsolete thylakoid part)
38 GO:0042651 (thylakoid membrane)
40 GO:0042170 (plastid membrane)
37 GO:0009941 (chloroplast envelope)
50 GO:0031967 (organelle envelope)
50 GO:0031975 (envelope)
41 GO:0031976 (plastid thylakoid)
44 GO:0031984 (organelle subcompartment)
38 GO:0055035 (plastid thylakoid membrane)
44 GO:0031090 (organelle membrane)
86 GO:0044444 (obsolete cytoplasmic part)
6 GO:0098807 (chloroplast thylakoid membrane protein complex)
89 GO:0005737 (cytoplasm)
14 GO:0048046 (apoplast)
17 GO:0005576 (extracellular region)
60 GO:0044446 (obsolete intracellular organelle part)
60 GO:0044422 (obsolete organelle part)
6 GO:0031977 (thylakoid lumen)
61 GO:0016020 (membrane)
87 GO:0043231 (intracellular membrane-bounded organelle)
87 GO:0043227 (membrane-bounded organelle)
87 GO:0043229 (intracellular organelle)
87 GO:0043226 (organelle)
94 GO:0044424 (obsolete intracellular part)
94 GO:0005622 (intracellular anatomical structure)
8 GO:0098796 (membrane protein complex)
23 GO:0019684 (photosynthesis, light reaction)
14 GO:0009765 (photosynthesis, light harvesting)
14 GO:0009768 (photosynthesis, light harvesting in photosystem I)
26 GO:0015979 (photosynthesis)
27 GO:0006091 (generation of precursor metabolites and energy)
23 GO:0009416 (response to light stimulus)
39 GO:0009628 (response to abiotic stimulus)
7 GO:0009767 (photosynthetic electron transport chain)
23 GO:0009314 (response to radiation)
68 GO:0050896 (response to stimulus)
10 GO:0009642 (response to light intensity)
6 GO:0009644 (response to high light intensity)
8 GO:0022900 (electron transport chain)
18 GO:0098542 (defense response to other organism)
11 GO:0042742 (defense response to bacterium)
20 GO:0051707 (response to other organism)
20 GO:0043207 (response to external biotic stimulus)
5 GO:0005996 (monosaccharide metabolic process)
20 GO:0009607 (response to biotic stimulus)
20 GO:0044419 (biological process involved in interspecies interaction between organisms)
20 GO:0055114 (obsolete oxidation-reduction process)
37 GO:0006950 (response to stress)
18 GO:0006952 (defense response)
5 GO:0080167 (response to karrikin)
11 GO:0009617 (response to bacterium)
10 GO:0009409 (response to cold)
23 GO:0044281 (small molecule metabolic process)
13 GO:1901135 (carbohydrate derivative metabolic process)
23 GO:0009605 (response to external stimulus)
12 GO:0044283 (small molecule biosynthetic process)
6 GO:1901657 (glycosyl compound metabolic process)
38 GO:0042221 (response to chemical)
17 GO:0010035 (response to inorganic substance)
17 GO:0001101 (response to acid chemical)
5 GO:1901293 (nucleoside phosphate biosynthetic process)
5 GO:0009165 (nucleotide biosynthetic process)
20 GO:0051704 (obsolete multi-organism process)
7 GO:0090407 (organophosphate biosynthetic process)
6 GO:0016051 (carbohydrate biosynthetic process)
6 GO:0006790 (sulfur compound metabolic process)
6 GO:0072330 (monocarboxylic acid biosynthetic process)
6 GO:0009639 (response to red or far red light)
8 GO:0006805 (xenobiotic metabolic process)
9 GO:0005975 (carbohydrate metabolic process)
10 GO:0009266 (response to temperature stimulus)
7 GO:1901137 (carbohydrate derivative biosynthetic process)
7 GO:0009414 (response to water deprivation)
7 GO:0009415 (response to water)
14 GO:0043436 (oxoacid metabolic process)
14 GO:0006082 (organic acid metabolic process)
9 GO:0071466 (cellular response to xenobiotic stimulus)
0
5
10
15
20
25
30
35
-log(p_value)
Class
Molecular function
Cellular component
Biological process

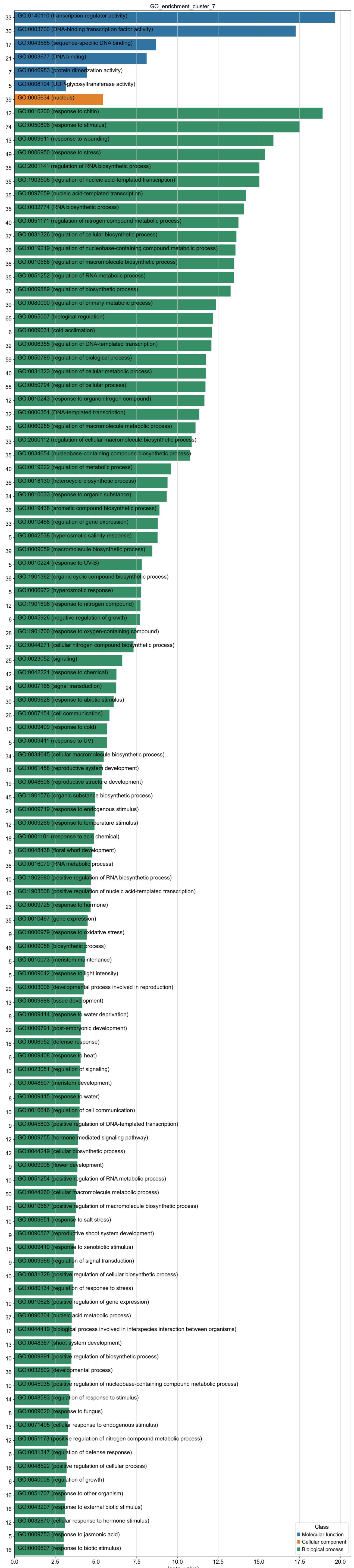
GO_enrichment_cluster_7
33 GO:0140110 (transcription regulator activity)
30 GO:0003700 (DNA-binding transcription factor activity)
17 GO:0043565 (sequence-specific DNA binding)
21 GO:0003677 (DNA binding)
7 GO:0046983 (protein dimerization activity)
5 GO:0008194 (UDP-glycosyltransferase activity)
39 GO:0005634 (nucleus)
12 GO:0010200 (response to chitin)
74 GO:0050896 (response to stimulus)
13 GO:0009611 (response to wounding)
49 GO:0006950 (response to stress)
35 GO:2001141 (regulation of RNA biosynthetic process)
35 GO:1903506 (regulation of nucleic acid-templated transcription)
35 GO:0097659 (nucleic acid-templated transcription)
35 GO:0032774 (RNA biosynthetic process)
40 GO:0051171 (regulation of nitrogen compound metabolic process)
37 GO:0031326 (regulation of cellular biosynthetic process)
36 GO:0019219 (regulation of nucleobase-containing compound metabolic process)
36 GO:0010556 (regulation of macromolecule biosynthetic process)
35 GO:0051252 (regulation of RNA metabolic process)
37 GO:0009889 (regulation of biosynthetic process)
39 GO:0080090 (regulation of primary metabolic process)
65 GO:0065007 (biological regulation)
6 GO:0009631 (cold acclimation)
32 GO:0006355 (regulation of DNA-templated transcription)
59 GO:0050789 (regulation of biological process)
40 GO:0031323 (regulation of cellular metabolic process)
55 GO:0050794 (regulation of cellular process)
12 GO:0010243 (response to organonitrogen compound)
32 GO:0006351 (DNA-templated transcription)
39 GO:0060255 (regulation of macromolecule metabolic process)
33 GO:2000112 (regulation of cellular macromolecule biosynthetic process)
35 GO:0034654 (nucleobase-containing compound biosynthetic process)
40 GO:0019222 (regulation of metabolic process)
36 GO:0018130 (heterocycle biosynthetic process)
34 GO:0010033 (response to organic substance)
36 GO:0019438 (aromatic compound biosynthetic process)
33 GO:0010468 (regulation of gene expression)
5 GO:0042538 (hyperosmotic salinity response)
39 GO:0009059 (macromolecule biosynthetic process)
5 GO:0010224 (response to UV-B)
36 GO:1901362 (organic cyclic compound biosynthetic process)
5 GO:0006972 (hyperosmotic response)
12 GO:1901698 (response to nitrogen compound)
6 GO:0045926 (negative regulation of growth)
28 GO:1901700 (response to oxygen-containing compound)
37 GO:0044271 (cellular nitrogen compound biosynthetic process)
25 GO:0023052 (signaling)
42 GO:0042221 (response to chemical)
24 GO:0007165 (signal transduction)
30 GO:0009628 (response to abiotic stimulus)
26 GO:0007154 (cell communication)
10 GO:0009409 (response to cold)
5 GO:0009411 (response to UV)
34 GO:0034645 (cellular macromolecule biosynthetic process)
19 GO:0061458 (reproductive system development)
19 GO:0048608 (reproductive structure development)
45 GO:1901576 (organic substance biosynthetic process)
24 GO:0009719 (response to endogenous stimulus)
12 GO:0009266 (response to temperature stimulus)
18 GO:0001101 (response to acid chemical)
6 GO:0048438 (floral whorl development)
36 GO:0016070 (RNA metabolic process)
10 GO:1902680 (positive regulation of RNA biosynthetic process)
10 GO:1903508 (positive regulation of nucleic acid-templated transcription)
23 GO:0009725 (response to hormone)
35 GO:0010467 (gene expression)
9 GO:0006979 (response to oxidative stress)
46 GO:0009058 (biosynthetic process)
5 GO:0010073 (meristem maintenance)
5 GO:0009642 (response to light intensity)
20 GO:0003006 (developmental process involved in reproduction)
13 GO:0009888 (tissue development)
8 GO:0009414 (response to water deprivation)
22 GO:0009791 (post-embryonic development)
16 GO:0006952 (defense response)
6 GO:0009408 (response to heat)
10 GO:0023051 (regulation of signaling)
7 GO:0048507 (meristem development)
8 GO:0009415 (response to water)
10 GO:0010646 (regulation of cell communication)
9 GO:0045893 (positive regulation of DNA-templated transcription)
12 GO:0009755 (hormone-mediated signaling pathway)
42 GO:0044249 (cellular biosynthetic process)
9 GO:0009908 (flower development)
10 GO:0051254 (positive regulation of RNA metabolic process)
50 GO:0044260 (cellular macromolecule metabolic process)
10 GO:0010557 (positive regulation of macromolecule biosynthetic process)
10 GO:0009651 (response to salt stress)
9 GO:0090567 (reproductive shoot system development)
15 GO:0009410 (response to xenobiotic stimulus)
9 GO:0009966 (regulation of signal transduction)
10 GO:0031328 (positive regulation of cellular biosynthetic process)
8 GO:0080134 (regulation of response to stress)
10 GO:0010628 (positive regulation of gene expression)
37 GO:0090304 (nucleic acid metabolic process)
17 GO:0044419 (biological process involved in interspecies interaction between organisms)
13 GO:0048367 (shoot system development)
10 GO:0009891 (positive regulation of biosynthetic process)
36 GO:0032502 (developmental process)
10 GO:0045935 (positive regulation of nucleobase-containing compound metabolic process)
14 GO:0048583 (regulation of response to stimulus)
8 GO:0009620 (response to fungus)
13 GO:0071495 (cellular response to endogenous stimulus)
12 GO:0051173 (positive regulation of nitrogen compound metabolic process)
6 GO:0031347 (regulation of defense response)
16 GO:0048522 (positive regulation of cellular process)
6 GO:0040008 (regulation of growth)
16 GO:0051707 (response to other organism)
16 GO:0043207 (response to external biotic stimulus)
12 GO:0032870 (cellular response to hormone stimulus)
5 GO:0009753 (response to jasmonic acid)
16 GO:0009607 (response to biotic stimulus)
0.0
2.5
5.0
7.5
10.0
12.5
15.0
17.5
20.0
-log(p_value)
Class
Molecular function
Cellular component
Biological process

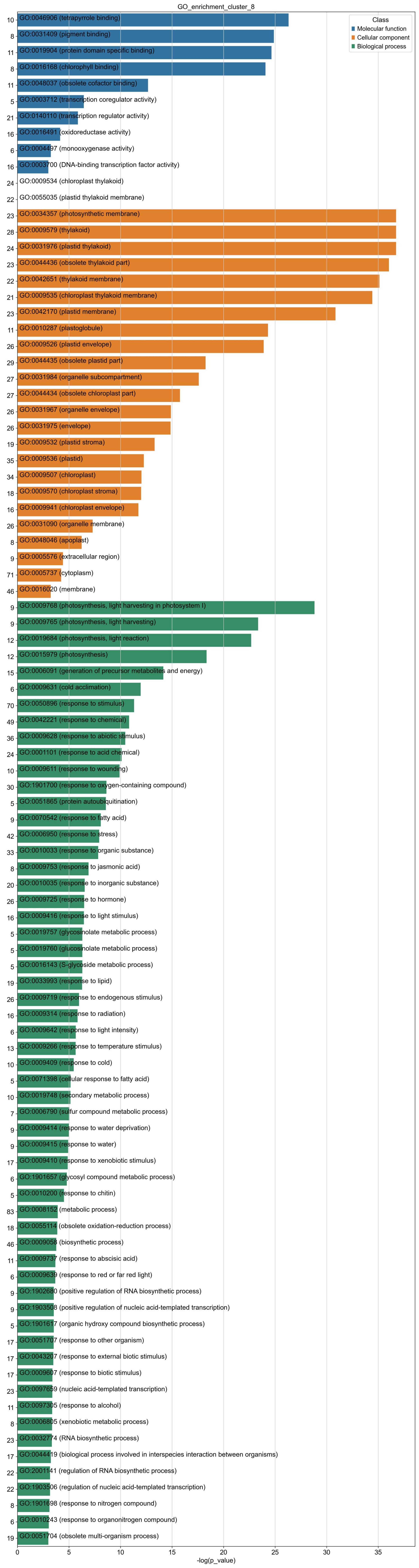

GO_enrichment_cluster_8
Class
Molecular function
Cellular component
Biological process
10 GO:0046906 (tetrapyrrole binding)
8 GO:0031409 (pigment binding)
11 GO:0019904 (protein domain specific binding)
8 GO:0016168 (chlorophyll binding)
11 GO:0048037 (obsolete cofactor binding)
5 GO:0003712 (transcription coregulator activity)
21 GO:0140110 (transcription regulator activity)
16 GO:0016491 (oxidoreductase activity)
6 GO:0004497 (monooxygenase activity)
16 GO:0003700 (DNA-binding transcription factor activity)
24 GO:0009534 (chloroplast thylakoid)
22 GO:0055035 (plastid thylakoid membrane)
23 GO:0034357 (photosynthetic membrane)
28 GO:0009579 (thylakoid)
24 GO:0031976 (plastid thylakoid)
23 GO:0044436 (obsolete thylakoid part)
22 GO:0042651 (thylakoid membrane)
21 GO:0009535 (chloroplast thylakoid membrane)
23 GO:0042170 (plastid membrane)
11 GO:0010287 (plastoglobule)
26 GO:0009526 (plastid envelope)
29 GO:0044435 (obsolete plastid part)
27 GO:0031984 (organelle subcompartment)
27 GO:0044434 (obsolete chloroplast part)
26 GO:0031967 (organelle envelope)
26 GO:0031975 (envelope)
19 GO:0009532 (plastid stroma)
35 GO:0009536 (plastid)
34 GO:0009507 (chloroplast)
18 GO:0009570 (chloroplast stroma)
16 GO:0009941 (chloroplast envelope)
26 GO:0031090 (organelle membrane)
8 GO:0048046 (apoplast)
9 GO:0005576 (extracellular region)
71 GO:0005737 (cytoplasm)
46 GO:0016020 (membrane)
9 GO:0009768 (photosynthesis, light harvesting in photosystem I)
9 GO:0009765 (photosynthesis, light harvesting)
12 GO:0019684 (photosynthesis, light reaction)
12 GO:0015979 (photosynthesis)
15 GO:0006091 (generation of precursor metabolites and energy)
6 GO:0009631 (cold acclimation)
70 GO:0050896 (response to stimulus)
49 GO:0042221 (response to chemical)
36 GO:0009628 (response to abiotic stimulus)
24 GO:0001101 (response to acid chemical)
10 GO:0009611 (response to wounding)
30 GO:1901700 (response to oxygen-containing compound)
5 GO:0051865 (protein autoubiquitination)
9 GO:0070542 (response to fatty acid)
42 GO:0006950 (response to stress)
33 GO:0010033 (response to organic substance)
8 GO:0009753 (response to jasmonic acid)
20 GO:0010035 (response to inorganic substance)
26 GO:0009725 (response to hormone)
16 GO:0009416 (response to light stimulus)
5 GO:0019757 (glycosinolate metabolic process)
5 GO:0019760 (glucosinolate metabolic process)
5 GO:0016143 (S-glycoside metabolic process)
19 GO:0033993 (response to lipid)
26 GO:0009719 (response to endogenous stimulus)
16 GO:0009314 (response to radiation)
6 GO:0009642 (response to light intensity)
13 GO:0009266 (response to temperature stimulus)
10 GO:0009409 (response to cold)
5 GO:0071398 (cellular response to fatty acid)
10 GO:0019748 (secondary metabolic process)
7 GO:0006790 (sulfur compound metabolic process)
9 GO:0009414 (response to water deprivation)
9 GO:0009415 (response to water)
17 GO:0009410 (response to xenobiotic stimulus)
6 GO:1901657 (glycosyl compound metabolic process)
5 GO:0010200 (response to chitin)
83 GO:0008152 (metabolic process)
18 GO:0055114 (obsolete oxidation-reduction process)
46 GO:0009058 (biosynthetic process)
11 GO:0009737 (response to abscisic acid)
6 GO:0009639 (response to red or far red light)
9 GO:1902680 (positive regulation of RNA biosynthetic process)
9 GO:1903508 (positive regulation of nucleic acid-templated transcription)
5 GO:1901617 (organic hydroxy compound biosynthetic process)
17 GO:0051707 (response to other organism)
17 GO:0043207 (response to external biotic stimulus)
17 GO:0009607 (response to biotic stimulus)
23 GO:0097659 (nucleic acid-templated transcription)
11 GO:0097305 (response to alcohol)
8 GO:0006805 (xenobiotic metabolic process)
23 GO:0032774 (RNA biosynthetic process)
17 GO:0044419 (biological process involved in interspecies interaction between organisms)
22 GO:2001141 (regulation of RNA biosynthetic process)
22 GO:1903506 (regulation of nucleic acid-templated transcription)
8 GO:1901698 (response to nitrogen compound)
6 GO:0010243 (response to organonitrogen compound)
19 GO:0051704 (obsolete multi-organism process)
0
5
10
15
20
25
30
35
-log(p_value)

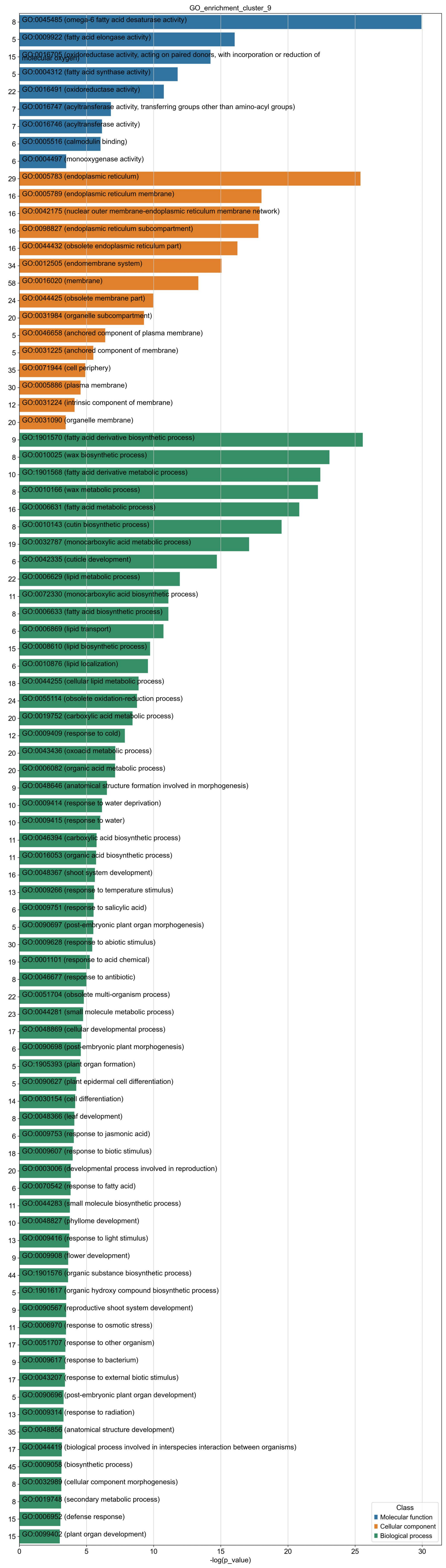
GO_enrichment_cluster_9
8 GO:0045485 (omega-6 fatty acid desaturase activity)
5 GO:0009922 (fatty acid elongase activity)
15 GO:0016705 (oxidoreductase activity, acting on paired donors, with incorporation or reduction of molecular oxygen)
5 GO:0004312 (fatty acid synthase activity)
22 GO:0016491 (oxidoreductase activity)
7 GO:0016747 (acyltransferase activity, transferring groups other than amino-acyl groups)
7 GO:0016746 (acyltransferase activity)
6 GO:0005516 (calmodulin binding)
6 GO:0004497 (monooxygenase activity)
29 GO:0005783 (endoplasmic reticulum)
16 GO:0005789 (endoplasmic reticulum membrane)
16 GO:0042175 (nuclear outer membrane-endoplasmic reticulum membrane network)
16 GO:0098827 (endoplasmic reticulum subcompartment)
16 GO:0044432 (obsolete endoplasmic reticulum part)
34 GO:0012505 (endomembrane system)
58 GO:0016020 (membrane)
24 GO:0044425 (obsolete membrane part)
20 GO:0031984 (organelle subcompartment)
5 GO:0046658 (anchored component of plasma membrane)
5 GO:0031225 (anchored component of membrane)
35 GO:0071944 (cell periphery)
30 GO:0005886 (plasma membrane)
12 GO:0031224 (intrinsic component of membrane)
20 GO:0031090 (organelle membrane)
9 GO:1901570 (fatty acid derivative biosynthetic process)
8 GO:0010025 (wax biosynthetic process)
10 GO:1901568 (fatty acid derivative metabolic process)
8 GO:0010166 (wax metabolic process)
16 GO:0006631 (fatty acid metabolic process)
8 GO:0010143 (cutin biosynthetic process)
19 GO:0032787 (monocarboxylic acid metabolic process)
6 GO:0042335 (cuticle development)
22 GO:0006629 (lipid metabolic process)
11 GO:0072330 (monocarboxylic acid biosynthetic process)
8 GO:0006633 (fatty acid biosynthetic process)
6 GO:0006869 (lipid transport)
15 GO:0008610 (lipid biosynthetic process)
6 GO:0010876 (lipid localization)
18 GO:0044255 (cellular lipid metabolic process)
24 GO:0055114 (obsolete oxidation-reduction process)
20 GO:0019752 (carboxylic acid metabolic process)
12 GO:0009409 (response to cold)
20 GO:0043436 (oxoacid metabolic process)
20 GO:0006082 (organic acid metabolic process)
9 GO:0048646 (anatomical structure formation involved in morphogenesis)
10 GO:0009414 (response to water deprivation)
10 GO:0009415 (response to water)
11 GO:0046394 (carboxylic acid biosynthetic process)
11 GO:0016053 (organic acid biosynthetic process)
16 GO:0048367 (shoot system development)
13 GO:0009266 (response to temperature stimulus)
6 GO:0009751 (response to salicylic acid)
5 GO:0090697 (post-embryonic plant organ morphogenesis)
30 GO:0009628 (response to abiotic stimulus)
19 GO:0001101 (response to acid chemical)
8 GO:0046677 (response to antibiotic)
22 GO:0051704 (obsolete multi-organism process)
23 GO:0044281 (small molecule metabolic process)
17 GO:0048869 (cellular developmental process)
6 GO:0090698 (post-embryonic plant morphogenesis)
5 GO:1905393 (plant organ formation)
5 GO:0090627 (plant epidermal cell differentiation)
14 GO:0030154 (cell differentiation)
8 GO:0048366 (leaf development)
6 GO:0009753 (response to jasmonic acid)
18 GO:0009607 (response to biotic stimulus)
20 GO:0003006 (developmental process involved in reproduction)
6 GO:0070542 (response to fatty acid)
11 GO:0044283 (small molecule biosynthetic process)
10 GO:0048827 (phyllome development)
13 GO:0009416 (response to light stimulus)
9 GO:0009908 (flower development)
44 GO:1901576 (organic substance biosynthetic process)
5 GO:1901617 (organic hydroxy compound biosynthetic process)
9 GO:0090567 (reproductive shoot system development)
11 GO:0006970 (response to osmotic stress)
17 GO:0051707 (response to other organism)
9 GO:0009617 (response to bacterium)
17 GO:0043207 (response to external biotic stimulus)
5 GO:0090696 (post-embryonic plant organ development)
13 GO:0009314 (response to radiation)
35 GO:0048856 (anatomical structure development)
17 GO:0044419 (biological process involved in interspecies interaction between organisms)
45 GO:0009058 (biosynthetic process)
8 GO:0032989 (cellular component morphogenesis)
8 GO:0019748 (secondary metabolic process)
15 GO:0006952 (defense response)
15 GO:0099402 (plant organ development)
Class
Molecular function
Cellular component
Biological process
0
5
10
15
20
25
30
-log(p_value)

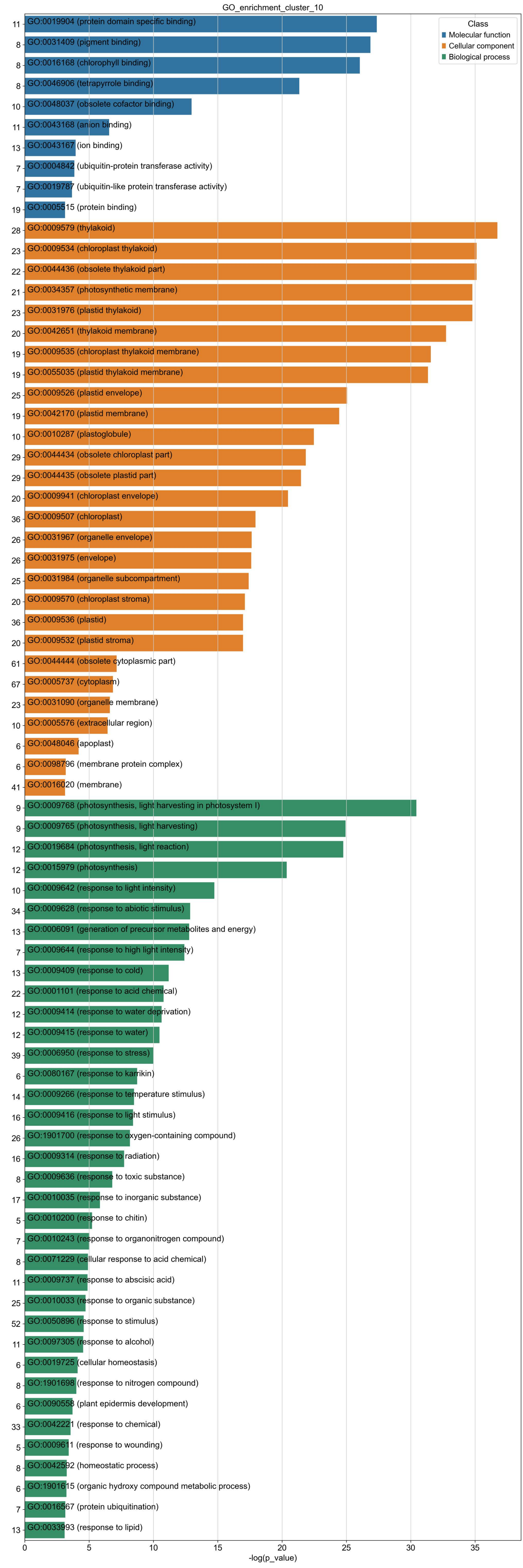
GO_enrichment_cluster_10
Class
Molecular function
Cellular component
Biological process
11
GO:0019904 (protein domain specific binding)
8
GO:0031409 (pigment binding)
8
GO:0016168 (chlorophyll binding)
8
GO:0046906 (tetrapyrrole binding)
10
GO:0048037 (obsolete cofactor binding)
11
GO:0043168 (anion binding)
13
GO:0043167 (ion binding)
7
GO:0004842 (ubiquitin-protein transferase activity)
7
GO:0019787 (ubiquitin-like protein transferase activity)
19
GO:0005515 (protein binding)
28
GO:0009579 (thylakoid)
23
GO:0009534 (chloroplast thylakoid)
22
GO:0044436 (obsolete thylakoid part)
21
GO:0034357 (photosynthetic membrane)
23
GO:0031976 (plastid thylakoid)
20
GO:0042651 (thylakoid membrane)
19
GO:0009535 (chloroplast thylakoid membrane)
19
GO:0055035 (plastid thylakoid membrane)
25
GO:0009526 (plastid envelope)
19
GO:0042170 (plastid membrane)
10
GO:0010287 (plastoglobule)
29
GO:0044434 (obsolete chloroplast part)
29
GO:0044435 (obsolete plastid part)
20
GO:0009941 (chloroplast envelope)
36
GO:0009507 (chloroplast)
26
GO:0031967 (organelle envelope)
26
GO:0031975 (envelope)
25
GO:0031984 (organelle subcompartment)
20
GO:0009570 (chloroplast stroma)
36
GO:0009536 (plastid)
20
GO:0009532 (plastid stroma)
61
GO:0044444 (obsolete cytoplasmic part)
67
GO:0005737 (cytoplasm)
23
GO:0031090 (organelle membrane)
10
GO:0005576 (extracellular region)
6
GO:0048046 (apoplast)
6
GO:0098796 (membrane protein complex)
41
GO:0016020 (membrane)
9
GO:0009768 (photosynthesis, light harvesting in photosystem I)
9
GO:0009765 (photosynthesis, light harvesting)
12
GO:0019684 (photosynthesis, light reaction)
12
GO:0015979 (photosynthesis)
10
GO:0009642 (response to light intensity)
34
GO:0009628 (response to abiotic stimulus)
13
GO:0006091 (generation of precursor metabolites and energy)
7
GO:0009644 (response to high light intensity)
13
GO:0009409 (response to cold)
22
GO:0001101 (response to acid chemical)
12
GO:0009414 (response to water deprivation)
12
GO:0009415 (response to water)
39
GO:0006950 (response to stress)
6
GO:0080167 (response to karrikin)
14
GO:0009266 (response to temperature stimulus)
16
GO:0009416 (response to light stimulus)
26
GO:1901700 (response to oxygen-containing compound)
16
GO:0009314 (response to radiation)
8
GO:0009636 (response to toxic substance)
17
GO:0010035 (response to inorganic substance)
5
GO:0010200 (response to chitin)
7
GO:0010243 (response to organonitrogen compound)
8
GO:0071229 (cellular response to acid chemical)
11
GO:0009737 (response to abscisic acid)
25
GO:0010033 (response to organic substance)
52
GO:0050896 (response to stimulus)
11
GO:0097305 (response to alcohol)
6
GO:0019725 (cellular homeostasis)
8
GO:1901698 (response to nitrogen compound)
6
GO:0090558 (plant epidermis development)
33
GO:0042221 (response to chemical)
5
GO:0009611 (response to wounding)
8
GO:0042592 (homeostatic process)
6
GO:1901615 (organic hydroxy compound metabolic process)
7
GO:0016567 (protein ubiquitination)
13
GO:0033993 (response to lipid)
0
5
10
15
20
25
30
35
-log(p_value)

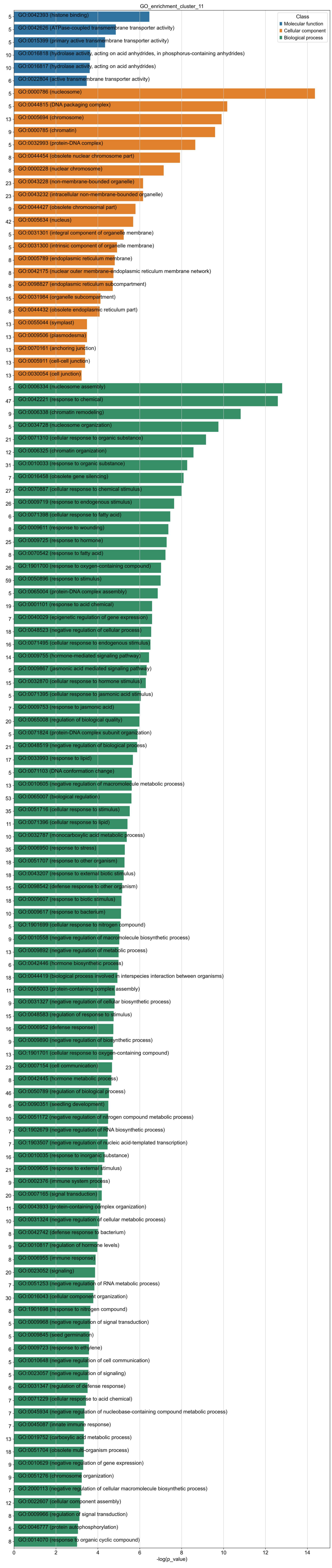

GO_enrichment_cluster_11
Class
Molecular function
Cellular component
Biological process
5 GO:0042393 (histone binding)
5 GO:0042626 (ATPase-coupled transmembrane transporter activity)
5 GO:0015399 (primary active transmembrane transporter activity)
10 GO:0016818 (hydrolase activity, acting on acid anhydrides, in phosphorus-containing anhydrides)
10 GO:0016817 (hydrolase activity, acting on acid anhydrides)
6 GO:0022804 (active transmembrane transporter activity)
5 GO:0000786 (nucleosome)
5 GO:0044815 (DNA packaging complex)
13 GO:0005694 (chromosome)
9 GO:0000785 (chromatin)
5 GO:0032993 (protein-DNA complex)
8 GO:0044454 (obsolete nuclear chromosome part)
8 GO:0000228 (nuclear chromosome)
23 GO:0043228 (non-membrane-bounded organelle)
23 GO:0043232 (intracellular non-membrane-bounded organelle)
9 GO:0044427 (obsolete chromosomal part)
42 GO:0005634 (nucleus)
5 GO:0031301 (integral component of organelle membrane)
5 GO:0031300 (intrinsic component of organelle membrane)
8 GO:0005789 (endoplasmic reticulum membrane)
8 GO:0042175 (nuclear outer membrane-endoplasmic reticulum membrane network)
8 GO:0098827 (endoplasmic reticulum subcompartment)
15 GO:0031984 (organelle subcompartment)
8 GO:0044432 (obsolete endoplasmic reticulum part)
13 GO:0055044 (symplast)
13 GO:0009506 (plasmodesma)
13 GO:0070161 (anchoring junction)
13 GO:0005911 (cell-cell junction)
13 GO:0030054 (cell junction)
5 GO:0006334 (nucleosome assembly)
47 GO:0042221 (response to chemical)
9 GO:0006338 (chromatin remodeling)
5 GO:0034728 (nucleosome organization)
21 GO:0071310 (cellular response to organic substance)
12 GO:0006325 (chromatin organization)
31 GO:0010033 (response to organic substance)
7 GO:0016458 (obsolete gene silencing)
27 GO:0070887 (cellular response to chemical stimulus)
26 GO:0009719 (response to endogenous stimulus)
6 GO:0071398 (cellular response to fatty acid)
8 GO:0009611 (response to wounding)
25 GO:0009725 (response to hormone)
8 GO:0070542 (response to fatty acid)
26 GO:1901700 (response to oxygen-containing compound)
59 GO:0050896 (response to stimulus)
5 GO:0065004 (protein-DNA complex assembly)
19 GO:0001101 (response to acid chemical)
7 GO:0040029 (epigenetic regulation of gene expression)
18 GO:0048523 (negative regulation of cellular process)
16 GO:0071495 (cellular response to endogenous stimulus)
14 GO:0009755 (hormone-mediated signaling pathway)
5 GO:0009867 (jasmonic acid mediated signaling pathway)
15 GO:0032870 (cellular response to hormone stimulus)
5 GO:0071395 (cellular response to jasmonic acid stimulus)
7 GO:0009753 (response to jasmonic acid)
20 GO:0065008 (regulation of biological quality)
5 GO:0071824 (protein-DNA complex subunit organization)
21 GO:0048519 (negative regulation of biological process)
17 GO:0033993 (response to lipid)
5 GO:0071103 (DNA conformation change)
13 GO:0010605 (negative regulation of macromolecule metabolic process)
53 GO:0065007 (biological regulation)
35 GO:0051716 (cellular response to stimulus)
11 GO:0071396 (cellular response to lipid)
10 GO:0032787 (monocarboxylic acid metabolic process)
35 GO:0006950 (response to stress)
18 GO:0051707 (response to other organism)
18 GO:0043207 (response to external biotic stimulus)
15 GO:0098542 (defense response to other organism)
18 GO:0009607 (response to biotic stimulus)
10 GO:0009617 (response to bacterium)
5 GO:1901699 (cellular response to nitrogen compound)
9 GO:0010558 (negative regulation of macromolecule biosynthetic process)
13 GO:0009892 (negative regulation of metabolic process)
6 GO:0042446 (hormone biosynthetic process)
18 GO:0044419 (biological process involved in interspecies interaction between organisms)
11 GO:0065003 (protein-containing complex assembly)
9 GO:0031327 (negative regulation of cellular biosynthetic process)
15 GO:0048583 (regulation of response to stimulus)
16 GO:0006952 (defense response)
9 GO:0009890 (negative regulation of biosynthetic process)
13 GO:1901701 (cellular response to oxygen-containing compound)
23 GO:0007154 (cell communication)
8 GO:0042445 (hormone metabolic process)
46 GO:0050789 (regulation of biological process)
6 GO:0090351 (seedling development)
10 GO:0051172 (negative regulation of nitrogen compound metabolic process)
7 GO:1902679 (negative regulation of RNA biosynthetic process)
7 GO:1903507 (negative regulation of nucleic acid-templated transcription)
16 GO:0010035 (response to inorganic substance)
21 GO:0009605 (response to external stimulus)
9 GO:0002376 (immune system process)
20 GO:0007165 (signal transduction)
11 GO:0043933 (protein-containing complex organization)
10 GO:0031324 (negative regulation of cellular metabolic process)
8 GO:0042742 (defense response to bacterium)
9 GO:0010817 (regulation of hormone levels)
8 GO:0006955 (immune response)
20 GO:0023052 (signaling)
7 GO:0051253 (negative regulation of RNA metabolic process)
30 GO:0016043 (cellular component organization)
8 GO:1901698 (response to nitrogen compound)
5 GO:0009968 (negative regulation of signal transduction)
5 GO:0009845 (seed germination)
6 GO:0009723 (response to ethylene)
5 GO:0010648 (negative regulation of cell communication)
5 GO:0023057 (negative regulation of signaling)
6 GO:0031347 (regulation of defense response)
7 GO:0071229 (cellular response to acid chemical)
7 GO:0045934 (negative regulation of nucleobase-containing compound metabolic process)
7 GO:0045087 (innate immune response)
13 GO:0019752 (carboxylic acid metabolic process)
18 GO:0051704 (obsolete multi-organism process)
9 GO:0010629 (negative regulation of gene expression)
9 GO:0051276 (chromosome organization)
7 GO:2000113 (negative regulation of cellular macromolecule biosynthetic process)
12 GO:0022607 (cellular component assembly)
8 GO:0009966 (regulation of signal transduction)
5 GO:0046777 (protein autophosphorylation)
8 GO:0014070 (response to organic cyclic compound)
0
2
4
6
8
10
12
14
-log(p_value)

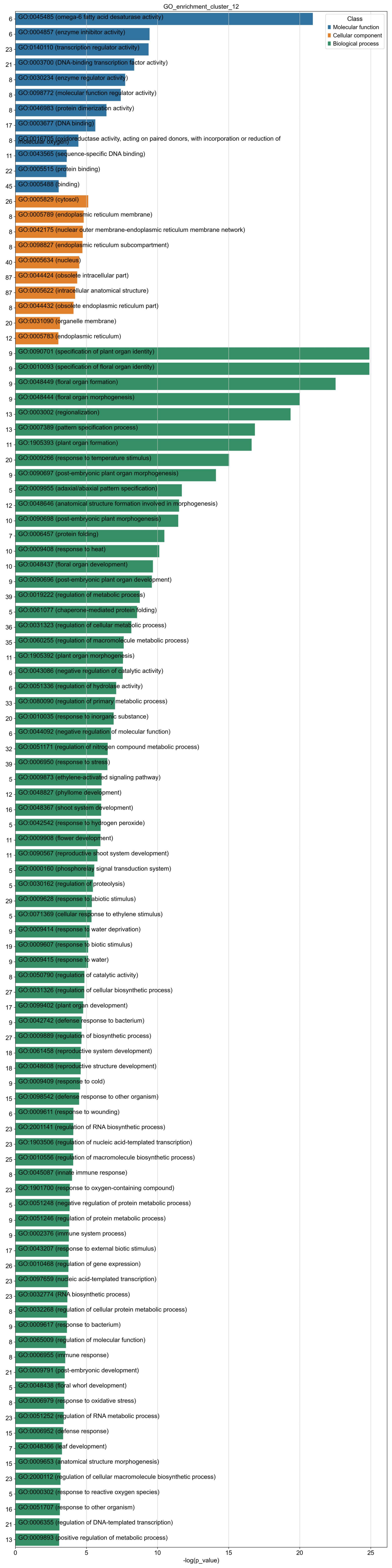
GO_enrichment_cluster_12
Class
Molecular function
Cellular component
Biological process
6 GO:0045485 (omega-6 fatty acid desaturase activity)
6 GO:0004857 (enzyme inhibitor activity)
23 GO:0140110 (transcription regulator activity)
21 GO:0003700 (DNA-binding transcription factor activity)
8 GO:0030234 (enzyme regulator activity)
8 GO:0098772 (molecular function regulator activity)
8 GO:0046983 (protein dimerization activity)
17 GO:0003677 (DNA binding)
8 GO:0016705 (oxidoreductase activity, acting on paired donors, with incorporation or reduction of molecular oxygen)
11 GO:0043565 (sequence-specific DNA binding)
22 GO:0005515 (protein binding)
45 GO:0005488 (binding)
26 GO:0005829 (cytosol)
8 GO:0005789 (endoplasmic reticulum membrane)
8 GO:0042175 (nuclear outer membrane-endoplasmic reticulum membrane network)
8 GO:0098827 (endoplasmic reticulum subcompartment)
40 GO:0005634 (nucleus)
87 GO:0044424 (obsolete intracellular part)
87 GO:0005622 (intracellular anatomical structure)
8 GO:0044432 (obsolete endoplasmic reticulum part)
20 GO:0031090 (organelle membrane)
12 GO:0005783 (endoplasmic reticulum)
9 GO:0090701 (specification of plant organ identity)
9 GO:0010093 (specification of floral organ identity)
9 GO:0048449 (floral organ formation)
9 GO:0048444 (floral organ morphogenesis)
13 GO:0003002 (regionalization)
13 GO:0007389 (pattern specification process)
11 GO:1905393 (plant organ formation)
20 GO:0009266 (response to temperature stimulus)
9 GO:0090697 (post-embryonic plant organ morphogenesis)
5 GO:0009955 (adaxial/abaxial pattern specification)
12 GO:0048646 (anatomical structure formation involved in morphogenesis)
10 GO:0090698 (post-embryonic plant morphogenesis)
7 GO:0006457 (protein folding)
10 GO:0009408 (response to heat)
10 GO:0048437 (floral organ development)
9 GO:0090696 (post-embryonic plant organ development)
39 GO:0019222 (regulation of metabolic process)
5 GO:0061077 (chaperone-mediated protein folding)
36 GO:0031323 (regulation of cellular metabolic process)
35 GO:0060255 (regulation of macromolecule metabolic process)
11 GO:1905392 (plant organ morphogenesis)
6 GO:0043086 (negative regulation of catalytic activity)
6 GO:0051336 (regulation of hydrolase activity)
33 GO:0080090 (regulation of primary metabolic process)
20 GO:0010035 (response to inorganic substance)
6 GO:0044092 (negative regulation of molecular function)
32 GO:0051171 (regulation of nitrogen compound metabolic process)
39 GO:0006950 (response to stress)
5 GO:0009873 (ethylene-activated signaling pathway)
12 GO:0048827 (phyllome development)
16 GO:0048367 (shoot system development)
5 GO:0042542 (response to hydrogen peroxide)
11 GO:0009908 (flower development)
11 GO:0090567 (reproductive shoot system development)
5 GO:0000160 (phosphorelay signal transduction system)
5 GO:0030162 (regulation of proteolysis)
29 GO:0009628 (response to abiotic stimulus)
5 GO:0071369 (cellular response to ethylene stimulus)
9 GO:0009414 (response to water deprivation)
19 GO:0009607 (response to biotic stimulus)
9 GO:0009415 (response to water)
8 GO:0050790 (regulation of catalytic activity)
27 GO:0031326 (regulation of cellular biosynthetic process)
17 GO:0099402 (plant organ development)
9 GO:0042742 (defense response to bacterium)
27 GO:0009889 (regulation of biosynthetic process)
18 GO:0061458 (reproductive system development)
18 GO:0048608 (reproductive structure development)
9 GO:0009409 (response to cold)
15 GO:0098542 (defense response to other organism)
6 GO:0009611 (response to wounding)
23 GO:2001141 (regulation of RNA biosynthetic process)
23 GO:1903506 (regulation of nucleic acid-templated transcription)
25 GO:0010556 (regulation of macromolecule biosynthetic process)
8 GO:0045087 (innate immune response)
23 GO:1901700 (response to oxygen-containing compound)
5 GO:0051248 (negative regulation of protein metabolic process)
9 GO:0051246 (regulation of protein metabolic process)
9 GO:0002376 (immune system process)
17 GO:0043207 (response to external biotic stimulus)
26 GO:0010468 (regulation of gene expression)
23 GO:0097659 (nucleic acid-templated transcription)
23 GO:0032774 (RNA biosynthetic process)
8 GO:0032268 (regulation of cellular protein metabolic process)
9 GO:0009617 (response to bacterium)
8 GO:0065009 (regulation of molecular function)
8 GO:0006955 (immune response)
21 GO:0009791 (post-embryonic development)
5 GO:0048438 (floral whorl development)
8 GO:0006979 (response to oxidative stress)
23 GO:0051252 (regulation of RNA metabolic process)
15 GO:0006952 (defense response)
7 GO:0048366 (leaf development)
15 GO:0009653 (anatomical structure morphogenesis)
23 GO:2000112 (regulation of cellular macromolecule biosynthetic process)
5 GO:0000302 (response to reactive oxygen species)
16 GO:0051707 (response to other organism)
21 GO:0006355 (regulation of DNA-templated transcription)
13 GO:0009893 (positive regulation of metabolic process)
0
5
10
15
20
25
-log(p_value)

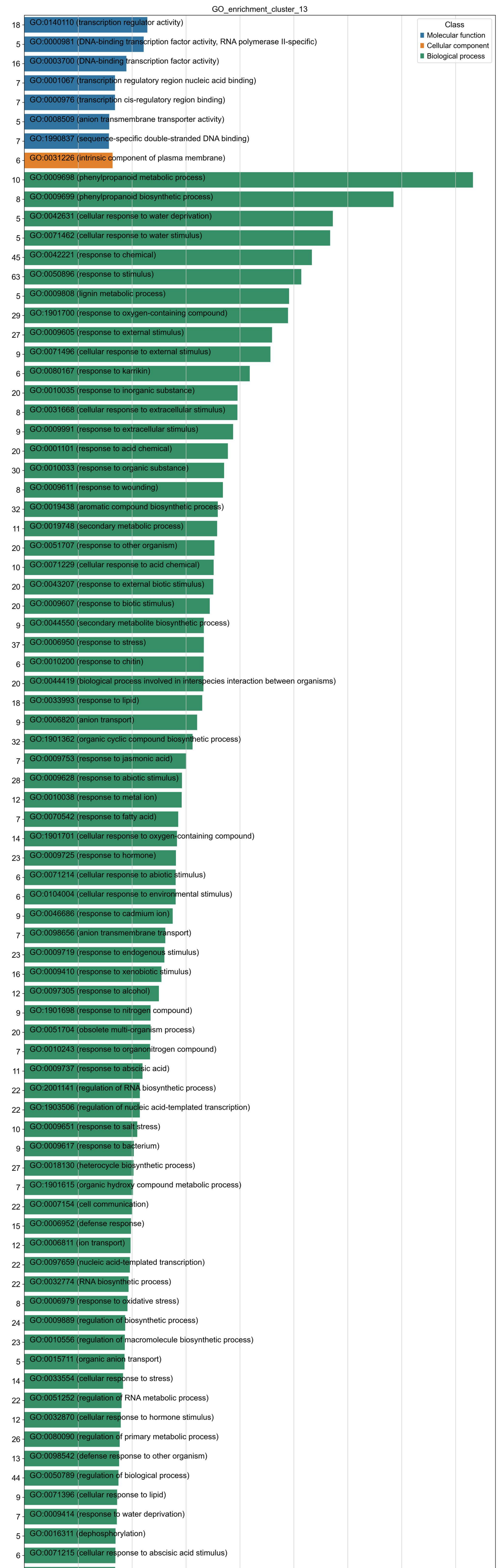
GO_enrichment_cluster_13
Class
Molecular function
Cellular component
Biological process
18 GO:0140110 (transcription regulator activity)
5 GO:0000981 (DNA-binding transcription factor activity, RNA polymerase II-specific)
16 GO:0003700 (DNA-binding transcription factor activity)
7 GO:0001067 (transcription regulatory region nucleic acid binding)
7 GO:0000976 (transcription cis-regulatory region binding)
5 GO:0008509 (anion transmembrane transporter activity)
7 GO:1990837 (sequence-specific double-stranded DNA binding)
6 GO:0031226 (intrinsic component of plasma membrane)
10 GO:0009698 (phenylpropanoid metabolic process)
8 GO:0009699 (phenylpropanoid biosynthetic process)
5 GO:0042631 (cellular response to water deprivation)
5 GO:0071462 (cellular response to water stimulus)
45 GO:0042221 (response to chemical)
63 GO:0050896 (response to stimulus)
5 GO:0009808 (lignin metabolic process)
29 GO:1901700 (response to oxygen-containing compound)
27 GO:0009605 (response to external stimulus)
9 GO:0071496 (cellular response to external stimulus)
6 GO:0080167 (response to karrikin)
20 GO:0010035 (response to inorganic substance)
8 GO:0031668 (cellular response to extracellular stimulus)
9 GO:0009991 (response to extracellular stimulus)
20 GO:0001101 (response to acid chemical)
30 GO:0010033 (response to organic substance)
8 GO:0009611 (response to wounding)
32 GO:0019438 (aromatic compound biosynthetic process)
11 GO:0019748 (secondary metabolic process)
20 GO:0051707 (response to other organism)
10 GO:0071229 (cellular response to acid chemical)
20 GO:0043207 (response to external biotic stimulus)
20 GO:0009607 (response to biotic stimulus)
9 GO:0044550 (secondary metabolite biosynthetic process)
37 GO:0006950 (response to stress)
6 GO:0010200 (response to chitin)
20 GO:0044419 (biological process involved in interspecies interaction between organisms)
18 GO:0033993 (response to lipid)
9 GO:0006820 (anion transport)
32 GO:1901362 (organic cyclic compound biosynthetic process)
7 GO:0009753 (response to jasmonic acid)
28 GO:0009628 (response to abiotic stimulus)
12 GO:0010038 (response to metal ion)
7 GO:0070542 (response to fatty acid)
14 GO:1901701 (cellular response to oxygen-containing compound)
23 GO:0009725 (response to hormone)
6 GO:0071214 (cellular response to abiotic stimulus)
6 GO:0104004 (cellular response to environmental stimulus)
9 GO:0046686 (response to cadmium ion)
7 GO:0098656 (anion transmembrane transport)
23 GO:0009719 (response to endogenous stimulus)
16 GO:0009410 (response to xenobiotic stimulus)
12 GO:0097305 (response to alcohol)
9 GO:1901698 (response to nitrogen compound)
20 GO:0051704 (obsolete multi-organism process)
7 GO:0010243 (response to organonitrogen compound)
11 GO:0009737 (response to abscisic acid)
22 GO:2001141 (regulation of RNA biosynthetic process)
22 GO:1903506 (regulation of nucleic acid-templated transcription)
10 GO:0009651 (response to salt stress)
9 GO:0009617 (response to bacterium)
27 GO:0018130 (heterocycle biosynthetic process)
7 GO:1901615 (organic hydroxy compound metabolic process)
22 GO:0007154 (cell communication)
15 GO:0006952 (defense response)
12 GO:0006811 (ion transport)
22 GO:0097659 (nucleic acid-templated transcription)
22 GO:0032774 (RNA biosynthetic process)
8 GO:0006979 (response to oxidative stress)
24 GO:0009889 (regulation of biosynthetic process)
23 GO:0010556 (regulation of macromolecule biosynthetic process)
5 GO:0015711 (organic anion transport)
14 GO:0033554 (cellular response to stress)
22 GO:0051252 (regulation of RNA metabolic process)
12 GO:0032870 (cellular response to hormone stimulus)
26 GO:0080090 (regulation of primary metabolic process)
13 GO:0098542 (defense response to other organism)
44 GO:0050789 (regulation of biological process)
9 GO:0071396 (cellular response to lipid)
7 GO:0009414 (response to water deprivation)
5 GO:0016311 (dephosphorylation)
6 GO:0071215 (cellular response to abscisic acid stimulus)
6 GO:0097306 (cellular response to alcohol)
10 GO:0006970 (response to osmotic stress)
10 GO:0034220 (ion transmembrane transport)
29 GO:0019222 (regulation of metabolic process)
7 GO:0009415 (response to water)
8 GO:0002376 (immune system process)
20 GO:0006355 (regulation of DNA-templated transcription)
22 GO:0019219 (regulation of nucleobase-containing compound metabolic process)
25 GO:0051171 (regulation of nitrogen compound metabolic process)
12 GO:0071495 (cellular response to endogenous stimulus)
0 2 4 6 8 10 12 14 16
-log(p_value)

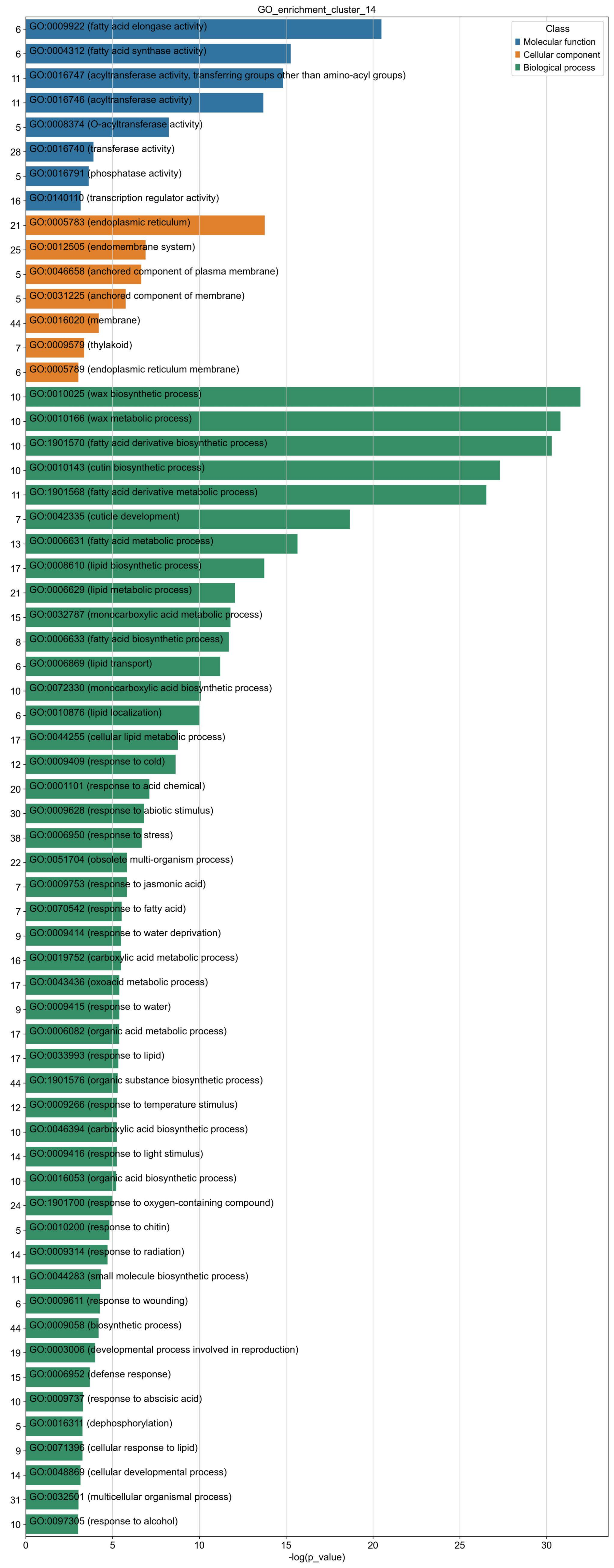
GO_enrichment_cluster_14
Class
Molecular function
Cellular component
Biological process
6 GO:0009922 (fatty acid elongase activity)
6 GO:0004312 (fatty acid synthase activity)
11 GO:0016747 (acyltransferase activity, transferring groups other than amino-acyl groups)
11 GO:0016746 (acyltransferase activity)
5 GO:0008374 (O-acyltransferase activity)
28 GO:0016740 (transferase activity)
5 GO:0016791 (phosphatase activity)
16 GO:0140110 (transcription regulator activity)
21 GO:0005783 (endoplasmic reticulum)
25 GO:0012505 (endomembrane system)
5 GO:0046658 (anchored component of plasma membrane)
5 GO:0031225 (anchored component of membrane)
44 GO:0016020 (membrane)
7 GO:0009579 (thylakoid)
6 GO:0005789 (endoplasmic reticulum membrane)
10 GO:0010025 (wax biosynthetic process)
10 GO:0010166 (wax metabolic process)
10 GO:1901570 (fatty acid derivative biosynthetic process)
10 GO:0010143 (cutin biosynthetic process)
11 GO:1901568 (fatty acid derivative metabolic process)
7 GO:0042335 (cuticle development)
13 GO:0006631 (fatty acid metabolic process)
17 GO:0008610 (lipid biosynthetic process)
21 GO:0006629 (lipid metabolic process)
15 GO:0032787 (monocarboxylic acid metabolic process)
8 GO:0006633 (fatty acid biosynthetic process)
6 GO:0006869 (lipid transport)
10 GO:0072330 (monocarboxylic acid biosynthetic process)
6 GO:0010876 (lipid localization)
17 GO:0044255 (cellular lipid metabolic process)
12 GO:0009409 (response to cold)
20 GO:0001101 (response to acid chemical)
30 GO:0009628 (response to abiotic stimulus)
38 GO:0006950 (response to stress)
22 GO:0051704 (obsolete multi-organism process)
7 GO:0009753 (response to jasmonic acid)
7 GO:0070542 (response to fatty acid)
9 GO:0009414 (response to water deprivation)
16 GO:0019752 (carboxylic acid metabolic process)
17 GO:0043436 (oxoacid metabolic process)
9 GO:0009415 (response to water)
17 GO:0006082 (organic acid metabolic process)
17 GO:0033993 (response to lipid)
44 GO:1901576 (organic substance biosynthetic process)
12 GO:0009266 (response to temperature stimulus)
10 GO:0046394 (carboxylic acid biosynthetic process)
14 GO:0009416 (response to light stimulus)
10 GO:0016053 (organic acid biosynthetic process)
24 GO:1901700 (response to oxygen-containing compound)
5 GO:0010200 (response to chitin)
14 GO:0009314 (response to radiation)
11 GO:0044283 (small molecule biosynthetic process)
6 GO:0009611 (response to wounding)
44 GO:0009058 (biosynthetic process)
19 GO:0003006 (developmental process involved in reproduction)
15 GO:0006952 (defense response)
10 GO:0009737 (response to abscisic acid)
5 GO:0016311 (dephosphorylation)
9 GO:0071396 (cellular response to lipid)
14 GO:0048869 (cellular developmental process)
31 GO:0032501 (multicellular organismal process)
10 GO:0097305 (response to alcohol)
0
5
10
15
20
25
30
-log(p_value)

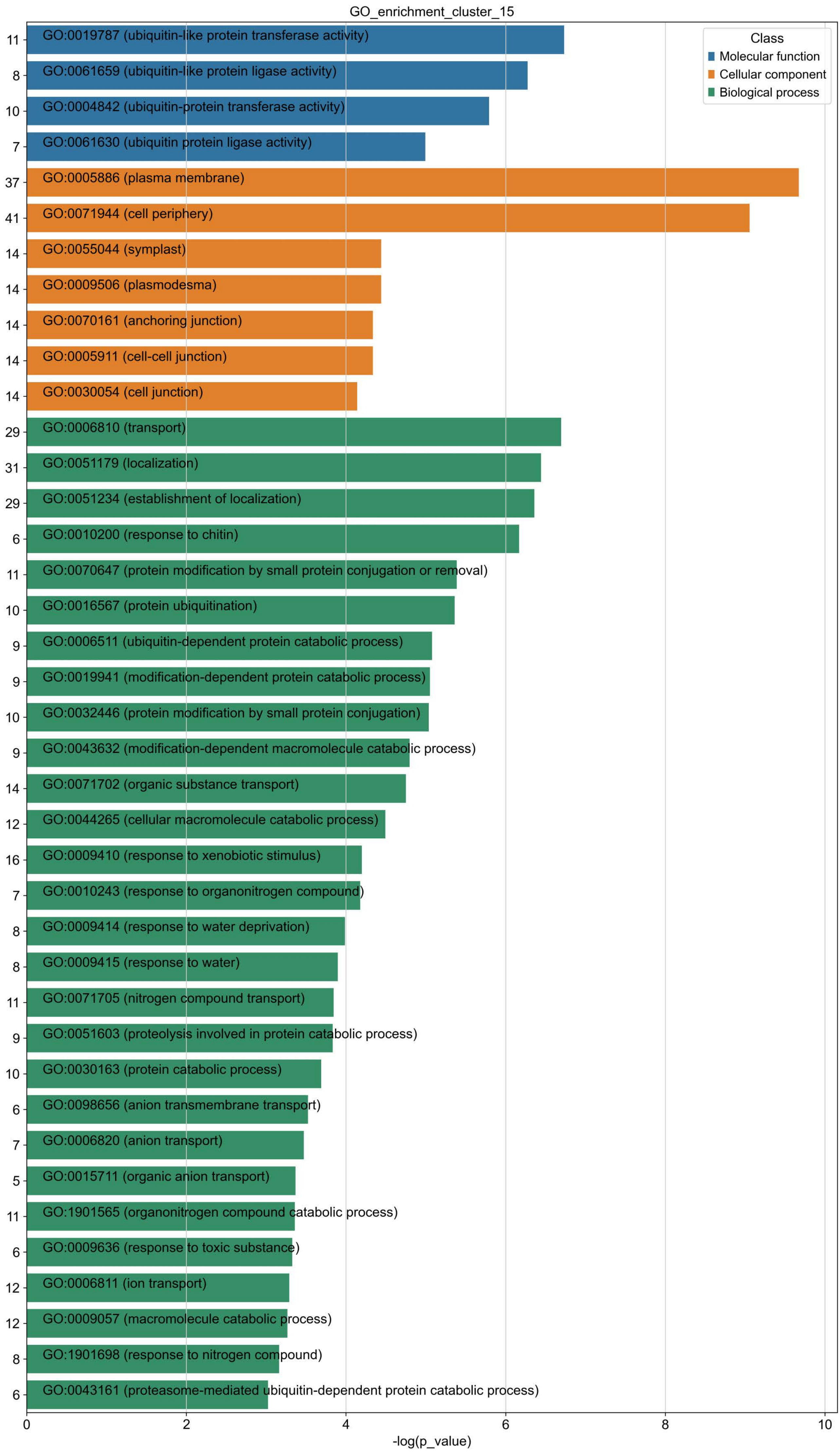

GO_enrichment_cluster_15
Class
Molecular function
Cellular component
Biological process
11 GO:0019787 (ubiquitin-like protein transferase activity)
8 GO:0061659 (ubiquitin-like protein ligase activity)
10 GO:0004842 (ubiquitin-protein transferase activity)
7 GO:0061630 (ubiquitin protein ligase activity)
37 GO:0005886 (plasma membrane)
41 GO:0071944 (cell periphery)
14 GO:0055044 (symplast)
14 GO:0009506 (plasmodesma)
14 GO:0070161 (anchoring junction)
14 GO:0005911 (cell-cell junction)
14 GO:0030054 (cell junction)
29 GO:0006810 (transport)
31 GO:0051179 (localization)
29 GO:0051234 (establishment of localization)
6 GO:0010200 (response to chitin)
11 GO:0070647 (protein modification by small protein conjugation or removal)
10 GO:0016567 (protein ubiquitination)
9 GO:0006511 (ubiquitin-dependent protein catabolic process)
9 GO:0019941 (modification-dependent protein catabolic process)
10 GO:0032446 (protein modification by small protein conjugation)
9 GO:0043632 (modification-dependent macromolecule catabolic process)
14 GO:0071702 (organic substance transport)
12 GO:0044265 (cellular macromolecule catabolic process)
16 GO:0009410 (response to xenobiotic stimulus)
7 GO:0010243 (response to organonitrogen compound)
8 GO:0009414 (response to water deprivation)
8 GO:0009415 (response to water)
11 GO:0071705 (nitrogen compound transport)
9 GO:0051603 (proteolysis involved in protein catabolic process)
10 GO:0030163 (protein catabolic process)
6 GO:0098656 (anion transmembrane transport)
7 GO:0006820 (anion transport)
5 GO:0015711 (organic anion transport)
11 GO:1901565 (organonitrogen compound catabolic process)
6 GO:0009636 (response to toxic substance)
12 GO:0006811 (ion transport)
12 GO:0009057 (macromolecule catabolic process)
8 GO:1901698 (response to nitrogen compound)
6 GO:0043161 (proteasome-mediated ubiquitin-dependent protein catabolic process)
0
2
4
6
8
10
-log(p_value)

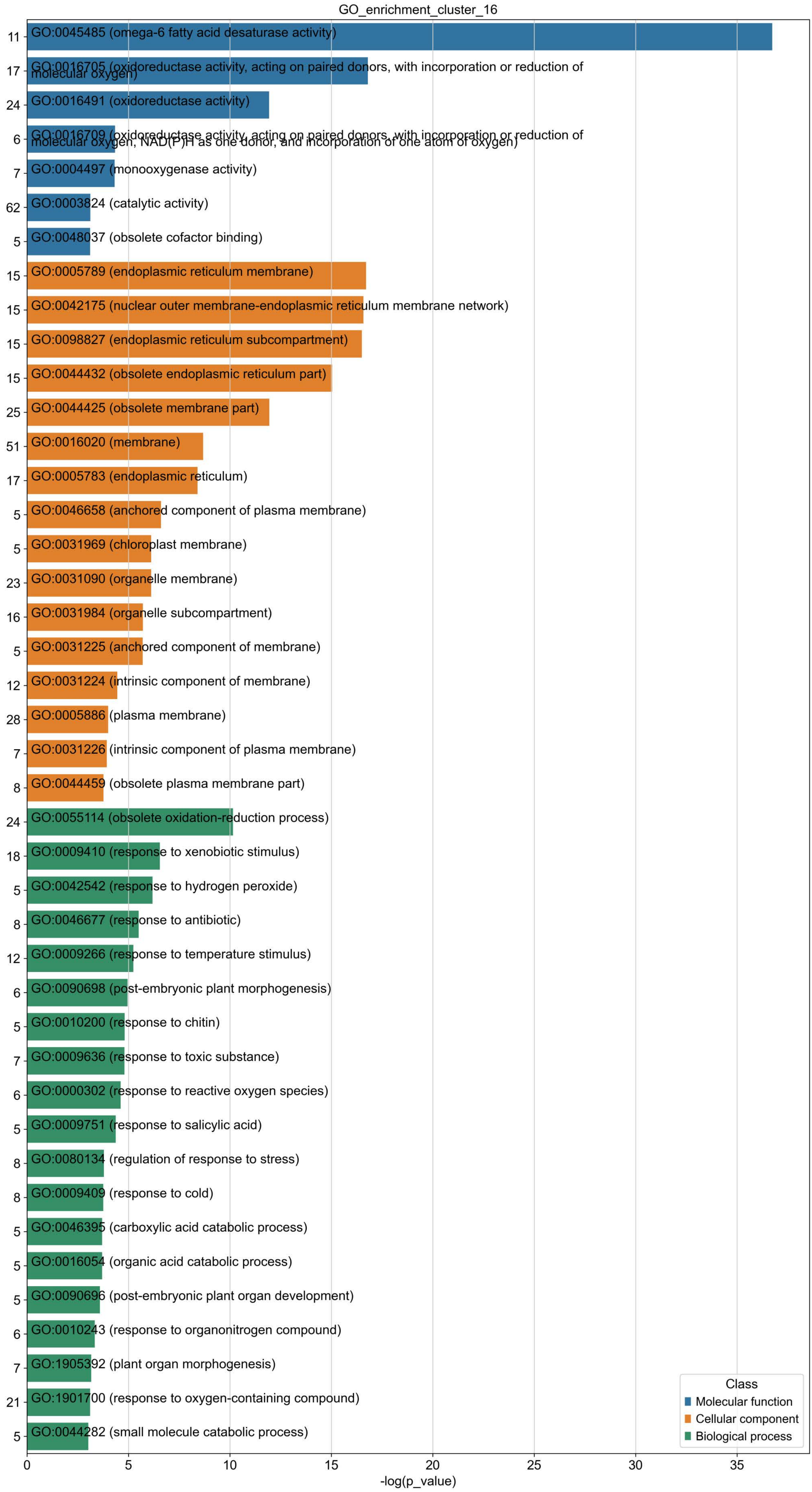

GO_enrichment_cluster_16
11
GO:0045485 (omega-6 fatty acid desaturase activity)
17
GO:0016705 (oxidoreductase activity, acting on paired donors, with incorporation or reduction of molecular oxygen)
24
GO:0016491 (oxidoreductase activity)
6
GO:0016709 (oxidoreductase activity, acting on paired donors, with incorporation or reduction of molecular oxygen, NAD(P)H as one donor, and incorporation of one atom of oxygen)
7
GO:0004497 (monooxygenase activity)
62
GO:0003824 (catalytic activity)
5
GO:0048037 (obsolete cofactor binding)
15
GO:0005789 (endoplasmic reticulum membrane)
15
GO:0042175 (nuclear outer membrane-endoplasmic reticulum membrane network)
15
GO:0098827 (endoplasmic reticulum subcompartment)
15
GO:0044432 (obsolete endoplasmic reticulum part)
25
GO:0044425 (obsolete membrane part)
51
GO:0016020 (membrane)
17
GO:0005783 (endoplasmic reticulum)
5
GO:0046658 (anchored component of plasma membrane)
5
GO:0031969 (chloroplast membrane)
23
GO:0031090 (organelle membrane)
16
GO:0031984 (organelle subcompartment)
5
GO:0031225 (anchored component of membrane)
12
GO:0031224 (intrinsic component of membrane)
28
GO:0005886 (plasma membrane)
7
GO:0031226 (intrinsic component of plasma membrane)
8
GO:0044459 (obsolete plasma membrane part)
24
GO:0055114 (obsolete oxidation-reduction process)
18
GO:0009410 (response to xenobiotic stimulus)
5
GO:0042542 (response to hydrogen peroxide)
8
GO:0046677 (response to antibiotic)
12
GO:0009266 (response to temperature stimulus)
6
GO:0090698 (post-embryonic plant morphogenesis)
5
GO:0010200 (response to chitin)
7
GO:0009636 (response to toxic substance)
6
GO:0000302 (response to reactive oxygen species)
5
GO:0009751 (response to salicylic acid)
8
GO:0080134 (regulation of response to stress)
8
GO:0009409 (response to cold)
5
GO:0046395 (carboxylic acid catabolic process)
5
GO:0016054 (organic acid catabolic process)
5
GO:0090696 (post-embryonic plant organ development)
6
GO:0010243 (response to organonitrogen compound)
7
GO:1905392 (plant organ morphogenesis)
21
GO:1901700 (response to oxygen-containing compound)
5
GO:0044282 (small molecule catabolic process)
Class
Molecular function
Cellular component
Biological process
0
5
10
15
20
25
30
35
-log(p_value)

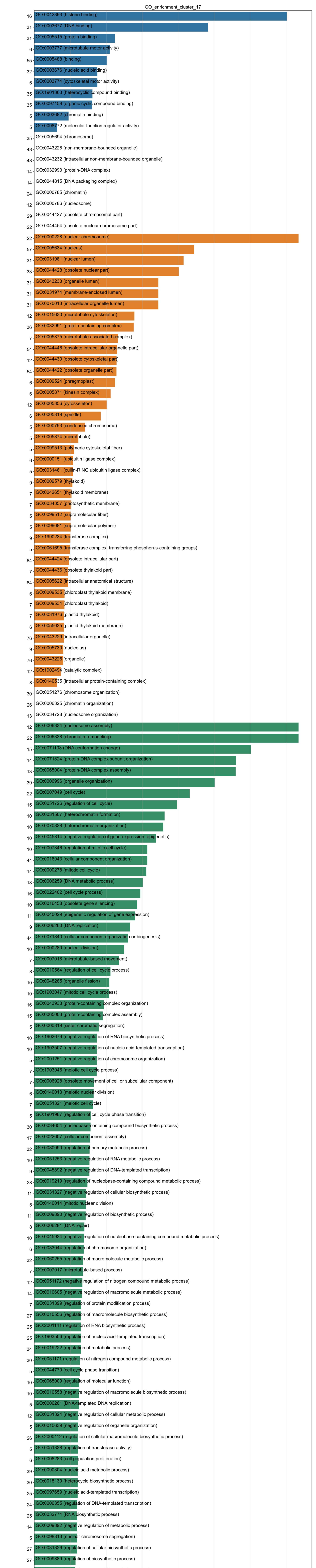
GO_enrichment_cluster_17
16 GO:0042393 (histone binding)
31 GO:0003677 (DNA binding)
31 GO:0005515 (protein binding)
6 GO:0003777 (microtubule motor activity)
55 GO:0005488 (binding)
32 GO:0003676 (nucleic acid binding)
6 GO:0003774 (cytoskeletal motor activity)
35 GO:1901363 (heterocyclic compound binding)
35 GO:0097159 (organic cyclic compound binding)
5 GO:0003682 (chromatin binding)
5 GO:0098772 (molecular function regulator activity)
35 GO:0005694 (chromosome)
48 GO:0043228 (non-membrane-bounded organelle)
48 GO:0043232 (intracellular non-membrane-bounded organelle)
14 GO:0032993 (protein-DNA complex)
14 GO:0044815 (DNA packaging complex)
24 GO:0000785 (chromatin)
12 GO:0000786 (nucleosome)
29 GO:0044427 (obsolete chromosomal part)
22 GO:0044454 (obsolete nuclear chromosome part)
22 GO:0000228 (nuclear chromosome)
57 GO:0005634 (nucleus)
31 GO:0031981 (nuclear lumen)
33 GO:0044428 (obsolete nuclear part)
31 GO:0043233 (organelle lumen)
31 GO:0031974 (membrane-enclosed lumen)
31 GO:0070013 (intracellular organelle lumen)
12 GO:0015630 (microtubule cytoskeleton)
36 GO:0032991 (protein-containing complex)
7 GO:0005875 (microtubule associated complex)
54 GO:0044446 (obsolete intracellular organelle part)
12 GO:0044430 (obsolete cytoskeletal part)
54 GO:0044422 (obsolete organelle part)
6 GO:0009524 (phragmoplast)
6 GO:0005871 (kinesin complex)
12 GO:0005856 (cytoskeleton)
6 GO:0005819 (spindle)
5 GO:0000793 (condensed chromosome)
5 GO:0005874 (microtubule)
5 GO:0099513 (polymeric cytoskeletal fiber)
6 GO:0000151 (ubiquitin ligase complex)
5 GO:0031461 (cullin-RING ubiquitin ligase complex)
9 GO:0009579 (thylakoid)
7 GO:0042651 (thylakoid membrane)
7 GO:0034357 (photosynthetic membrane)
5 GO:0099512 (supramolecular fiber)
5 GO:0099081 (supramolecular polymer)
9 GO:1990234 (transferase complex)
5 GO:0061695 (transferase complex, transferring phosphorus-containing groups)
84 GO:0044424 (obsolete intracellular part)
7 GO:0044436 (obsolete thylakoid part)
84 GO:0005622 (intracellular anatomical structure)
6 GO:0009535 (chloroplast thylakoid membrane)
7 GO:0009534 (chloroplast thylakoid)
7 GO:0031976 (plastid thylakoid)
6 GO:0055035 (plastid thylakoid membrane)
76 GO:0043229 (intracellular organelle)
9 GO:0005730 (nucleolus)
76 GO:0043226 (organelle)
12 GO:1902494 (catalytic complex)
8 GO:0140535 (intracellular protein-containing complex)
30 GO:0051276 (chromosome organization)
26 GO:0006325 (chromatin organization)
13 GO:0034728 (nucleosome organization)
12 GO:0006334 (nucleosome assembly)
22 GO:0006338 (chromatin remodeling)
15 GO:0071103 (DNA conformation change)
14 GO:0071824 (protein-DNA complex subunit organization)
13 GO:0065004 (protein-DNA complex assembly)
39 GO:0006996 (organelle organization)
22 GO:0007049 (cell cycle)
15 GO:0051726 (regulation of cell cycle)
10 GO:0031507 (heterochromatin formation)
10 GO:0070828 (heterochromatin organization)
10 GO:0045814 (negative regulation of gene expression, epigenetic)
10 GO:0007346 (regulation of mitotic cell cycle)
44 GO:0016043 (cellular component organization)
14 GO:0000278 (mitotic cell cycle)
18 GO:0006259 (DNA metabolic process)
16 GO:0022402 (cell cycle process)
10 GO:0016458 (obsolete gene silencing)
11 GO:0040029 (epigenetic regulation of gene expression)
9 GO:0006260 (DNA replication)
44 GO:0071840 (cellular component organization or biogenesis)
10 GO:0000280 (nuclear division)
7 GO:0007018 (microtubule-based movement)
8 GO:0010564 (regulation of cell cycle process)
10 GO:0048285 (organelle fission)
10 GO:1903047 (mitotic cell cycle process)
16 GO:0043933 (protein-containing complex organization)
15 GO:0065003 (protein-containing complex assembly)
5 GO:0000819 (sister chromatid segregation)
10 GO:1902679 (negative regulation of RNA biosynthetic process)
10 GO:1903507 (negative regulation of nucleic acid-templated transcription)
5 GO:2001251 (negative regulation of chromosome organization)
7 GO:1903046 (meiotic cell cycle process)
7 GO:0006928 (obsolete movement of cell or subcellular component)
6 GO:0140013 (meiotic nuclear division)
7 GO:0051321 (meiotic cell cycle)
5 GO:1901987 (regulation of cell cycle phase transition)
30 GO:0034654 (nucleobase-containing compound biosynthetic process)
17 GO:0022607 (cellular component assembly)
32 GO:0080090 (regulation of primary metabolic process)
10 GO:0051253 (negative regulation of RNA metabolic process)
9 GO:0045892 (negative regulation of DNA-templated transcription)
28 GO:0019219 (regulation of nucleobase-containing compound metabolic process)
11 GO:0031327 (negative regulation of cellular biosynthetic process)
5 GO:0140014 (mitotic nuclear division)
11 GO:0009890 (negative regulation of biosynthetic process)
8 GO:0006281 (DNA repair)
10 GO:0045934 (negative regulation of nucleobase-containing compound metabolic process)
6 GO:0033044 (regulation of chromosome organization)
32 GO:0060255 (regulation of macromolecule metabolic process)
7 GO:0007017 (microtubule-based process)
12 GO:0051172 (negative regulation of nitrogen compound metabolic process)
14 GO:0010605 (negative regulation of macromolecule metabolic process)
7 GO:0031399 (regulation of protein modification process)
27 GO:0010556 (regulation of macromolecule biosynthetic process)
25 GO:2001141 (regulation of RNA biosynthetic process)
25 GO:1903506 (regulation of nucleic acid-templated transcription)
34 GO:0019222 (regulation of metabolic process)
30 GO:0051171 (regulation of nitrogen compound metabolic process)
5 GO:0044770 (cell cycle phase transition)
10 GO:0065009 (regulation of molecular function)
10 GO:0010558 (negative regulation of macromolecule biosynthetic process)
5 GO:0006261 (DNA-templated DNA replication)
12 GO:0031324 (negative regulation of cellular metabolic process)
5 GO:0010639 (negative regulation of organelle organization)
26 GO:2000112 (regulation of cellular macromolecule biosynthetic process)
5 GO:0051338 (regulation of transferase activity)
6 GO:0008283 (cell population proliferation)
39 GO:0090304 (nucleic acid metabolic process)
30 GO:0018130 (heterocycle biosynthetic process)
25 GO:0097659 (nucleic acid-templated transcription)
24 GO:0006355 (regulation of DNA-templated transcription)
25 GO:0032774 (RNA biosynthetic process)
14 GO:0009892 (negative regulation of metabolic process)
5 GO:0098813 (nuclear chromosome segregation)
27 GO:0031326 (regulation of cellular biosynthetic process)
27 GO:0009889 (regulation of biosynthetic process)
30 GO:0019438 (aromatic compound biosynthetic process)
6 GO:0043085 (positive regulation of catalytic activity)
25 GO:0051252 (regulation of RNA metabolic process)
17 GO:0048523 (negative regulation of cellular process)
33 GO:0009059 (macromolecule biosynthetic process)
24 GO:0006351 (DNA-templated transcription)
5 GO:0031401 (positive regulation of protein modification process)
5 GO:0007059 (chromosome segregation)
32 GO:0034645 (cellular macromolecule biosynthetic process)
5 GO:0051052 (regulation of DNA metabolic process)
8 GO:0006974 (cellular response to DNA damage stimulus)
9 GO:2000113 (negative regulation of cellular macromolecule biosynthetic process)
27 GO:0010468 (regulation of gene expression)
5 GO:0051129 (negative regulation of cellular component organization)
5 GO:0006310 (DNA recombination)
11 GO:0010629 (negative regulation of gene expression)
41 GO:0006139 (nucleobase-containing compound metabolic process)
30 GO:0031323 (regulation of cellular metabolic process)
6 GO:0044093 (positive regulation of molecular function)
20 GO:0048519 (negative regulation of biological process)
43 GO:0050794 (regulation of cellular process)
30 GO:1901362 (organic cyclic compound biosynthetic process)
7 GO:0033043 (regulation of organelle organization)
46 GO:0050789 (regulation of biological process)
17 GO:0044085 (cellular component biogenesis)
7 GO:0050790 (regulation of catalytic activity)
30 GO:0044271 (cellular nitrogen compound biosynthetic process)
5 GO:1901657 (glycosyl compound metabolic process)
41 GO:0046483 (heterocycle metabolic process)
49 GO:0065007 (biological regulation)
42 GO:0034641 (cellular nitrogen compound metabolic process)
41 GO:0006725 (cellular aromatic compound metabolic process)
42 GO:1901360 (organic cyclic compound metabolic process)
5 GO:0051247 (positive regulation of protein metabolic process)
7 GO:0032268 (regulation of cellular protein metabolic process)
0 5 10 15 20 25 30 35
-log(p_value)
Class
Molecular function
Cellular component
Biological process

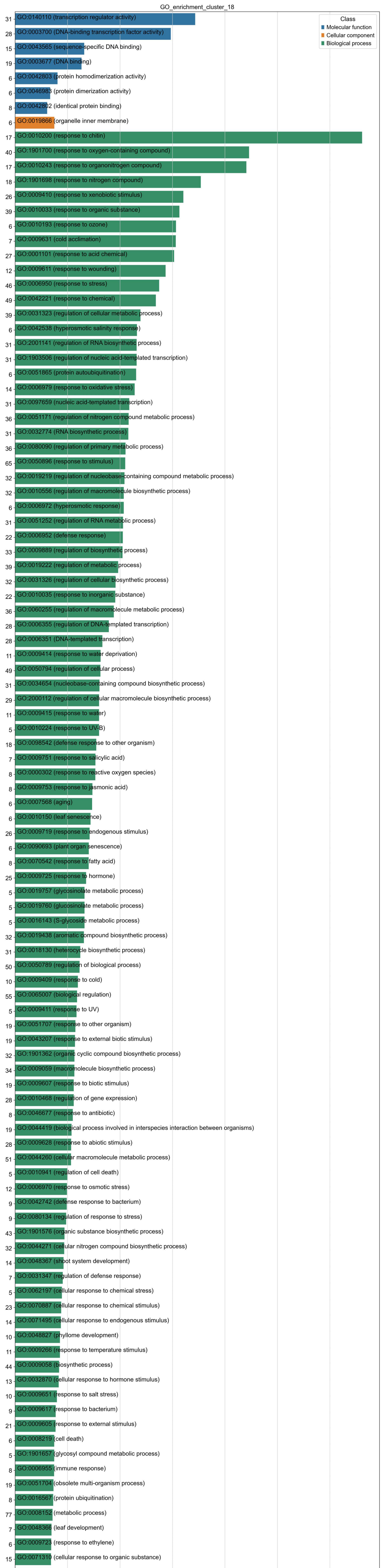
GO_enrichment_cluster_18
Class
Molecular function
Cellular component
Biological process
31 GO:0140110 (transcription regulator activity)
28 GO:0003700 (DNA-binding transcription factor activity)
15 GO:0043565 (sequence-specific DNA binding)
19 GO:0003677 (DNA binding)
6 GO:0042803 (protein homodimerization activity)
6 GO:0046983 (protein dimerization activity)
8 GO:0042802 (identical protein binding)
6 GO:0019866 (organelle inner membrane)
17 GO:0010200 (response to chitin)
40 GO:1901700 (response to oxygen-containing compound)
17 GO:0010243 (response to organonitrogen compound)
18 GO:1901698 (response to nitrogen compound)
26 GO:0009410 (response to xenobiotic stimulus)
39 GO:0010033 (response to organic substance)
6 GO:0010193 (response to ozone)
7 GO:0009631 (cold acclimation)
27 GO:0001101 (response to acid chemical)
12 GO:0009611 (response to wounding)
46 GO:0006950 (response to stress)
49 GO:0042221 (response to chemical)
39 GO:0031323 (regulation of cellular metabolic process)
6 GO:0042538 (hyperosmotic salinity response)
31 GO:2001141 (regulation of RNA biosynthetic process)
31 GO:1903506 (regulation of nucleic acid-templated transcription)
6 GO:0051865 (protein autoubiquitination)
14 GO:0006979 (response to oxidative stress)
31 GO:0097659 (nucleic acid-templated transcription)
36 GO:0051171 (regulation of nitrogen compound metabolic process)
31 GO:0032774 (RNA biosynthetic process)
36 GO:0080090 (regulation of primary metabolic process)
65 GO:0050896 (response to stimulus)
32 GO:0019219 (regulation of nucleobase-containing compound metabolic process)
32 GO:0010556 (regulation of macromolecule biosynthetic process)
6 GO:0006972 (hyperosmotic response)
31 GO:0051252 (regulation of RNA metabolic process)
22 GO:0006952 (defense response)
33 GO:0009889 (regulation of biosynthetic process)
39 GO:0019222 (regulation of metabolic process)
32 GO:0031326 (regulation of cellular biosynthetic process)
22 GO:0010035 (response to inorganic substance)
36 GO:0060255 (regulation of macromolecule metabolic process)
28 GO:0006355 (regulation of DNA-templated transcription)
28 GO:0006351 (DNA-templated transcription)
11 GO:0009414 (response to water deprivation)
49 GO:0050794 (regulation of cellular process)
31 GO:0034654 (nucleobase-containing compound biosynthetic process)
29 GO:2000112 (regulation of cellular macromolecule biosynthetic process)
11 GO:0009415 (response to water)
5 GO:0010224 (response to UV-B)
18 GO:0098542 (defense response to other organism)
7 GO:0009751 (response to salicylic acid)
8 GO:0000302 (response to reactive oxygen species)
8 GO:0009753 (response to jasmonic acid)
6 GO:0007568 (aging)
6 GO:0010150 (leaf senescence)
26 GO:0009719 (response to endogenous stimulus)
6 GO:0090693 (plant organ senescence)
8 GO:0070542 (response to fatty acid)
25 GO:0009725 (response to hormone)
5 GO:0019757 (glycosinolate metabolic process)
5 GO:0019760 (glucosinolate metabolic process)
5 GO:0016143 (S-glycoside metabolic process)
32 GO:0019438 (aromatic compound biosynthetic process)
31 GO:0018130 (heterocycle biosynthetic process)
50 GO:0050789 (regulation of biological process)
10 GO:0009409 (response to cold)
55 GO:0065007 (biological regulation)
5 GO:0009411 (response to UV)
19 GO:0051707 (response to other organism)
19 GO:0043207 (response to external biotic stimulus)
32 GO:1901362 (organic cyclic compound biosynthetic process)
34 GO:0009059 (macromolecule biosynthetic process)
19 GO:0009607 (response to biotic stimulus)
28 GO:0010468 (regulation of gene expression)
8 GO:0046677 (response to antibiotic)
19 GO:0044419 (biological process involved in interspecies interaction between organisms)
28 GO:0009628 (response to abiotic stimulus)
51 GO:0044260 (cellular macromolecule metabolic process)
5 GO:0010941 (regulation of cell death)
12 GO:0006970 (response to osmotic stress)
9 GO:0042742 (defense response to bacterium)
9 GO:0080134 (regulation of response to stress)
43 GO:1901576 (organic substance biosynthetic process)
32 GO:0044271 (cellular nitrogen compound biosynthetic process)
14 GO:0048367 (shoot system development)
7 GO:0031347 (regulation of defense response)
5 GO:0062197 (cellular response to chemical stress)
23 GO:0070887 (cellular response to chemical stimulus)
14 GO:0071495 (cellular response to endogenous stimulus)
10 GO:0048827 (phyllome development)
11 GO:0009266 (response to temperature stimulus)
44 GO:0009058 (biosynthetic process)
13 GO:0032870 (cellular response to hormone stimulus)
10 GO:0009651 (response to salt stress)
9 GO:0009617 (response to bacterium)
21 GO:0009605 (response to external stimulus)
6 GO:0008219 (cell death)
5 GO:1901657 (glycosyl compound metabolic process)
8 GO:0006955 (immune response)
19 GO:0051704 (obsolete multi-organism process)
8 GO:0016567 (protein ubiquitination)
77 GO:0008152 (metabolic process)
7 GO:0048366 (leaf development)
6 GO:0009723 (response to ethylene)
15 GO:0071310 (cellular response to organic substance)
8 GO:0032446 (protein modification by small protein conjugation)
11 GO:0009755 (hormone-mediated signaling pathway)
6 GO:0040008 (regulation of growth)
57 GO:0043170 (macromolecule metabolic process)
29 GO:0034645 (cellular macromolecule biosynthetic process)
7 GO:0045087 (innate immune response)
8 GO:0002376 (immune system process)
14 GO:0033993 (response to lipid)
8 GO:1902680 (positive regulation of RNA biosynthetic process)
8 GO:1903508 (positive regulation of nucleic acid-templated transcription)
31 GO:0010467 (gene expression)
6 GO:0009733 (response to auxin)
0
5
10
15
20
25
30
-log(p_value)

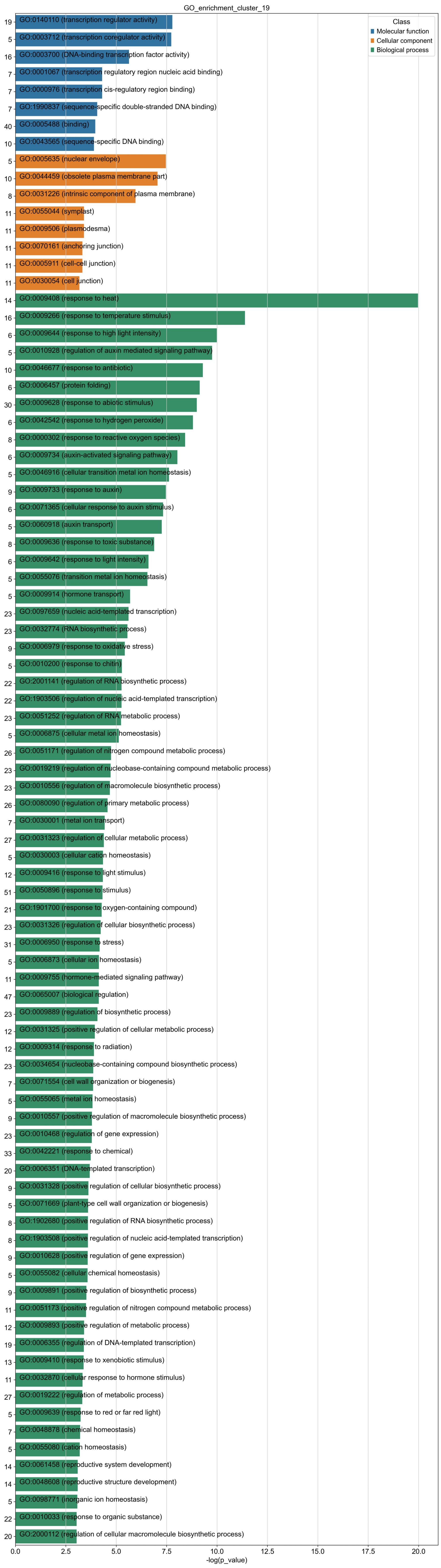
GO_enrichment_cluster_19
Class
Molecular function
Cellular component
Biological process
19 GO:0140110 (transcription regulator activity)
5 GO:0003712 (transcription coregulator activity)
16 GO:0003700 (DNA-binding transcription factor activity)
7 GO:0001067 (transcription regulatory region nucleic acid binding)
7 GO:0000976 (transcription cis-regulatory region binding)
7 GO:1990837 (sequence-specific double-stranded DNA binding)
40 GO:0005488 (binding)
10 GO:0043565 (sequence-specific DNA binding)
5 GO:0005635 (nuclear envelope)
10 GO:0044459 (obsolete plasma membrane part)
8 GO:0031226 (intrinsic component of plasma membrane)
11 GO:0055044 (symplast)
11 GO:0009506 (plasmodesma)
11 GO:0070161 (anchoring junction)
11 GO:0005911 (cell-cell junction)
11 GO:0030054 (cell junction)
14 GO:0009408 (response to heat)
16 GO:0009266 (response to temperature stimulus)
6 GO:0009644 (response to high light intensity)
5 GO:0010928 (regulation of auxin mediated signaling pathway)
10 GO:0046677 (response to antibiotic)
6 GO:0006457 (protein folding)
30 GO:0009628 (response to abiotic stimulus)
6 GO:0042542 (response to hydrogen peroxide)
8 GO:0000302 (response to reactive oxygen species)
6 GO:0009734 (auxin-activated signaling pathway)
5 GO:0046916 (cellular transition metal ion homeostasis)
9 GO:0009733 (response to auxin)
6 GO:0071365 (cellular response to auxin stimulus)
5 GO:0060918 (auxin transport)
8 GO:0009636 (response to toxic substance)
6 GO:0009642 (response to light intensity)
5 GO:0055076 (transition metal ion homeostasis)
5 GO:0009914 (hormone transport)
23 GO:0097659 (nucleic acid-templated transcription)
23 GO:0032774 (RNA biosynthetic process)
9 GO:0006979 (response to oxidative stress)
5 GO:0010200 (response to chitin)
22 GO:2001141 (regulation of RNA biosynthetic process)
22 GO:1903506 (regulation of nucleic acid-templated transcription)
23 GO:0051252 (regulation of RNA metabolic process)
5 GO:0006875 (cellular metal ion homeostasis)
26 GO:0051171 (regulation of nitrogen compound metabolic process)
23 GO:0019219 (regulation of nucleobase-containing compound metabolic process)
23 GO:0010556 (regulation of macromolecule biosynthetic process)
26 GO:0080090 (regulation of primary metabolic process)
7 GO:0030001 (metal ion transport)
27 GO:0031323 (regulation of cellular metabolic process)
5 GO:0030003 (cellular cation homeostasis)
12 GO:0009416 (response to light stimulus)
51 GO:0050896 (response to stimulus)
21 GO:1901700 (response to oxygen-containing compound)
23 GO:0031326 (regulation of cellular biosynthetic process)
31 GO:0006950 (response to stress)
5 GO:0006873 (cellular ion homeostasis)
11 GO:0009755 (hormone-mediated signaling pathway)
47 GO:0065007 (biological regulation)
23 GO:0009889 (regulation of biosynthetic process)
12 GO:0031325 (positive regulation of cellular metabolic process)
12 GO:0009314 (response to radiation)
23 GO:0034654 (nucleobase-containing compound biosynthetic process)
7 GO:0071554 (cell wall organization or biogenesis)
5 GO:0055065 (metal ion homeostasis)
9 GO:0010557 (positive regulation of macromolecule biosynthetic process)
23 GO:0010468 (regulation of gene expression)
33 GO:0042221 (response to chemical)
20 GO:0006351 (DNA-templated transcription)
9 GO:0031328 (positive regulation of cellular biosynthetic process)
5 GO:0071669 (plant-type cell wall organization or biogenesis)
8 GO:1902680 (positive regulation of RNA biosynthetic process)
8 GO:1903508 (positive regulation of nucleic acid-templated transcription)
9 GO:0010628 (positive regulation of gene expression)
5 GO:0055082 (cellular chemical homeostasis)
9 GO:0009891 (positive regulation of biosynthetic process)
11 GO:0051173 (positive regulation of nitrogen compound metabolic process)
12 GO:0009893 (positive regulation of metabolic process)
19 GO:0006355 (regulation of DNA-templated transcription)
13 GO:0009410 (response to xenobiotic stimulus)
11 GO:0032870 (cellular response to hormone stimulus)
27 GO:0019222 (regulation of metabolic process)
5 GO:0009639 (response to red or far red light)
7 GO:0048878 (chemical homeostasis)
5 GO:0055080 (cation homeostasis)
14 GO:0061458 (reproductive system development)
14 GO:0048608 (reproductive structure development)
5 GO:0098771 (inorganic ion homeostasis)
22 GO:0010033 (response to organic substance)
20 GO:2000112 (regulation of cellular macromolecule biosynthetic process)
0.0
2.5
5.0
7.5
10.0
12.5
15.0
17.5
20.0
-log(p_value)

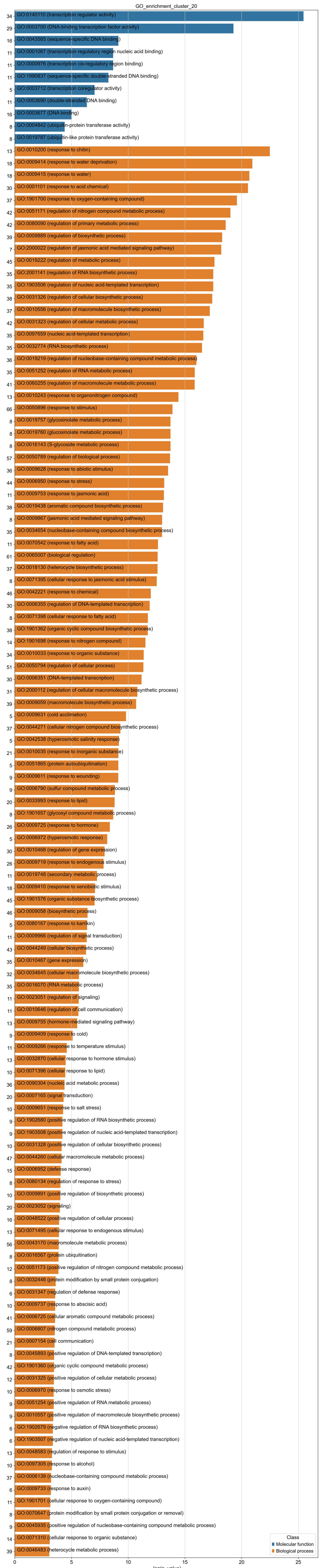
GO_enrichment_cluster_20
34 GO:0140110 (transcription regulator activity)
29 GO:0003700 (DNA-binding transcription factor activity)
16 GO:0043565 (sequence-specific DNA binding)
11 GO:0001067 (transcription regulatory region nucleic acid binding)
11 GO:0000976 (transcription cis-regulatory region binding)
11 GO:1990837 (sequence-specific double-stranded DNA binding)
5 GO:0003712 (transcription coregulator activity)
11 GO:0003690 (double-stranded DNA binding)
16 GO:0003677 (DNA binding)
8 GO:0004842 (ubiquitin-protein transferase activity)
8 GO:0019787 (ubiquitin-like protein transferase activity)
13 GO:0010200 (response to chitin)
18 GO:0009414 (response to water deprivation)
18 GO:0009415 (response to water)
30 GO:0001101 (response to acid chemical)
37 GO:1901700 (response to oxygen-containing compound)
42 GO:0051171 (regulation of nitrogen compound metabolic process)
42 GO:0080090 (regulation of primary metabolic process)
39 GO:0009889 (regulation of biosynthetic process)
7 GO:2000022 (regulation of jasmonic acid mediated signaling pathway)
45 GO:0019222 (regulation of metabolic process)
35 GO:2001141 (regulation of RNA biosynthetic process)
35 GO:1903506 (regulation of nucleic acid-templated transcription)
38 GO:0031326 (regulation of cellular biosynthetic process)
37 GO:0010556 (regulation of macromolecule biosynthetic process)
42 GO:0031323 (regulation of cellular metabolic process)
35 GO:0097659 (nucleic acid-templated transcription)
35 GO:0032774 (RNA biosynthetic process)
36 GO:0019219 (regulation of nucleobase-containing compound metabolic process)
35 GO:0051252 (regulation of RNA metabolic process)
41 GO:0060255 (regulation of macromolecule metabolic process)
13 GO:0010243 (response to organonitrogen compound)
66 GO:0050896 (response to stimulus)
8 GO:0019757 (glycosinolate metabolic process)
8 GO:0019760 (glucosinolate metabolic process)
8 GO:0016143 (S-glycoside metabolic process)
57 GO:0050789 (regulation of biological process)
36 GO:0009628 (response to abiotic stimulus)
44 GO:0006950 (response to stress)
11 GO:0009753 (response to jasmonic acid)
38 GO:0019438 (aromatic compound biosynthetic process)
8 GO:0009867 (jasmonic acid mediated signaling pathway)
35 GO:0034654 (nucleobase-containing compound biosynthetic process)
11 GO:0070542 (response to fatty acid)
61 GO:0065007 (biological regulation)
37 GO:0018130 (heterocycle biosynthetic process)
8 GO:0071395 (cellular response to jasmonic acid stimulus)
46 GO:0042221 (response to chemical)
30 GO:0006355 (regulation of DNA-templated transcription)
8 GO:0071398 (cellular response to fatty acid)
38 GO:1901362 (organic cyclic compound biosynthetic process)
14 GO:1901698 (response to nitrogen compound)
34 GO:0010033 (response to organic substance)
51 GO:0050794 (regulation of cellular process)
30 GO:0006351 (DNA-templated transcription)
31 GO:2000112 (regulation of cellular macromolecule biosynthetic process)
39 GO:0009059 (macromolecule biosynthetic process)
5 GO:0009631 (cold acclimation)
37 GO:0044271 (cellular nitrogen compound biosynthetic process)
5 GO:0042538 (hyperosmotic salinity response)
21 GO:0010035 (response to inorganic substance)
5 GO:0051865 (protein autoubiquitination)
9 GO:0009611 (response to wounding)
9 GO:0006790 (sulfur compound metabolic process)
20 GO:0033993 (response to lipid)
8 GO:1901657 (glycosyl compound metabolic process)
26 GO:0009725 (response to hormone)
5 GO:0006972 (hyperosmotic response)
30 GO:0010468 (regulation of gene expression)
26 GO:0009719 (response to endogenous stimulus)
11 GO:0019748 (secondary metabolic process)
18 GO:0009410 (response to xenobiotic stimulus)
45 GO:1901576 (organic substance biosynthetic process)
46 GO:0009058 (biosynthetic process)
5 GO:0080167 (response to karrikin)
11 GO:0009966 (regulation of signal transduction)
43 GO:0044249 (cellular biosynthetic process)
35 GO:0010467 (gene expression)
32 GO:0034645 (cellular macromolecule biosynthetic process)
35 GO:0016070 (RNA metabolic process)
11 GO:0023051 (regulation of signaling)
11 GO:0010646 (regulation of cell communication)
13 GO:0009755 (hormone-mediated signaling pathway)
9 GO:0009409 (response to cold)
11 GO:0009266 (response to temperature stimulus)
13 GO:0032870 (cellular response to hormone stimulus)
10 GO:0071396 (cellular response to lipid)
36 GO:0090304 (nucleic acid metabolic process)
20 GO:0007165 (signal transduction)
10 GO:0009651 (response to salt stress)
9 GO:1902680 (positive regulation of RNA biosynthetic process)
9 GO:1903508 (positive regulation of nucleic acid-templated transcription)
10 GO:0031328 (positive regulation of cellular biosynthetic process)
47 GO:0044260 (cellular macromolecule metabolic process)
15 GO:0006952 (defense response)
8 GO:0080134 (regulation of response to stress)
10 GO:0009891 (positive regulation of biosynthetic process)
20 GO:0023052 (signaling)
16 GO:0048522 (positive regulation of cellular process)
13 GO:0071495 (cellular response to endogenous stimulus)
56 GO:0043170 (macromolecule metabolic process)
8 GO:0016567 (protein ubiquitination)
12 GO:0051173 (positive regulation of nitrogen compound metabolic process)
8 GO:0032446 (protein modification by small protein conjugation)
6 GO:0031347 (regulation of defense response)
10 GO:0009737 (response to abscisic acid)
41 GO:0006725 (cellular aromatic compound metabolic process)
59 GO:0006807 (nitrogen compound metabolic process)
21 GO:0007154 (cell communication)
8 GO:0045893 (positive regulation of DNA-templated transcription)
42 GO:1901360 (organic cyclic compound metabolic process)
12 GO:0031325 (positive regulation of cellular metabolic process)
10 GO:0006970 (response to osmotic stress)
9 GO:0051254 (positive regulation of RNA metabolic process)
9 GO:0010557 (positive regulation of macromolecule biosynthetic process)
6 GO:1902679 (negative regulation of RNA biosynthetic process)
6 GO:1903507 (negative regulation of nucleic acid-templated transcription)
13 GO:0048583 (regulation of response to stimulus)
10 GO:0097305 (response to alcohol)
37 GO:0006139 (nucleobase-containing compound metabolic process)
6 GO:0009733 (response to auxin)
11 GO:1901701 (cellular response to oxygen-containing compound)
8 GO:0070647 (protein modification by small protein conjugation or removal)
9 GO:0045935 (positive regulation of nucleobase-containing compound metabolic process)
14 GO:0071310 (cellular response to organic substance)
39 GO:0046483 (heterocycle metabolic process)
0
5
10
15
20
25
-log(p_value)
Class
Molecular function
Biological process

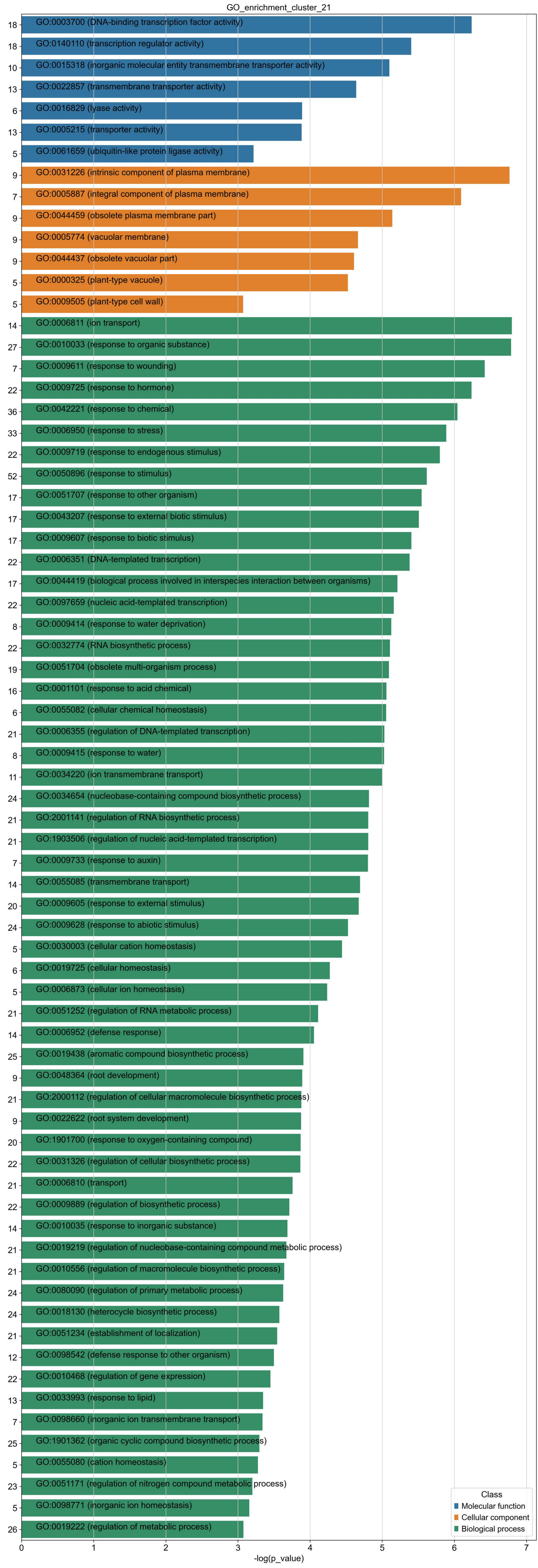
GO_enrichment_cluster_21
18 GO:0003700 (DNA-binding transcription factor activity)
18 GO:0140110 (transcription regulator activity)
10 GO:0015318 (inorganic molecular entity transmembrane transporter activity)
13 GO:0022857 (transmembrane transporter activity)
6 GO:0016829 (lyase activity)
13 GO:0005215 (transporter activity)
5 GO:0061659 (ubiquitin-like protein ligase activity)
9 GO:0031226 (intrinsic component of plasma membrane)
7 GO:0005887 (integral component of plasma membrane)
9 GO:0044459 (obsolete plasma membrane part)
9 GO:0005774 (vacuolar membrane)
9 GO:0044437 (obsolete vacuolar part)
5 GO:0000325 (plant-type vacuole)
5 GO:0009505 (plant-type cell wall)
14 GO:0006811 (ion transport)
27 GO:0010033 (response to organic substance)
7 GO:0009611 (response to wounding)
22 GO:0009725 (response to hormone)
36 GO:0042221 (response to chemical)
33 GO:0006950 (response to stress)
22 GO:0009719 (response to endogenous stimulus)
52 GO:0050896 (response to stimulus)
17 GO:0051707 (response to other organism)
17 GO:0043207 (response to external biotic stimulus)
17 GO:0009607 (response to biotic stimulus)
22 GO:0006351 (DNA-templated transcription)
17 GO:0044419 (biological process involved in interspecies interaction between organisms)
22 GO:0097659 (nucleic acid-templated transcription)
8 GO:0009414 (response to water deprivation)
22 GO:0032774 (RNA biosynthetic process)
19 GO:0051704 (obsolete multi-organism process)
16 GO:0001101 (response to acid chemical)
6 GO:0055082 (cellular chemical homeostasis)
21 GO:0006355 (regulation of DNA-templated transcription)
8 GO:0009415 (response to water)
11 GO:0034220 (ion transmembrane transport)
24 GO:0034654 (nucleobase-containing compound biosynthetic process)
21 GO:2001141 (regulation of RNA biosynthetic process)
21 GO:1903506 (regulation of nucleic acid-templated transcription)
7 GO:0009733 (response to auxin)
14 GO:0055085 (transmembrane transport)
20 GO:0009605 (response to external stimulus)
24 GO:0009628 (response to abiotic stimulus)
5 GO:0030003 (cellular cation homeostasis)
6 GO:0019725 (cellular homeostasis)
5 GO:0006873 (cellular ion homeostasis)
21 GO:0051252 (regulation of RNA metabolic process)
14 GO:0006952 (defense response)
25 GO:0019438 (aromatic compound biosynthetic process)
9 GO:0048364 (root development)
21 GO:2000112 (regulation of cellular macromolecule biosynthetic process)
9 GO:0022622 (root system development)
20 GO:1901700 (response to oxygen-containing compound)
22 GO:0031326 (regulation of cellular biosynthetic process)
21 GO:0006810 (transport)
22 GO:0009889 (regulation of biosynthetic process)
14 GO:0010035 (response to inorganic substance)
21 GO:0019219 (regulation of nucleobase-containing compound metabolic process)
21 GO:0010556 (regulation of macromolecule biosynthetic process)
24 GO:0080090 (regulation of primary metabolic process)
24 GO:0018130 (heterocycle biosynthetic process)
21 GO:0051234 (establishment of localization)
12 GO:0098542 (defense response to other organism)
22 GO:0010468 (regulation of gene expression)
13 GO:0033993 (response to lipid)
7 GO:0098660 (inorganic ion transmembrane transport)
25 GO:1901362 (organic cyclic compound biosynthetic process)
5 GO:0055080 (cation homeostasis)
23 GO:0051171 (regulation of nitrogen compound metabolic process)
5 GO:0098771 (inorganic ion homeostasis)
26 GO:0019222 (regulation of metabolic process)
0 1 2 3 4 5 6 7
-log(p_value)
Class
Molecular function
Cellular component
Biological process

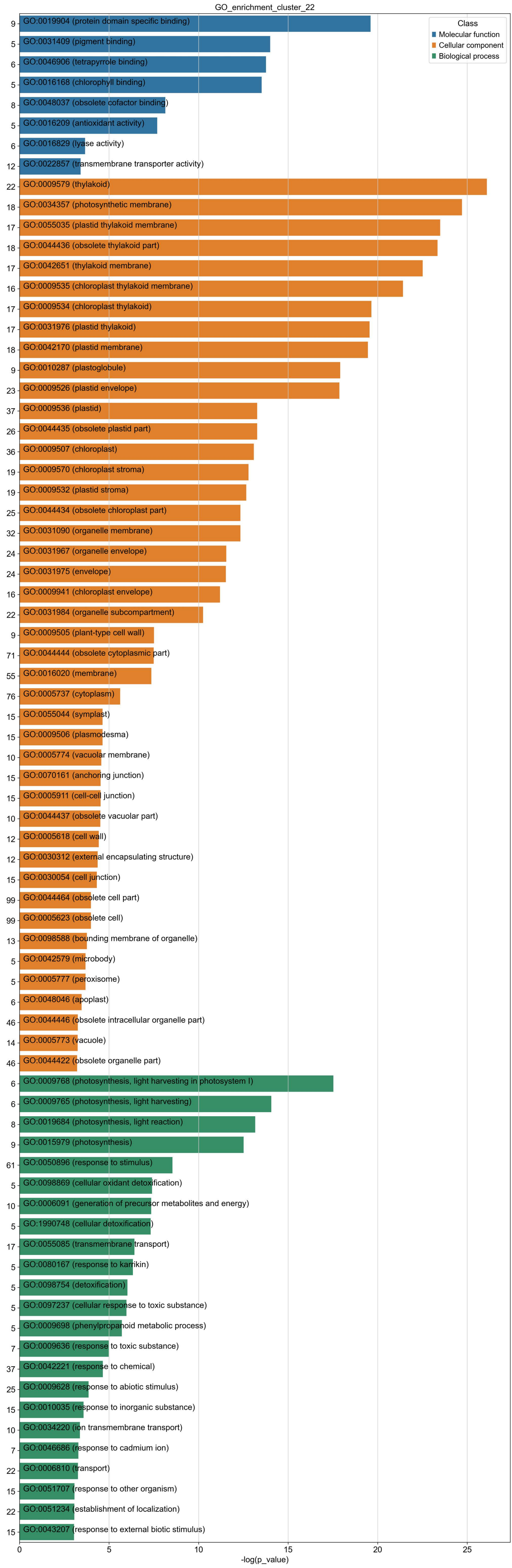

GO_enrichment_cluster_22
Class
Molecular function
Cellular component
Biological process
GO:0019904 (protein domain specific binding)
GO:0031409 (pigment binding)
GO:0046906 (tetrapyrrole binding)
GO:0016168 (chlorophyll binding)
GO:0048037 (obsolete cofactor binding)
GO:0016209 (antioxidant activity)
GO:0016829 (lyase activity)
GO:0022857 (transmembrane transporter activity)
GO:0009579 (thylakoid)
GO:0034357 (photosynthetic membrane)
GO:0055035 (plastid thylakoid membrane)
GO:0044436 (obsolete thylakoid part)
GO:0042651 (thylakoid membrane)
GO:0009535 (chloroplast thylakoid membrane)
GO:0009534 (chloroplast thylakoid)
GO:0031976 (plastid thylakoid)
GO:0042170 (plastid membrane)
GO:0010287 (plastoglobule)
GO:0009526 (plastid envelope)
GO:0009536 (plastid)
GO:0044435 (obsolete plastid part)
GO:0009507 (chloroplast)
GO:0009570 (chloroplast stroma)
GO:0009532 (plastid stroma)
GO:0044434 (obsolete chloroplast part)
GO:0031090 (organelle membrane)
GO:0031967 (organelle envelope)
GO:0031975 (envelope)
GO:0009941 (chloroplast envelope)
GO:0031984 (organelle subcompartment)
GO:0009505 (plant-type cell wall)
GO:0044444 (obsolete cytoplasmic part)
GO:0016020 (membrane)
GO:0005737 (cytoplasm)
GO:0055044 (symplast)
GO:0009506 (plasmodesma)
GO:0005774 (vacuolar membrane)
GO:0070161 (anchoring junction)
GO:0005911 (cell-cell junction)
GO:0044437 (obsolete vacuolar part)
GO:0005618 (cell wall)
GO:0030312 (external encapsulating structure)
GO:0030054 (cell junction)
GO:0044464 (obsolete cell part)
GO:0005623 (obsolete cell)
GO:0098588 (bounding membrane of organelle)
GO:0042579 (microbody)
GO:0005777 (peroxisome)
GO:0048046 (apoplast)
GO:0044446 (obsolete intracellular organelle part)
GO:0005773 (vacuole)
GO:0044422 (obsolete organelle part)
GO:0009768 (photosynthesis, light harvesting in photosystem I)
GO:0009765 (photosynthesis, light harvesting)
GO:0019684 (photosynthesis, light reaction)
GO:0015979 (photosynthesis)
GO:0050896 (response to stimulus)
GO:0098869 (cellular oxidant detoxification)
GO:0006091 (generation of precursor metabolites and energy)
GO:1990748 (cellular detoxification)
GO:0055085 (transmembrane transport)
GO:0080167 (response to karrikin)
GO:0098754 (detoxification)
GO:0097237 (cellular response to toxic substance)
GO:0009698 (phenylpropanoid metabolic process)
GO:0009636 (response to toxic substance)
GO:0042221 (response to chemical)
GO:0009628 (response to abiotic stimulus)
GO:0010035 (response to inorganic substance)
GO:0034220 (ion transmembrane transport)
GO:0046686 (response to cadmium ion)
GO:0006810 (transport)
GO:0051707 (response to other organism)
GO:0051234 (establishment of localization)
GO:0043207 (response to external biotic stimulus)
9
5
6
5
8
5
6
12
22
18
17
18
17
16
17
17
18
9
23
37
26
36
19
19
25
32
24
24
16
22
9
71
55
76
15
15
10
15
15
10
12
12
15
99
99
13
5
5
6
46
14
46
6
6
8
9
61
5
10
5
17
5
5
5
5
7
37
25
15
10
7
22
15
22
15
0
5
10
15
20
25
-log(p_value)

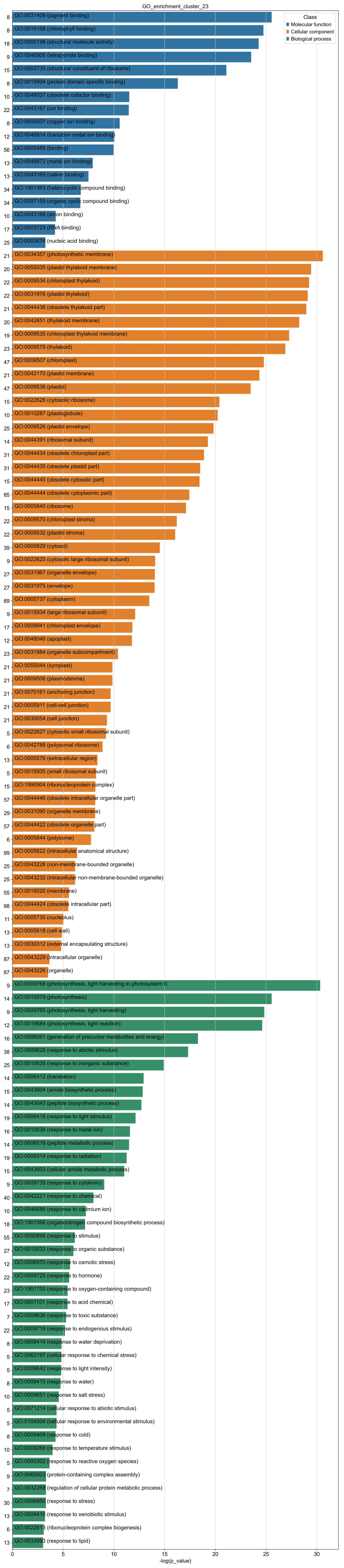
GO_enrichment_cluster_23
Class
Molecular function
Cellular component
Biological process
8 GO:0031409 (pigment binding)
8 GO:0016168 (chlorophyll binding)
18 GO:0005198 (structural molecule activity)
9 GO:0046906 (tetrapyrrole binding)
15 GO:0003735 (structural constituent of ribosome)
8 GO:0019904 (protein domain specific binding)
10 GO:0048037 (obsolete cofactor binding)
22 GO:0043167 (ion binding)
8 GO:0005507 (copper ion binding)
12 GO:0046914 (transition metal ion binding)
56 GO:0005488 (binding)
13 GO:0046872 (metal ion binding)
13 GO:0043169 (cation binding)
34 GO:1901363 (heterocyclic compound binding)
34 GO:0097159 (organic cyclic compound binding)
10 GO:0043168 (anion binding)
17 GO:0003723 (RNA binding)
25 GO:0003676 (nucleic acid binding)
21 GO:0034357 (photosynthetic membrane)
20 GO:0055035 (plastid thylakoid membrane)
22 GO:0009534 (chloroplast thylakoid)
22 GO:0031976 (plastid thylakoid)
21 GO:0044436 (obsolete thylakoid part)
20 GO:0042651 (thylakoid membrane)
19 GO:0009535 (chloroplast thylakoid membrane)
23 GO:0009579 (thylakoid)
47 GO:0009507 (chloroplast)
21 GO:0042170 (plastid membrane)
47 GO:0009536 (plastid)
15 GO:0022626 (cytosolic ribosome)
10 GO:0010287 (plastoglobule)
25 GO:0009526 (plastid envelope)
14 GO:0044391 (ribosomal subunit)
31 GO:0044434 (obsolete chloroplast part)
31 GO:0044435 (obsolete plastid part)
15 GO:0044445 (obsolete cytosolic part)
85 GO:0044444 (obsolete cytoplasmic part)
15 GO:0005840 (ribosome)
22 GO:0009570 (chloroplast stroma)
22 GO:0009532 (plastid stroma)
39 GO:0005829 (cytosol)
9 GO:0022625 (cytosolic large ribosomal subunit)
27 GO:0031967 (organelle envelope)
27 GO:0031975 (envelope)
89 GO:0005737 (cytoplasm)
9 GO:0015934 (large ribosomal subunit)
17 GO:0009941 (chloroplast envelope)
12 GO:0048046 (apoplast)
23 GO:0031984 (organelle subcompartment)
21 GO:0055044 (symplast)
21 GO:0009506 (plasmodesma)
21 GO:0070161 (anchoring junction)
21 GO:0005911 (cell-cell junction)
21 GO:0030054 (cell junction)
5 GO:0022627 (cytosolic small ribosomal subunit)
6 GO:0042788 (polysomal ribosome)
13 GO:0005576 (extracellular region)
5 GO:0015935 (small ribosomal subunit)
15 GO:1990904 (ribonucleoprotein complex)
57 GO:0044446 (obsolete intracellular organelle part)
29 GO:0031090 (organelle membrane)
57 GO:0044422 (obsolete organelle part)
6 GO:0005844 (polysome)
99 GO:0005622 (intracellular anatomical structure)
25 GO:0043228 (non-membrane-bounded organelle)
25 GO:0043232 (intracellular non-membrane-bounded organelle)
55 GO:0016020 (membrane)
98 GO:0044424 (obsolete intracellular part)
11 GO:0005730 (nucleolus)
13 GO:0005618 (cell wall)
13 GO:0030312 (external encapsulating structure)
87 GO:0043229 (intracellular organelle)
87 GO:0043226 (organelle)
9 GO:0009768 (photosynthesis, light harvesting in photosystem I)
14 GO:0015979 (photosynthesis)
9 GO:0009765 (photosynthesis, light harvesting)
12 GO:0019684 (photosynthesis, light reaction)
16 GO:0006091 (generation of precursor metabolites and energy)
38 GO:0009628 (response to abiotic stimulus)
25 GO:0010035 (response to inorganic substance)
14 GO:0006412 (translation)
15 GO:0043604 (amide biosynthetic process)
14 GO:0043043 (peptide biosynthetic process)
19 GO:0009416 (response to light stimulus)
16 GO:0010038 (response to metal ion)
14 GO:0006518 (peptide metabolic process)
19 GO:0009314 (response to radiation)
15 GO:0043603 (cellular amide metabolic process)
9 GO:0009735 (response to cytokinin)
40 GO:0042221 (response to chemical)
10 GO:0046686 (response to cadmium ion)
18 GO:1901566 (organonitrogen compound biosynthetic process)
55 GO:0050896 (response to stimulus)
27 GO:0010033 (response to organic substance)
12 GO:0006970 (response to osmotic stress)
22 GO:0009725 (response to hormone)
23 GO:1901700 (response to oxygen-containing compound)
17 GO:0001101 (response to acid chemical)
7 GO:0009636 (response to toxic substance)
22 GO:0009719 (response to endogenous stimulus)
8 GO:0009414 (response to water deprivation)
5 GO:0062197 (cellular response to chemical stress)
5 GO:0009642 (response to light intensity)
8 GO:0009415 (response to water)
10 GO:0009651 (response to salt stress)
5 GO:0071214 (cellular response to abiotic stimulus)
5 GO:0104004 (cellular response to environmental stimulus)
8 GO:0009409 (response to cold)
10 GO:0009266 (response to temperature stimulus)
5 GO:0000302 (response to reactive oxygen species)
9 GO:0065003 (protein-containing complex assembly)
7 GO:0032268 (regulation of cellular protein metabolic process)
30 GO:0006950 (response to stress)
13 GO:0009410 (response to xenobiotic stimulus)
6 GO:0022613 (ribonucleoprotein complex biogenesis)
13 GO:0033993 (response to lipid)
0
5
10
15
20
25
30
-log(p_value)

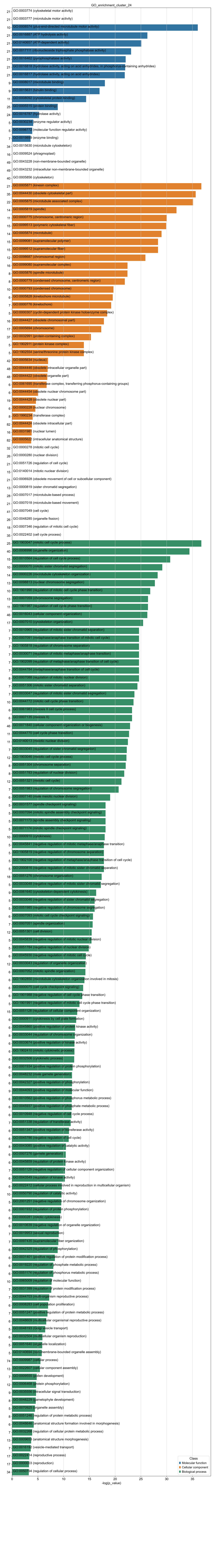
GO_enrichment_cluster_24
21 GO:0003774 (cytoskeletal motor activity)
21 GO:0003777 (microtubule motor activity)
10 GO:0008574 (plus-end-directed microtubule motor activity)
21 GO:0016887 (ATP hydrolysis activity)
21 GO:0140657 (ATP-dependent activity)
21 GO:0017111 (ribonucleoside triphosphate phosphatase activity)
21 GO:0016462 (pyrophosphatase activity)
21 GO:0016818 (hydrolase activity, acting on acid anhydrides, in phosphorus-containing anhydrides)
21 GO:0016817 (hydrolase activity, acting on acid anhydrides)
9 GO:0008017 (microtubule binding)
9 GO:0015631 (tubulin binding)
11 GO:0008092 (cytoskeletal protein binding)
25 GO:0005515 (protein binding)
24 GO:0016787 (hydrolase activity)
5 GO:0030234 (enzyme regulator activity)
5 GO:0098772 (molecular function regulator activity)
7 GO:0019899 (enzyme binding)
34 GO:0015630 (microtubule cytoskeleton)
16 GO:0009524 (phragmoplast)
49 GO:0043228 (non-membrane-bounded organelle)
49 GO:0043232 (intracellular non-membrane-bounded organelle)
40 GO:0005856 (cytoskeleton)
21 GO:0005871 (kinesin complex)
35 GO:0044430 (obsolete cytoskeletal part)
22 GO:0005875 (microtubule associated complex)
14 GO:0005819 (spindle)
11 GO:0000775 (chromosome, centromeric region)
15 GO:0099513 (polymeric cytoskeletal fiber)
14 GO:0005874 (microtubule)
15 GO:0099081 (supramolecular polymer)
15 GO:0099512 (supramolecular fiber)
12 GO:0098687 (chromosomal region)
16 GO:0099080 (supramolecular complex)
8 GO:0005876 (spindle microtubule)
8 GO:0000779 (condensed chromosome, centromeric region)
10 GO:0000793 (condensed chromosome)
6 GO:0005828 (kinetochore microtubule)
7 GO:0000776 (kinetochore)
5 GO:0000307 (cyclin-dependent protein kinase holoenzyme complex)
16 GO:0044427 (obsolete chromosomal part)
17 GO:0005694 (chromosome)
37 GO:0032991 (protein-containing complex)
5 GO:1902911 (protein kinase complex)
5 GO:1902554 (serine/threonine protein kinase complex)
42 GO:0005634 (nucleus)
48 GO:0044446 (obsolete intracellular organelle part)
48 GO:0044422 (obsolete organelle part)
6 GO:0061695 (transferase complex, transferring phosphorus-containing groups)
6 GO:0044454 (obsolete nuclear chromosome part)
19 GO:0044428 (obsolete nuclear part)
6 GO:0000228 (nuclear chromosome)
8 GO:1990234 (transferase complex)
82 GO:0044424 (obsolete intracellular part)
16 GO:0031981 (nuclear lumen)
82 GO:0005622 (intracellular anatomical structure)
32 GO:0000278 (mitotic cell cycle)
26 GO:0000280 (nuclear division)
21 GO:0051726 (regulation of cell cycle)
15 GO:0140014 (mitotic nuclear division)
21 GO:0006928 (obsolete movement of cell or subcellular component)
13 GO:0000819 (sister chromatid segregation)
28 GO:0007017 (microtubule-based process)
21 GO:0007018 (microtubule-based movement)
41 GO:0007049 (cell cycle)
26 GO:0048285 (organelle fission)
18 GO:0007346 (regulation of mitotic cell cycle)
35 GO:0022402 (cell cycle process)
25 GO:1903047 (mitotic cell cycle process)
40 GO:0006996 (organelle organization)
15 GO:0010564 (regulation of cell cycle process)
10 GO:0000070 (mitotic sister chromatid segregation)
14 GO:0000226 (microtubule cytoskeleton organization)
13 GO:0098813 (nuclear chromosome segregation)
10 GO:1901990 (regulation of mitotic cell cycle phase transition)
13 GO:0007059 (chromosome segregation)
11 GO:1901987 (regulation of cell cycle phase transition)
46 GO:0016043 (cellular component organization)
17 GO:0007010 (cytoskeleton organization)
7 GO:0010965 (regulation of mitotic sister chromatid separation)
7 GO:0007091 (metaphase/anaphase transition of mitotic cell cycle)
7 GO:1905818 (regulation of chromosome separation)
7 GO:0030071 (regulation of mitotic metaphase/anaphase transition)
7 GO:1902099 (regulation of metaphase/anaphase transition of cell cycle)
7 GO:0044784 (metaphase/anaphase transition of cell cycle)
8 GO:0007088 (regulation of mitotic nuclear division)
7 GO:0051306 (mitotic sister chromatid separation)
7 GO:0033047 (regulation of mitotic sister chromatid segregation)
10 GO:0044772 (mitotic cell cycle phase transition)
6 GO:0061983 (meiosis II cell cycle process)
6 GO:0007135 (meiosis II)
46 GO:0071840 (cellular component organization or biogenesis)
11 GO:0044770 (cell cycle phase transition)
11 GO:0140013 (meiotic nuclear division)
7 GO:0033045 (regulation of sister chromatid segregation)
12 GO:1903046 (meiotic cell cycle process)
8 GO:0051304 (chromosome separation)
8 GO:0051783 (regulation of nuclear division)
12 GO:0051321 (meiotic cell cycle)
7 GO:0051983 (regulation of chromosome segregation)
6 GO:0007140 (male meiotic nuclear division)
5 GO:0031577 (spindle checkpoint signaling)
5 GO:0007094 (mitotic spindle assembly checkpoint signaling)
5 GO:0071173 (spindle assembly checkpoint signaling)
5 GO:0071174 (mitotic spindle checkpoint signaling)
10 GO:0000910 (cytokinesis)
5 GO:0045841 (negative regulation of mitotic metaphase/anaphase transition)
5 GO:1905819 (negative regulation of chromosome separation)
5 GO:1902100 (negative regulation of metaphase/anaphase transition of cell cycle)
5 GO:2000816 (negative regulation of mitotic sister chromatid separation)
18 GO:0051276 (chromosome organization)
5 GO:0033048 (negative regulation of mitotic sister chromatid segregation)
9 GO:0061640 (cytoskeleton-dependent cytokinesis)
5 GO:0033046 (negative regulation of sister chromatid segregation)
5 GO:0051985 (negative regulation of chromosome segregation)
6 GO:0007093 (mitotic cell cycle checkpoint signaling)
7 GO:0007051 (spindle organization)
12 GO:0051301 (cell division)
5 GO:0045839 (negative regulation of mitotic nuclear division)
5 GO:0051784 (negative regulation of nuclear division)
6 GO:0045930 (negative regulation of mitotic cell cycle)
12 GO:0033043 (regulation of organelle organization)
6 GO:0007052 (mitotic spindle organization)
6 GO:1902850 (microtubule cytoskeleton organization involved in mitosis)
6 GO:0000075 (cell cycle checkpoint signaling)
6 GO:1901988 (negative regulation of cell cycle phase transition)
5 GO:1901991 (negative regulation of mitotic cell cycle phase transition)
15 GO:0051128 (regulation of cellular component organization)
6 GO:0000911 (cytokinesis by cell plate formation)
6 GO:0045860 (positive regulation of protein kinase activity)
8 GO:0033044 (regulation of chromosome organization)
6 GO:0033674 (positive regulation of kinase activity)
6 GO:1902410 (mitotic cytokinetic process)
6 GO:0032506 (cytokinetic process)
6 GO:0001934 (positive regulation of protein phosphorylation)
6 GO:0048232 (male gamete generation)
6 GO:0042327 (positive regulation of phosphorylation)
9 GO:0044093 (positive regulation of molecular function)
6 GO:0010562 (positive regulation of phosphorus metabolic process)
6 GO:0045937 (positive regulation of phosphate metabolic process)
6 GO:0010948 (negative regulation of cell cycle process)
7 GO:0051338 (regulation of transferase activity)
6 GO:0051347 (positive regulation of transferase activity)
6 GO:0045786 (negative regulation of cell cycle)
8 GO:0043085 (positive regulation of catalytic activity)
6 GO:0007276 (gamete generation)
6 GO:0045859 (regulation of protein kinase activity)
7 GO:0051129 (negative regulation of cellular component organization)
6 GO:0043549 (regulation of kinase activity)
6 GO:0022412 (cellular process involved in reproduction in multicellular organism)
10 GO:0050790 (regulation of catalytic activity)
5 GO:2001251 (negative regulation of chromosome organization)
6 GO:0001932 (regulation of protein phosphorylation)
6 GO:0000281 (mitotic cytokinesis)
6 GO:0010639 (negative regulation of organelle organization)
7 GO:0019953 (sexual reproduction)
7 GO:0097435 (supramolecular fiber organization)
6 GO:0042325 (regulation of phosphorylation)
6 GO:0031401 (positive regulation of protein modification process)
6 GO:0019220 (regulation of phosphate metabolic process)
6 GO:0051174 (regulation of phosphorus metabolic process)
10 GO:0065009 (regulation of molecular function)
7 GO:0031399 (regulation of protein modification process)
7 GO:0044703 (multi-organism reproductive process)
6 GO:0008283 (cell population proliferation)
7 GO:0051247 (positive regulation of protein metabolic process)
6 GO:0048609 (multicellular organismal reproductive process)
5 GO:0048193 (Golgi vesicle transport)
6 GO:0032504 (multicellular organism reproduction)
5 GO:0051640 (organelle localization)
5 GO:0140694 (non-membrane-bounded organelle assembly)
74 GO:0009987 (cellular process)
13 GO:0022607 (cellular component assembly)
7 GO:0009555 (pollen development)
12 GO:0006468 (protein phosphorylation)
9 GO:0035556 (intracellular signal transduction)
8 GO:0048229 (gametophyte development)
5 GO:0070925 (organelle assembly)
8 GO:0051246 (regulation of protein metabolic process)
6 GO:0048646 (anatomical structure formation involved in morphogenesis)
7 GO:0032268 (regulation of cellular protein metabolic process)
13 GO:0009653 (anatomical structure morphogenesis)
7 GO:0016192 (vesicle-mediated transport)
17 GO:0022414 (reproductive process)
17 GO:0000003 (reproduction)
34 GO:0050794 (regulation of cellular process)
0
5
10
15
20
25
30
35
-log(p_value)
Class
Molecular function
Cellular component
Biological process

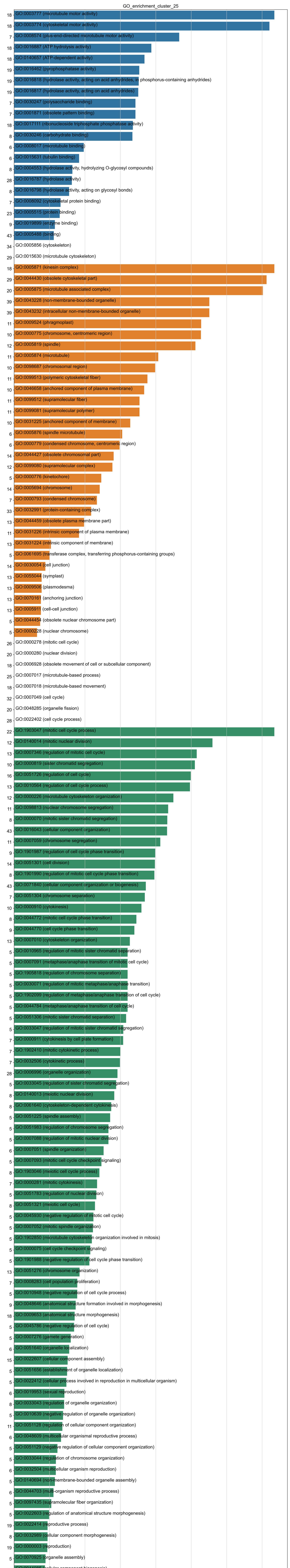
GO_enrichment_cluster_25
18 GO:0003777 (microtubule motor activity)
18 GO:0003774 (cytoskeletal motor activity)
7 GO:0008574 (plus-end-directed microtubule motor activity)
18 GO:0016887 (ATP hydrolysis activity)
18 GO:0140657 (ATP-dependent activity)
19 GO:0016462 (pyrophosphatase activity)
19 GO:0016818 (hydrolase activity, acting on acid anhydrides, in phosphorus-containing anhydrides)
19 GO:0016817 (hydrolase activity, acting on acid anhydrides)
7 GO:0030247 (polysaccharide binding)
7 GO:0001871 (obsolete pattern binding)
18 GO:0017111 (ribonucleoside triphosphate phosphatase activity)
8 GO:0030246 (carbohydrate binding)
6 GO:0008017 (microtubule binding)
6 GO:0015631 (tubulin binding)
8 GO:0004553 (hydrolase activity, hydrolyzing O-glycosyl compounds)
28 GO:0016787 (hydrolase activity)
8 GO:0016798 (hydrolase activity, acting on glycosyl bonds)
7 GO:0008092 (cytoskeletal protein binding)
23 GO:0005515 (protein binding)
9 GO:0019899 (enzyme binding)
43 GO:0005488 (binding)
34 GO:0005856 (cytoskeleton)
29 GO:0015630 (microtubule cytoskeleton)
18 GO:0005871 (kinesin complex)
29 GO:0044430 (obsolete cytoskeletal part)
20 GO:0005875 (microtubule associated complex)
39 GO:0043228 (non-membrane-bounded organelle)
39 GO:0043232 (intracellular non-membrane-bounded organelle)
11 GO:0009524 (phragmoplast)
10 GO:0000775 (chromosome, centromeric region)
12 GO:0005819 (spindle)
11 GO:0005874 (microtubule)
10 GO:0098687 (chromosomal region)
11 GO:0099513 (polymeric cytoskeletal fiber)
10 GO:0046658 (anchored component of plasma membrane)
11 GO:0099512 (supramolecular fiber)
11 GO:0099081 (supramolecular polymer)
10 GO:0031225 (anchored component of membrane)
6 GO:0005876 (spindle microtubule)
6 GO:0000779 (condensed chromosome, centromeric region)
14 GO:0044427 (obsolete chromosomal part)
12 GO:0099080 (supramolecular complex)
5 GO:0000776 (kinetochore)
14 GO:0005694 (chromosome)
7 GO:0000793 (condensed chromosome)
33 GO:0032991 (protein-containing complex)
13 GO:0044459 (obsolete plasma membrane part)
11 GO:0031226 (intrinsic component of plasma membrane)
13 GO:0031224 (intrinsic component of membrane)
5 GO:0061695 (transferase complex, transferring phosphorus-containing groups)
14 GO:0030054 (cell junction)
13 GO:0055044 (symplast)
13 GO:0009506 (plasmodesma)
13 GO:0070161 (anchoring junction)
13 GO:0005911 (cell-cell junction)
5 GO:0044454 (obsolete nuclear chromosome part)
5 GO:0000228 (nuclear chromosome)
26 GO:0000278 (mitotic cell cycle)
20 GO:0000280 (nuclear division)
18 GO:0006928 (obsolete movement of cell or subcellular component)
25 GO:0007017 (microtubule-based process)
18 GO:0007018 (microtubule-based movement)
32 GO:0007049 (cell cycle)
20 GO:0048285 (organelle fission)
28 GO:0022402 (cell cycle process)
22 GO:1903047 (mitotic cell cycle process)
12 GO:0140014 (mitotic nuclear division)
13 GO:0007346 (regulation of mitotic cell cycle)
10 GO:0000819 (sister chromatid segregation)
16 GO:0051726 (regulation of cell cycle)
13 GO:0010564 (regulation of cell cycle process)
12 GO:0000226 (microtubule cytoskeleton organization)
11 GO:0098813 (nuclear chromosome segregation)
8 GO:0000070 (mitotic sister chromatid segregation)
43 GO:0016043 (cellular component organization)
11 GO:0007059 (chromosome segregation)
9 GO:1901987 (regulation of cell cycle phase transition)
14 GO:0051301 (cell division)
8 GO:1901990 (regulation of mitotic cell cycle phase transition)
43 GO:0071840 (cellular component organization or biogenesis)
7 GO:0051304 (chromosome separation)
10 GO:0000910 (cytokinesis)
8 GO:0044772 (mitotic cell cycle phase transition)
9 GO:0044770 (cell cycle phase transition)
13 GO:0007010 (cytoskeleton organization)
5 GO:0010965 (regulation of mitotic sister chromatid separation)
5 GO:0007091 (metaphase/anaphase transition of mitotic cell cycle)
5 GO:1905818 (regulation of chromosome separation)
5 GO:0030071 (regulation of mitotic metaphase/anaphase transition)
5 GO:1902099 (regulation of metaphase/anaphase transition of cell cycle)
5 GO:0044784 (metaphase/anaphase transition of cell cycle)
5 GO:0051306 (mitotic sister chromatid separation)
5 GO:0033047 (regulation of mitotic sister chromatid segregation)
7 GO:0000911 (cytokinesis by cell plate formation)
7 GO:1902410 (mitotic cytokinetic process)
7 GO:0032506 (cytokinetic process)
28 GO:0006996 (organelle organization)
5 GO:0033045 (regulation of sister chromatid segregation)
8 GO:0140013 (meiotic nuclear division)
8 GO:0061640 (cytoskeleton-dependent cytokinesis)
5 GO:0051225 (spindle assembly)
5 GO:0051983 (regulation of chromosome segregation)
5 GO:0007088 (regulation of mitotic nuclear division)
6 GO:0007051 (spindle organization)
5 GO:0007093 (mitotic cell cycle checkpoint signaling)
8 GO:1903046 (meiotic cell cycle process)
7 GO:0000281 (mitotic cytokinesis)
5 GO:0051783 (regulation of nuclear division)
8 GO:0051321 (meiotic cell cycle)
5 GO:0045930 (negative regulation of mitotic cell cycle)
5 GO:0007052 (mitotic spindle organization)
5 GO:1902850 (microtubule cytoskeleton organization involved in mitosis)
5 GO:0000075 (cell cycle checkpoint signaling)
5 GO:1901988 (negative regulation of cell cycle phase transition)
13 GO:0051276 (chromosome organization)
7 GO:0008283 (cell population proliferation)
5 GO:0010948 (negative regulation of cell cycle process)
9 GO:0048646 (anatomical structure formation involved in morphogenesis)
18 GO:0009653 (anatomical structure morphogenesis)
5 GO:0045786 (negative regulation of cell cycle)
5 GO:0007276 (gamete generation)
6 GO:0051640 (organelle localization)
15 GO:0022607 (cellular component assembly)
5 GO:0051656 (establishment of organelle localization)
5 GO:0022412 (cellular process involved in reproduction in multicellular organism)
6 GO:0019953 (sexual reproduction)
8 GO:0033043 (regulation of organelle organization)
5 GO:0010639 (negative regulation of organelle organization)
11 GO:0051128 (regulation of cellular component organization)
6 GO:0048609 (multicellular organismal reproductive process)
5 GO:0051129 (negative regulation of cellular component organization)
5 GO:0033044 (regulation of chromosome organization)
6 GO:0032504 (multicellular organism reproduction)
5 GO:0140694 (non-membrane-bounded organelle assembly)
6 GO:0044703 (multi-organism reproductive process)
5 GO:0097435 (supramolecular fiber organization)
5 GO:0022603 (regulation of anatomical structure morphogenesis)
19 GO:0022414 (reproductive process)
8 GO:0032989 (cellular component morphogenesis)
19 GO:0000003 (reproduction)
5 GO:0070925 (organelle assembly)
15 GO:0044085 (cellular component biogenesis)
5 GO:0007389 (pattern specification process)
6 GO:0009555 (pollen development)
10 GO:0009888 (tissue development)
0 5 10 15 20 25 30 35
-log(p_value)
Class
Molecular function
Cellular component
Biological process

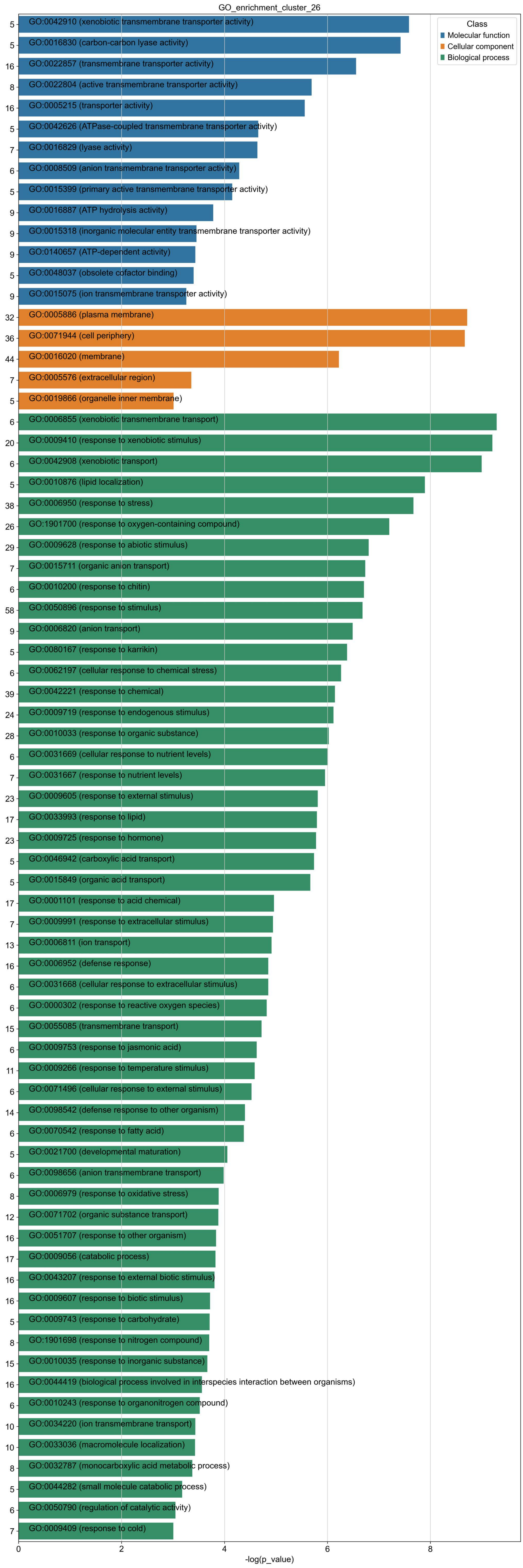
GO_enrichment_cluster_26
Class
Molecular function
Cellular component
Biological process
5 GO:0042910 (xenobiotic transmembrane transporter activity)
5 GO:0016830 (carbon-carbon lyase activity)
16 GO:0022857 (transmembrane transporter activity)
8 GO:0022804 (active transmembrane transporter activity)
16 GO:0005215 (transporter activity)
5 GO:0042626 (ATPase-coupled transmembrane transporter activity)
7 GO:0016829 (lyase activity)
6 GO:0008509 (anion transmembrane transporter activity)
5 GO:0015399 (primary active transmembrane transporter activity)
9 GO:0016887 (ATP hydrolysis activity)
9 GO:0015318 (inorganic molecular entity transmembrane transporter activity)
9 GO:0140657 (ATP-dependent activity)
5 GO:0048037 (obsolete cofactor binding)
9 GO:0015075 (ion transmembrane transporter activity)
32 GO:0005886 (plasma membrane)
36 GO:0071944 (cell periphery)
44 GO:0016020 (membrane)
7 GO:0005576 (extracellular region)
5 GO:0019866 (organelle inner membrane)
6 GO:0006855 (xenobiotic transmembrane transport)
20 GO:0009410 (response to xenobiotic stimulus)
6 GO:0042908 (xenobiotic transport)
5 GO:0010876 (lipid localization)
38 GO:0006950 (response to stress)
26 GO:1901700 (response to oxygen-containing compound)
29 GO:0009628 (response to abiotic stimulus)
7 GO:0015711 (organic anion transport)
6 GO:0010200 (response to chitin)
58 GO:0050896 (response to stimulus)
9 GO:0006820 (anion transport)
5 GO:0080167 (response to karrikin)
6 GO:0062197 (cellular response to chemical stress)
39 GO:0042221 (response to chemical)
24 GO:0009719 (response to endogenous stimulus)
28 GO:0010033 (response to organic substance)
6 GO:0031669 (cellular response to nutrient levels)
7 GO:0031667 (response to nutrient levels)
23 GO:0009605 (response to external stimulus)
17 GO:0033993 (response to lipid)
23 GO:0009725 (response to hormone)
5 GO:0046942 (carboxylic acid transport)
5 GO:0015849 (organic acid transport)
17 GO:0001101 (response to acid chemical)
7 GO:0009991 (response to extracellular stimulus)
13 GO:0006811 (ion transport)
16 GO:0006952 (defense response)
6 GO:0031668 (cellular response to extracellular stimulus)
6 GO:0000302 (response to reactive oxygen species)
15 GO:0055085 (transmembrane transport)
6 GO:0009753 (response to jasmonic acid)
11 GO:0009266 (response to temperature stimulus)
6 GO:0071496 (cellular response to external stimulus)
14 GO:0098542 (defense response to other organism)
6 GO:0070542 (response to fatty acid)
5 GO:0021700 (developmental maturation)
6 GO:0098656 (anion transmembrane transport)
8 GO:0006979 (response to oxidative stress)
12 GO:0071702 (organic substance transport)
16 GO:0051707 (response to other organism)
17 GO:0009056 (catabolic process)
16 GO:0043207 (response to external biotic stimulus)
16 GO:0009607 (response to biotic stimulus)
5 GO:0009743 (response to carbohydrate)
8 GO:1901698 (response to nitrogen compound)
15 GO:0010035 (response to inorganic substance)
16 GO:0044419 (biological process involved in interspecies interaction between organisms)
6 GO:0010243 (response to organonitrogen compound)
10 GO:0034220 (ion transmembrane transport)
10 GO:0033036 (macromolecule localization)
8 GO:0032787 (monocarboxylic acid metabolic process)
5 GO:0044282 (small molecule catabolic process)
6 GO:0050790 (regulation of catalytic activity)
7 GO:0009409 (response to cold)
0
2
4
6
8
-log(p_value)

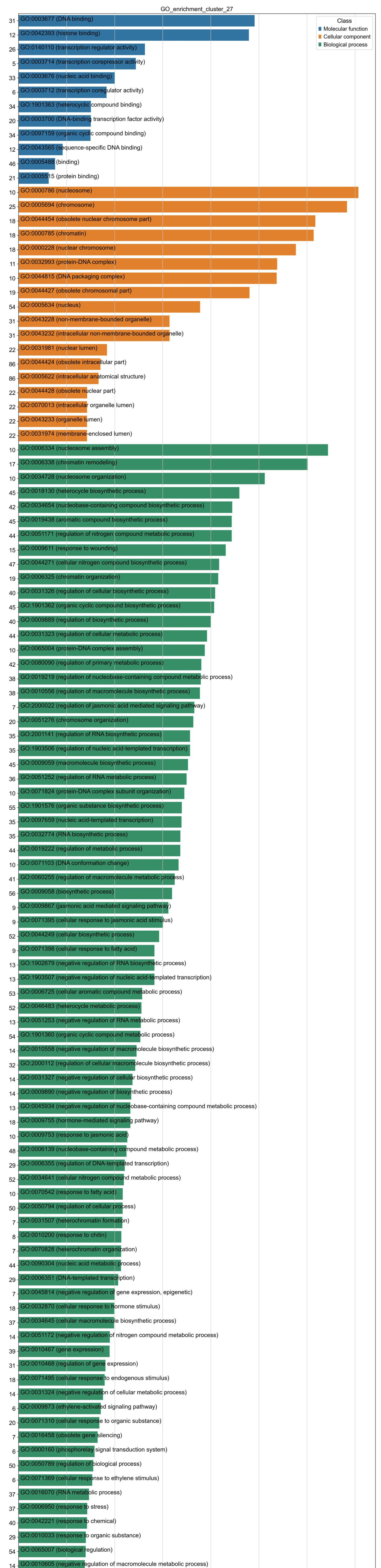
GO_enrichment_cluster_27
Class
Molecular function
Cellular component
Biological process
-log(p_value)

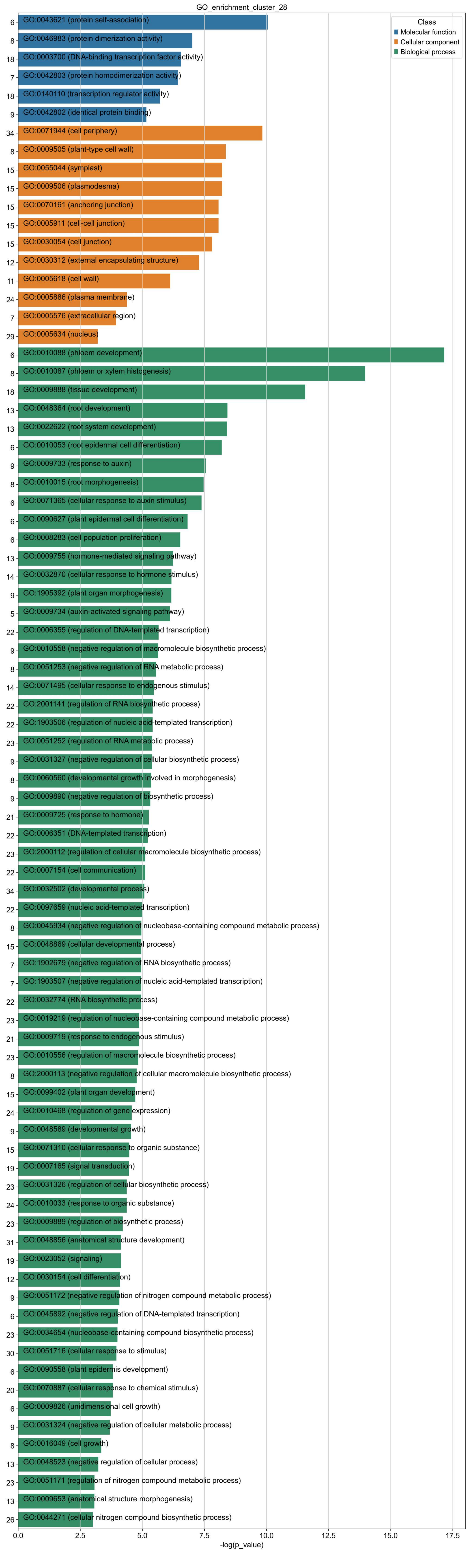
GO_enrichment_cluster_28
Class
Molecular function
Cellular component
Biological process
6 GO:0043621 (protein self-association)
8 GO:0046983 (protein dimerization activity)
18 GO:0003700 (DNA-binding transcription factor activity)
7 GO:0042803 (protein homodimerization activity)
18 GO:0140110 (transcription regulator activity)
9 GO:0042802 (identical protein binding)
34 GO:0071944 (cell periphery)
8 GO:0009505 (plant-type cell wall)
15 GO:0055044 (symplast)
15 GO:0009506 (plasmodesma)
15 GO:0070161 (anchoring junction)
15 GO:0005911 (cell-cell junction)
15 GO:0030054 (cell junction)
12 GO:0030312 (external encapsulating structure)
11 GO:0005618 (cell wall)
24 GO:0005886 (plasma membrane)
7 GO:0005576 (extracellular region)
29 GO:0005634 (nucleus)
6 GO:0010088 (phloem development)
8 GO:0010087 (phloem or xylem histogenesis)
18 GO:0009888 (tissue development)
13 GO:0048364 (root development)
13 GO:0022622 (root system development)
6 GO:0010053 (root epidermal cell differentiation)
9 GO:0009733 (response to auxin)
8 GO:0010015 (root morphogenesis)
6 GO:0071365 (cellular response to auxin stimulus)
6 GO:0090627 (plant epidermal cell differentiation)
6 GO:0008283 (cell population proliferation)
13 GO:0009755 (hormone-mediated signaling pathway)
14 GO:0032870 (cellular response to hormone stimulus)
9 GO:1905392 (plant organ morphogenesis)
5 GO:0009734 (auxin-activated signaling pathway)
22 GO:0006355 (regulation of DNA-templated transcription)
9 GO:0010558 (negative regulation of macromolecule biosynthetic process)
8 GO:0051253 (negative regulation of RNA metabolic process)
14 GO:0071495 (cellular response to endogenous stimulus)
22 GO:2001141 (regulation of RNA biosynthetic process)
22 GO:1903506 (regulation of nucleic acid-templated transcription)
23 GO:0051252 (regulation of RNA metabolic process)
9 GO:0031327 (negative regulation of cellular biosynthetic process)
8 GO:0060560 (developmental growth involved in morphogenesis)
9 GO:0009890 (negative regulation of biosynthetic process)
21 GO:0009725 (response to hormone)
22 GO:0006351 (DNA-templated transcription)
23 GO:2000112 (regulation of cellular macromolecule biosynthetic process)
22 GO:0007154 (cell communication)
34 GO:0032502 (developmental process)
22 GO:0097659 (nucleic acid-templated transcription)
8 GO:0045934 (negative regulation of nucleobase-containing compound metabolic process)
15 GO:0048869 (cellular developmental process)
7 GO:1902679 (negative regulation of RNA biosynthetic process)
7 GO:1903507 (negative regulation of nucleic acid-templated transcription)
22 GO:0032774 (RNA biosynthetic process)
23 GO:0019219 (regulation of nucleobase-containing compound metabolic process)
21 GO:0009719 (response to endogenous stimulus)
23 GO:0010556 (regulation of macromolecule biosynthetic process)
8 GO:2000113 (negative regulation of cellular macromolecule biosynthetic process)
15 GO:0099402 (plant organ development)
24 GO:0010468 (regulation of gene expression)
9 GO:0048589 (developmental growth)
15 GO:0071310 (cellular response to organic substance)
19 GO:0007165 (signal transduction)
23 GO:0031326 (regulation of cellular biosynthetic process)
24 GO:0010033 (response to organic substance)
23 GO:0009889 (regulation of biosynthetic process)
31 GO:0048856 (anatomical structure development)
19 GO:0023052 (signaling)
12 GO:0030154 (cell differentiation)
9 GO:0051172 (negative regulation of nitrogen compound metabolic process)
6 GO:0045892 (negative regulation of DNA-templated transcription)
23 GO:0034654 (nucleobase-containing compound biosynthetic process)
30 GO:0051716 (cellular response to stimulus)
6 GO:0090558 (plant epidermis development)
20 GO:0070887 (cellular response to chemical stimulus)
6 GO:0009826 (unidimensional cell growth)
9 GO:0031324 (negative regulation of cellular metabolic process)
8 GO:0016049 (cell growth)
13 GO:0048523 (negative regulation of cellular process)
23 GO:0051171 (regulation of nitrogen compound metabolic process)
13 GO:0009653 (anatomical structure morphogenesis)
26 GO:0044271 (cellular nitrogen compound biosynthetic process)
0.0
2.5
5.0
7.5
10.0
12.5
15.0
17.5
-log(p_value)

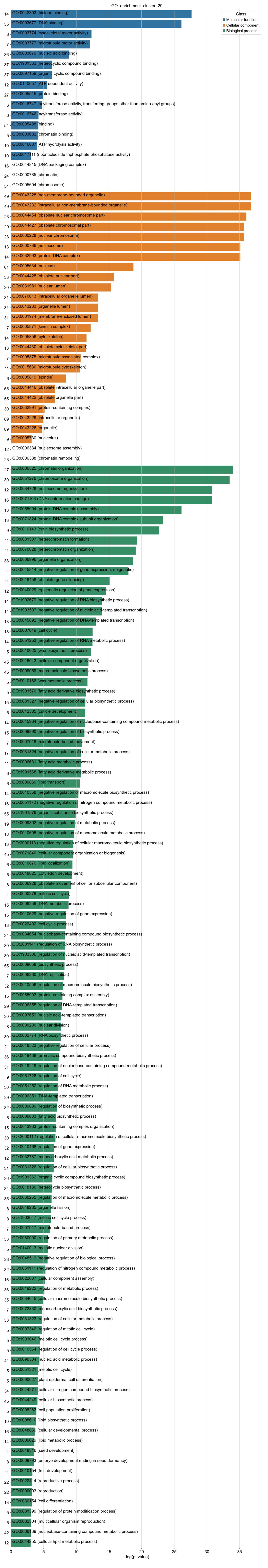

GO_enrichment_cluster_29
Class
Molecular function
Cellular component
Biological process
14 GO:0042393 (histone binding)
35 GO:0003677 (DNA binding)
8 GO:0003774 (cytoskeletal motor activity)
7 GO:0003777 (microtubule motor activity)
36 GO:0003676 (nucleic acid binding)
37 GO:1901363 (heterocyclic compound binding)
37 GO:0097159 (organic cyclic compound binding)
12 GO:0140657 (ATP-dependent activity)
27 GO:0005515 (protein binding)
6 GO:0016747 (acyltransferase activity, transferring groups other than amino-acyl groups)
6 GO:0016746 (acyltransferase activity)
54 GO:0005488 (binding)
5 GO:0003682 (chromatin binding)
10 GO:0016887 (ATP hydrolysis activity)
10 GO:0017111 (ribonucleoside triphosphate phosphatase activity)
16 GO:0044815 (DNA packaging complex)
24 GO:0000785 (chromatin)
34 GO:0005694 (chromosome)
49 GO:0043228 (non-membrane-bounded organelle)
49 GO:0043232 (intracellular non-membrane-bounded organelle)
23 GO:0044454 (obsolete nuclear chromosome part)
29 GO:0044427 (obsolete chromosomal part)
23 GO:0000228 (nuclear chromosome)
13 GO:0000786 (nucleosome)
14 GO:0032993 (protein-DNA complex)
61 GO:0005634 (nucleus)
33 GO:0044428 (obsolete nuclear part)
30 GO:0031981 (nuclear lumen)
31 GO:0070013 (intracellular organelle lumen)
31 GO:0043233 (organelle lumen)
31 GO:0031974 (membrane-enclosed lumen)
7 GO:0005871 (kinesin complex)
14 GO:0005856 (cytoskeleton)
13 GO:0044430 (obsolete cytoskeletal part)
7 GO:0005875 (microtubule associated complex)
11 GO:0015630 (microtubule cytoskeleton)
6 GO:0005819 (spindle)
55 GO:0044446 (obsolete intracellular organelle part)
55 GO:0044422 (obsolete organelle part)
30 GO:0032991 (protein-containing complex)
89 GO:0043229 (intracellular organelle)
89 GO:0043226 (organelle)
9 GO:0005730 (nucleolus)
12 GO:0006334 (nucleosome assembly)
23 GO:0006338 (chromatin remodeling)
27 GO:0006325 (chromatin organization)
30 GO:0051276 (chromosome organization)
12 GO:0034728 (nucleosome organization)
16 GO:0071103 (DNA conformation change)
13 GO:0065004 (protein-DNA complex assembly)
13 GO:0071824 (protein-DNA complex subunit organization)
9 GO:0010143 (cutin biosynthetic process)
11 GO:0031507 (heterochromatin formation)
11 GO:0070828 (heterochromatin organization)
38 GO:0006996 (organelle organization)
11 GO:0045814 (negative regulation of gene expression, epigenetic)
11 GO:0016458 (obsolete gene silencing)
12 GO:0040029 (epigenetic regulation of gene expression)
14 GO:1902679 (negative regulation of RNA biosynthetic process)
14 GO:1903507 (negative regulation of nucleic acid-templated transcription)
13 GO:0045892 (negative regulation of DNA-templated transcription)
18 GO:0007049 (cell cycle)
14 GO:0051253 (negative regulation of RNA metabolic process)
5 GO:0010025 (wax biosynthetic process)
45 GO:0016043 (cellular component organization)
45 GO:0009059 (macromolecule biosynthetic process)
5 GO:0010166 (wax metabolic process)
5 GO:1901570 (fatty acid derivative biosynthetic process)
15 GO:0031327 (negative regulation of cellular biosynthetic process)
5 GO:0042335 (cuticle development)
14 GO:0045934 (negative regulation of nucleobase-containing compound metabolic process)
15 GO:0009890 (negative regulation of biosynthetic process)
7 GO:0007018 (microtubule-based movement)
17 GO:0031324 (negative regulation of cellular metabolic process)
11 GO:0006631 (fatty acid metabolic process)
6 GO:1901568 (fatty acid derivative metabolic process)
6 GO:0006869 (lipid transport)
14 GO:0010558 (negative regulation of macromolecule biosynthetic process)
16 GO:0051172 (negative regulation of nitrogen compound metabolic process)
55 GO:1901576 (organic substance biosynthetic process)
19 GO:0009892 (negative regulation of metabolic process)
18 GO:0010605 (negative regulation of macromolecule metabolic process)
13 GO:2000113 (negative regulation of cellular macromolecule biosynthetic process)
45 GO:0071840 (cellular component organization or biogenesis)
6 GO:0010876 (lipid localization)
5 GO:0048825 (cotyledon development)
8 GO:0006928 (obsolete movement of cell or subcellular component)
11 GO:0000278 (mitotic cell cycle)
15 GO:0006259 (DNA metabolic process)
15 GO:0010629 (negative regulation of gene expression)
13 GO:0022402 (cell cycle process)
34 GO:0034654 (nucleobase-containing compound biosynthetic process)
30 GO:2001141 (regulation of RNA biosynthetic process)
30 GO:1903506 (regulation of nucleic acid-templated transcription)
55 GO:0009058 (biosynthetic process)
7 GO:0006260 (DNA replication)
32 GO:0010556 (regulation of macromolecule biosynthetic process)
15 GO:0065003 (protein-containing complex assembly)
29 GO:0006355 (regulation of DNA-templated transcription)
30 GO:0097659 (nucleic acid-templated transcription)
8 GO:0000280 (nuclear division)
30 GO:0032774 (RNA biosynthetic process)
21 GO:0048523 (negative regulation of cellular process)
36 GO:0019438 (aromatic compound biosynthetic process)
31 GO:0019219 (regulation of nucleobase-containing compound metabolic process)
9 GO:0051726 (regulation of cell cycle)
30 GO:0051252 (regulation of RNA metabolic process)
29 GO:0006351 (DNA-templated transcription)
32 GO:0009889 (regulation of biosynthetic process)
6 GO:0006633 (fatty acid biosynthetic process)
15 GO:0043933 (protein-containing complex organization)
30 GO:2000112 (regulation of cellular macromolecule biosynthetic process)
32 GO:0010468 (regulation of gene expression)
12 GO:0032787 (monocarboxylic acid metabolic process)
31 GO:0031326 (regulation of cellular biosynthetic process)
36 GO:1901362 (organic cyclic compound biosynthetic process)
34 GO:0018130 (heterocycle biosynthetic process)
35 GO:0060255 (regulation of macromolecule metabolic process)
8 GO:0048285 (organelle fission)
8 GO:1903047 (mitotic cell cycle process)
7 GO:0007017 (microtubule-based process)
33 GO:0080090 (regulation of primary metabolic process)
5 GO:0140013 (meiotic nuclear division)
23 GO:0048519 (negative regulation of biological process)
32 GO:0051171 (regulation of nitrogen compound metabolic process)
16 GO:0022607 (cellular component assembly)
36 GO:0019222 (regulation of metabolic process)
35 GO:0034645 (cellular macromolecule biosynthetic process)
7 GO:0072330 (monocarboxylic acid biosynthetic process)
33 GO:0031323 (regulation of cellular metabolic process)
5 GO:0007346 (regulation of mitotic cell cycle)
5 GO:1903046 (meiotic cell cycle process)
5 GO:0010564 (regulation of cell cycle process)
41 GO:0090304 (nucleic acid metabolic process)
5 GO:0051321 (meiotic cell cycle)
5 GO:0090627 (plant epidermal cell differentiation)
34 GO:0044271 (cellular nitrogen compound biosynthetic process)
45 GO:0044249 (cellular biosynthetic process)
5 GO:0008283 (cell population proliferation)
10 GO:0008610 (lipid biosynthetic process)
16 GO:0048869 (cellular developmental process)
14 GO:0006629 (lipid metabolic process)
11 GO:0048316 (seed development)
8 GO:0009793 (embryo development ending in seed dormancy)
11 GO:0010154 (fruit development)
22 GO:0022414 (reproductive process)
22 GO:0000003 (reproduction)
13 GO:0030154 (cell differentiation)
5 GO:0031399 (regulation of protein modification process)
5 GO:0032504 (multicellular organism reproduction)
42 GO:0006139 (nucleobase-containing compound metabolic process)
12 GO:0044255 (cellular lipid metabolic process)
0
5
10
15
20
25
30
35
-log(p_value)

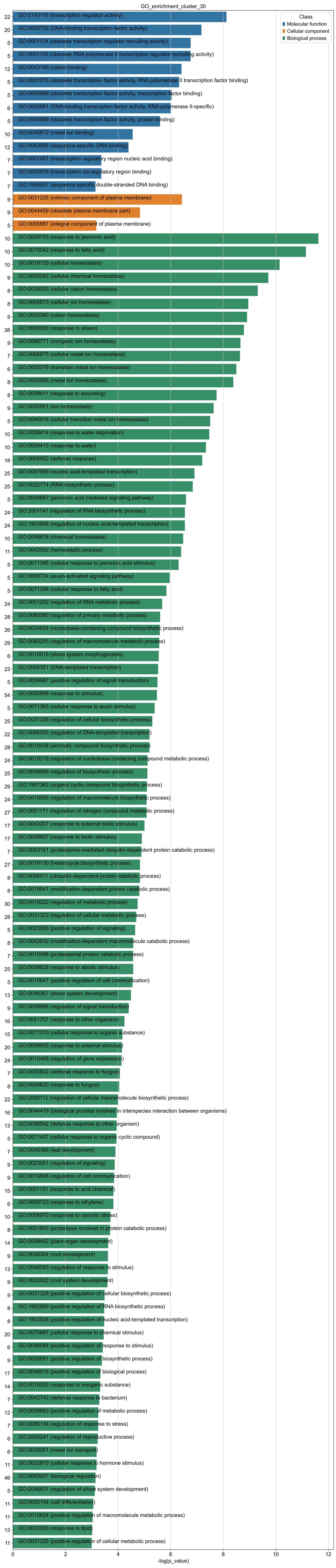
GO_enrichment_cluster_30
Class
Molecular function
Cellular component
Biological process
22 GO:0140110 (transcription regulator activity)
20 GO:0003700 (DNA-binding transcription factor activity)
5 GO:0001134 (obsolete transcription regulator recruiting activity)
5 GO:0001135 (obsolete RNA polymerase II transcription regulator recruiting activity)
12 GO:0043169 (cation binding)
5 GO:0001076 (obsolete transcription factor activity, RNA polymerase II transcription factor binding)
5 GO:0000989 (obsolete transcription factor activity, transcription factor binding)
6 GO:0000981 (DNA-binding transcription factor activity, RNA polymerase II-specific)
5 GO:0000988 (obsolete transcription factor activity, protein binding)
10 GO:0046872 (metal ion binding)
12 GO:0043565 (sequence-specific DNA binding)
7 GO:0001067 (transcription regulatory region nucleic acid binding)
7 GO:0000976 (transcription cis-regulatory region binding)
7 GO:1990837 (sequence-specific double-stranded DNA binding)
9 GO:0031226 (intrinsic component of plasma membrane)
9 GO:0044459 (obsolete plasma membrane part)
5 GO:0005887 (integral component of plasma membrane)
10 GO:0009753 (response to jasmonic acid)
10 GO:0070542 (response to fatty acid)
10 GO:0019725 (cellular homeostasis)
9 GO:0055082 (cellular chemical homeostasis)
8 GO:0030003 (cellular cation homeostasis)
8 GO:0006873 (cellular ion homeostasis)
9 GO:0055080 (cation homeostasis)
38 GO:0006950 (response to stress)
9 GO:0098771 (inorganic ion homeostasis)
7 GO:0006875 (cellular metal ion homeostasis)
6 GO:0055076 (transition metal ion homeostasis)
8 GO:0055065 (metal ion homeostasis)
8 GO:0009611 (response to wounding)
9 GO:0050801 (ion homeostasis)
5 GO:0046916 (cellular transition metal ion homeostasis)
10 GO:0009414 (response to water deprivation)
10 GO:0009415 (response to water)
18 GO:0006952 (defense response)
25 GO:0097659 (nucleic acid-templated transcription)
25 GO:0032774 (RNA biosynthetic process)
5 GO:0009867 (jasmonic acid mediated signaling pathway)
24 GO:2001141 (regulation of RNA biosynthetic process)
24 GO:1903506 (regulation of nucleic acid-templated transcription)
10 GO:0048878 (chemical homeostasis)
11 GO:0042592 (homeostatic process)
5 GO:0071395 (cellular response to jasmonic acid stimulus)
5 GO:0009734 (auxin-activated signaling pathway)
5 GO:0071398 (cellular response to fatty acid)
24 GO:0051252 (regulation of RNA metabolic process)
28 GO:0080090 (regulation of primary metabolic process)
26 GO:0034654 (nucleobase-containing compound biosynthetic process)
29 GO:0060255 (regulation of macromolecule metabolic process)
6 GO:0010016 (shoot system morphogenesis)
23 GO:0006351 (DNA-templated transcription)
5 GO:0009967 (positive regulation of signal transduction)
54 GO:0050896 (response to stimulus)
5 GO:0071365 (cellular response to auxin stimulus)
25 GO:0031326 (regulation of cellular biosynthetic process)
22 GO:0006355 (regulation of DNA-templated transcription)
28 GO:0019438 (aromatic compound biosynthetic process)
24 GO:0019219 (regulation of nucleobase-containing compound metabolic process)
25 GO:0009889 (regulation of biosynthetic process)
29 GO:1901362 (organic cyclic compound biosynthetic process)
24 GO:0010556 (regulation of macromolecule biosynthetic process)
27 GO:0051171 (regulation of nitrogen compound metabolic process)
17 GO:0043207 (response to external biotic stimulus)
17 GO:0009607 (response to biotic stimulus)
7 GO:0043161 (proteasome-mediated ubiquitin-dependent protein catabolic process)
27 GO:0018130 (heterocycle biosynthetic process)
8 GO:0006511 (ubiquitin-dependent protein catabolic process)
8 GO:0019941 (modification-dependent protein catabolic process)
30 GO:0019222 (regulation of metabolic process)
28 GO:0031323 (regulation of cellular metabolic process)
5 GO:0023056 (positive regulation of signaling)
8 GO:0043632 (modification-dependent macromolecule catabolic process)
7 GO:0010498 (proteasomal protein catabolic process)
25 GO:0009628 (response to abiotic stimulus)
5 GO:0010647 (positive regulation of cell communication)
13 GO:0048367 (shoot system development)
9 GO:0009966 (regulation of signal transduction)
16 GO:0051707 (response to other organism)
15 GO:0071310 (cellular response to organic substance)
20 GO:0009605 (response to external stimulus)
24 GO:0010468 (regulation of gene expression)
7 GO:0050832 (defense response to fungus)
8 GO:0009620 (response to fungus)
22 GO:2000112 (regulation of cellular macromolecule biosynthetic process)
16 GO:0044419 (biological process involved in interspecies interaction between organisms)
13 GO:0098542 (defense response to other organism)
5 GO:0071407 (cellular response to organic cyclic compound)
7 GO:0048366 (leaf development)
9 GO:0023051 (regulation of signaling)
9 GO:0010646 (regulation of cell communication)
15 GO:0001101 (response to acid chemical)
6 GO:0009723 (response to ethylene)
10 GO:0006970 (response to osmotic stress)
8 GO:0051603 (proteolysis involved in protein catabolic process)
14 GO:0099402 (plant organ development)
9 GO:0048364 (root development)
13 GO:0048583 (regulation of response to stimulus)
9 GO:0022622 (root system development)
9 GO:0031328 (positive regulation of cellular biosynthetic process)
8 GO:1902680 (positive regulation of RNA biosynthetic process)
8 GO:1903508 (positive regulation of nucleic acid-templated transcription)
20 GO:0070887 (cellular response to chemical stimulus)
6 GO:0048584 (positive regulation of response to stimulus)
9 GO:0009891 (positive regulation of biosynthetic process)
17 GO:0048518 (positive regulation of biological process)
14 GO:0010035 (response to inorganic substance)
7 GO:0042742 (defense response to bacterium)
12 GO:0009893 (positive regulation of metabolic process)
7 GO:0080134 (regulation of response to stress)
6 GO:2000241 (regulation of reproductive process)
6 GO:0030001 (metal ion transport)
11 GO:0032870 (cellular response to hormone stimulus)
46 GO:0065007 (biological regulation)
5 GO:0048831 (regulation of shoot system development)
11 GO:0030154 (cell differentiation)
11 GO:0010604 (positive regulation of macromolecule metabolic process)
13 GO:0033993 (response to lipid)
11 GO:0031325 (positive regulation of cellular metabolic process)
0 2 4 6 8 10 12
-log(p_value)

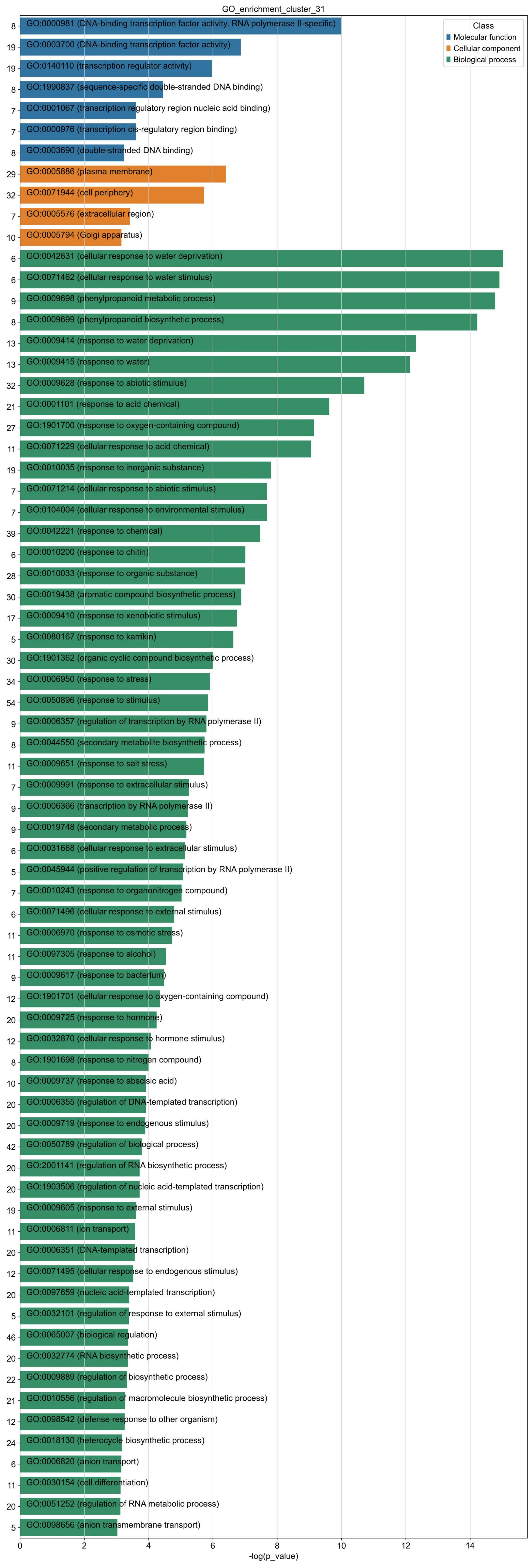
GO_enrichment_cluster_31
Class
Molecular function
Cellular component
Biological process
8 GO:0000981 (DNA-binding transcription factor activity, RNA polymerase II-specific)
19 GO:0003700 (DNA-binding transcription factor activity)
19 GO:0140110 (transcription regulator activity)
8 GO:1990837 (sequence-specific double-stranded DNA binding)
7 GO:0001067 (transcription regulatory region nucleic acid binding)
7 GO:0000976 (transcription cis-regulatory region binding)
8 GO:0003690 (double-stranded DNA binding)
29 GO:0005886 (plasma membrane)
32 GO:0071944 (cell periphery)
7 GO:0005576 (extracellular region)
10 GO:0005794 (Golgi apparatus)
6 GO:0042631 (cellular response to water deprivation)
6 GO:0071462 (cellular response to water stimulus)
9 GO:0009698 (phenylpropanoid metabolic process)
8 GO:0009699 (phenylpropanoid biosynthetic process)
13 GO:0009414 (response to water deprivation)
13 GO:0009415 (response to water)
32 GO:0009628 (response to abiotic stimulus)
21 GO:0001101 (response to acid chemical)
27 GO:1901700 (response to oxygen-containing compound)
11 GO:0071229 (cellular response to acid chemical)
19 GO:0010035 (response to inorganic substance)
7 GO:0071214 (cellular response to abiotic stimulus)
7 GO:0104004 (cellular response to environmental stimulus)
39 GO:0042221 (response to chemical)
6 GO:0010200 (response to chitin)
28 GO:0010033 (response to organic substance)
30 GO:0019438 (aromatic compound biosynthetic process)
17 GO:0009410 (response to xenobiotic stimulus)
5 GO:0080167 (response to karrikin)
30 GO:1901362 (organic cyclic compound biosynthetic process)
34 GO:0006950 (response to stress)
54 GO:0050896 (response to stimulus)
9 GO:0006357 (regulation of transcription by RNA polymerase II)
8 GO:0044550 (secondary metabolite biosynthetic process)
11 GO:0009651 (response to salt stress)
7 GO:0009991 (response to extracellular stimulus)
9 GO:0006366 (transcription by RNA polymerase II)
9 GO:0019748 (secondary metabolic process)
6 GO:0031668 (cellular response to extracellular stimulus)
5 GO:0045944 (positive regulation of transcription by RNA polymerase II)
7 GO:0010243 (response to organonitrogen compound)
6 GO:0071496 (cellular response to external stimulus)
11 GO:0006970 (response to osmotic stress)
11 GO:0097305 (response to alcohol)
9 GO:0009617 (response to bacterium)
12 GO:1901701 (cellular response to oxygen-containing compound)
20 GO:0009725 (response to hormone)
12 GO:0032870 (cellular response to hormone stimulus)
8 GO:1901698 (response to nitrogen compound)
10 GO:0009737 (response to abscisic acid)
20 GO:0006355 (regulation of DNA-templated transcription)
20 GO:0009719 (response to endogenous stimulus)
42 GO:0050789 (regulation of biological process)
20 GO:2001141 (regulation of RNA biosynthetic process)
20 GO:1903506 (regulation of nucleic acid-templated transcription)
19 GO:0009605 (response to external stimulus)
11 GO:0006811 (ion transport)
20 GO:0006351 (DNA-templated transcription)
12 GO:0071495 (cellular response to endogenous stimulus)
20 GO:0097659 (nucleic acid-templated transcription)
5 GO:0032101 (regulation of response to external stimulus)
46 GO:0065007 (biological regulation)
20 GO:0032774 (RNA biosynthetic process)
22 GO:0009889 (regulation of biosynthetic process)
21 GO:0010556 (regulation of macromolecule biosynthetic process)
12 GO:0098542 (defense response to other organism)
24 GO:0018130 (heterocycle biosynthetic process)
6 GO:0006820 (anion transport)
11 GO:0030154 (cell differentiation)
20 GO:0051252 (regulation of RNA metabolic process)
5 GO:0098656 (anion transmembrane transport)
0
2
4
6
8
10
12
14
-log(p_value)

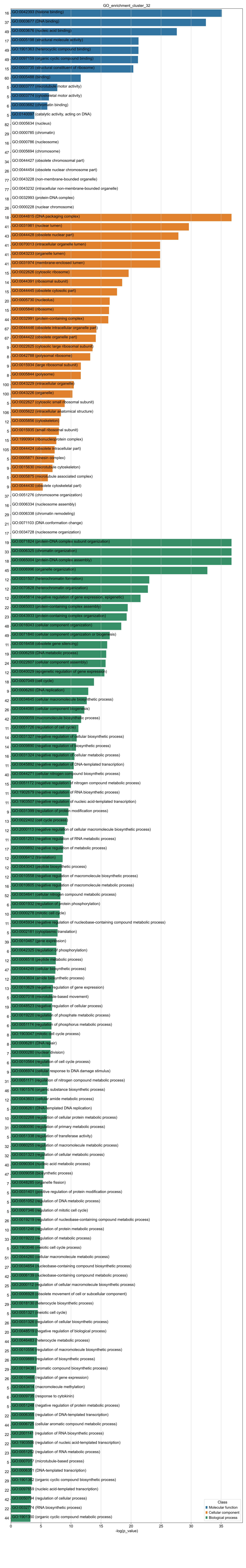
GO_enrichment_cluster_32
-log(p_value)
Class
Molecular function
Cellular component
Biological process
