## Supplementary Figures S1~S9 for "An integrative single-cell and spatial transcriptomics atlas highlights candidate regulatory factors in the development of gerbera capitulum"

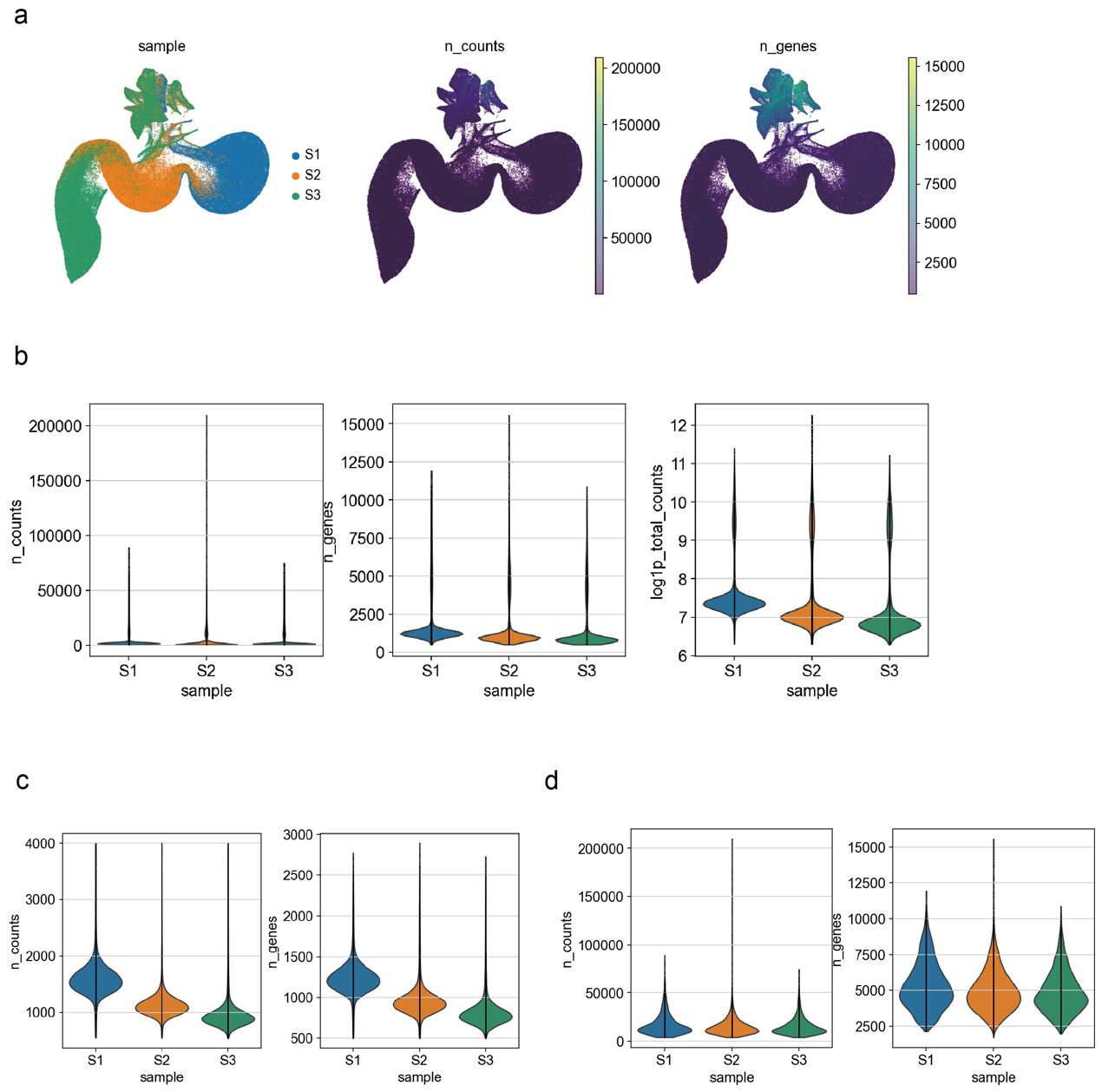

**Supplementary Figure S1.** The overview of unfiltered single-cell transcriptomes. (a) The distribution of the sample, gene expression counts, and expressed gene number of cells on the UMAP embedding. (b) The violin graph showing the cell distribution with certain gene expression count, expressed gene number, and logarithmized gene expression count (Y-axis) for each sample (X-axis). (c) The distribution of cells with gene expression counts of at most 4,000. (d) The distribution of cells with gene expression counts over 4,000.

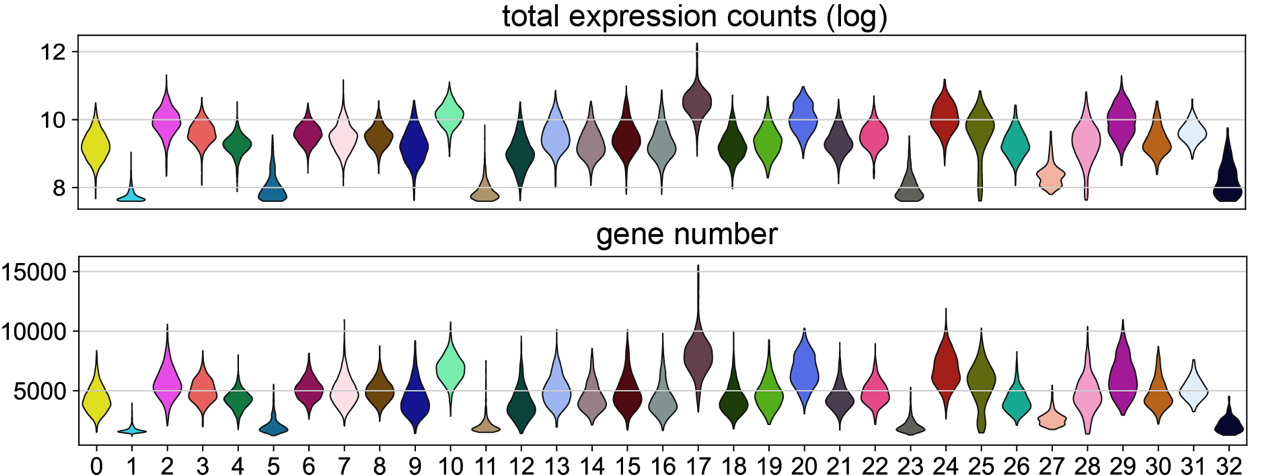

**Supplementary Figure S2.** The distribution of the gene expression counts and expressed gene number of all clusters for cells after filtering on the UMAP embedding.

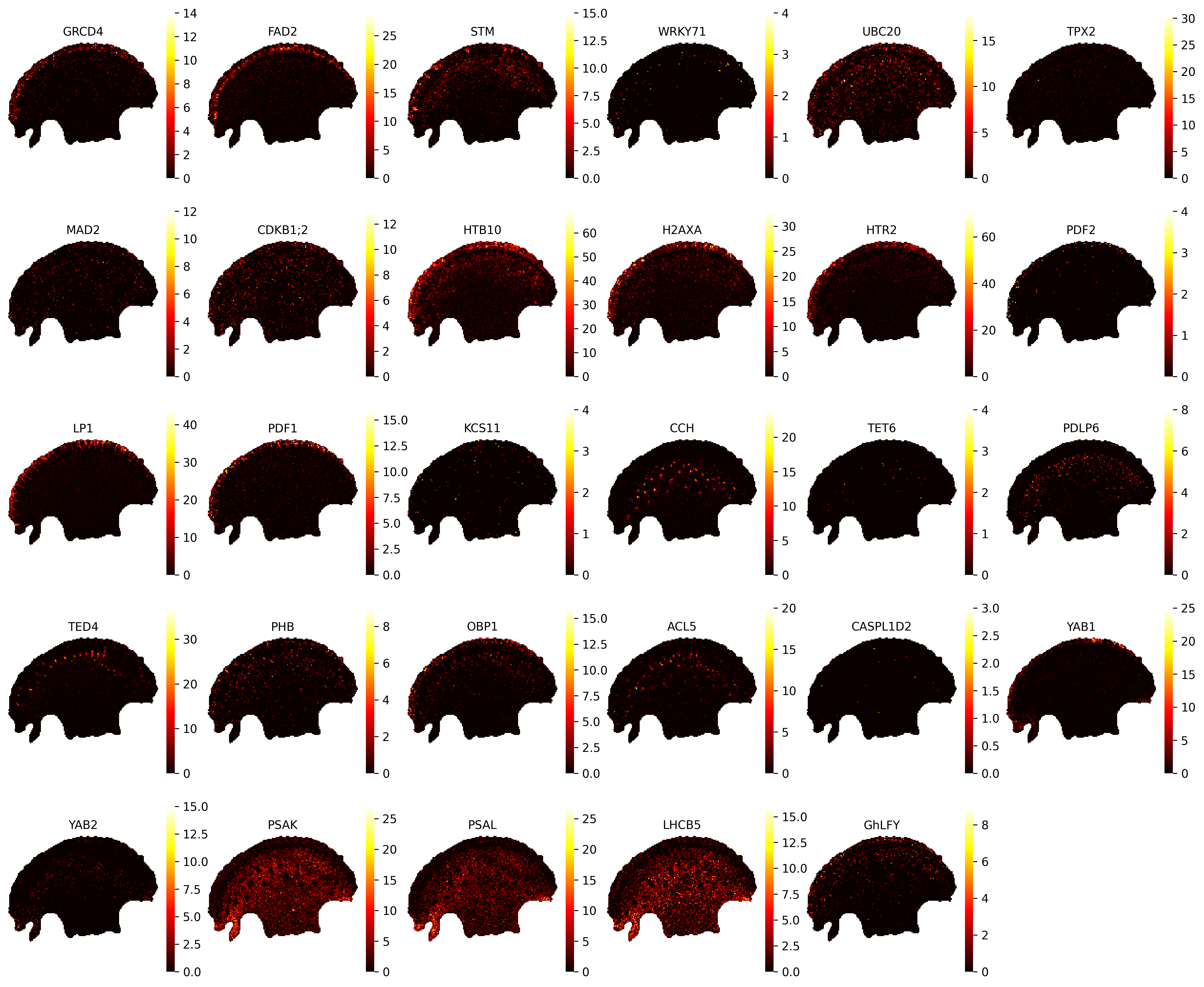

**Supplementary Figure S3.** The spatial expression of single-cell transcriptomics marker genes in the stereo-seq sample SP1_d.

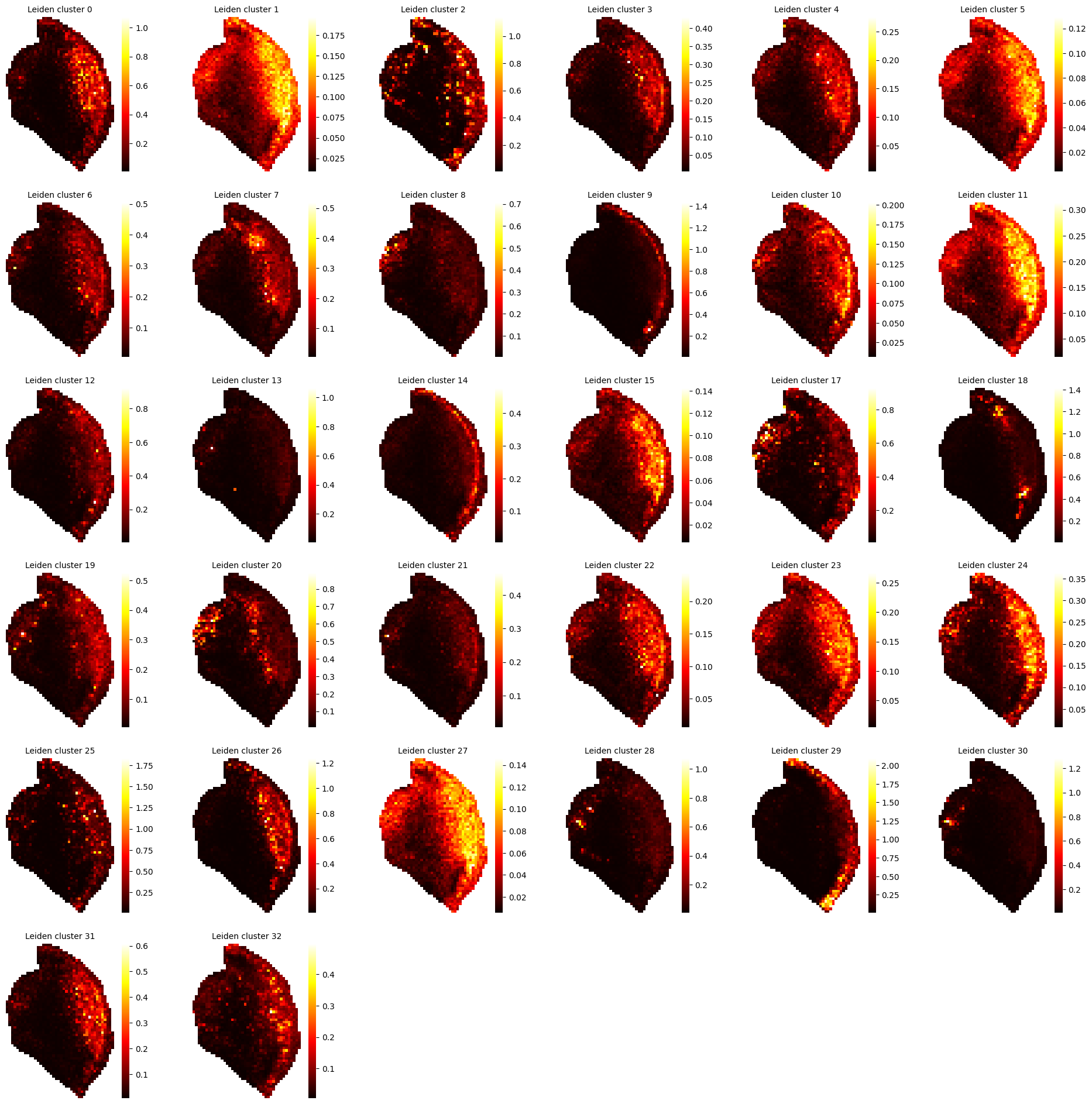

**Supplementary Figure S4.** The cell2location result of mapping cell cluster in single-cell transcriptome S1 onto spatial transcriptome SP3_g. The color shows the inferred cell count in each cluster within each pseudocell.

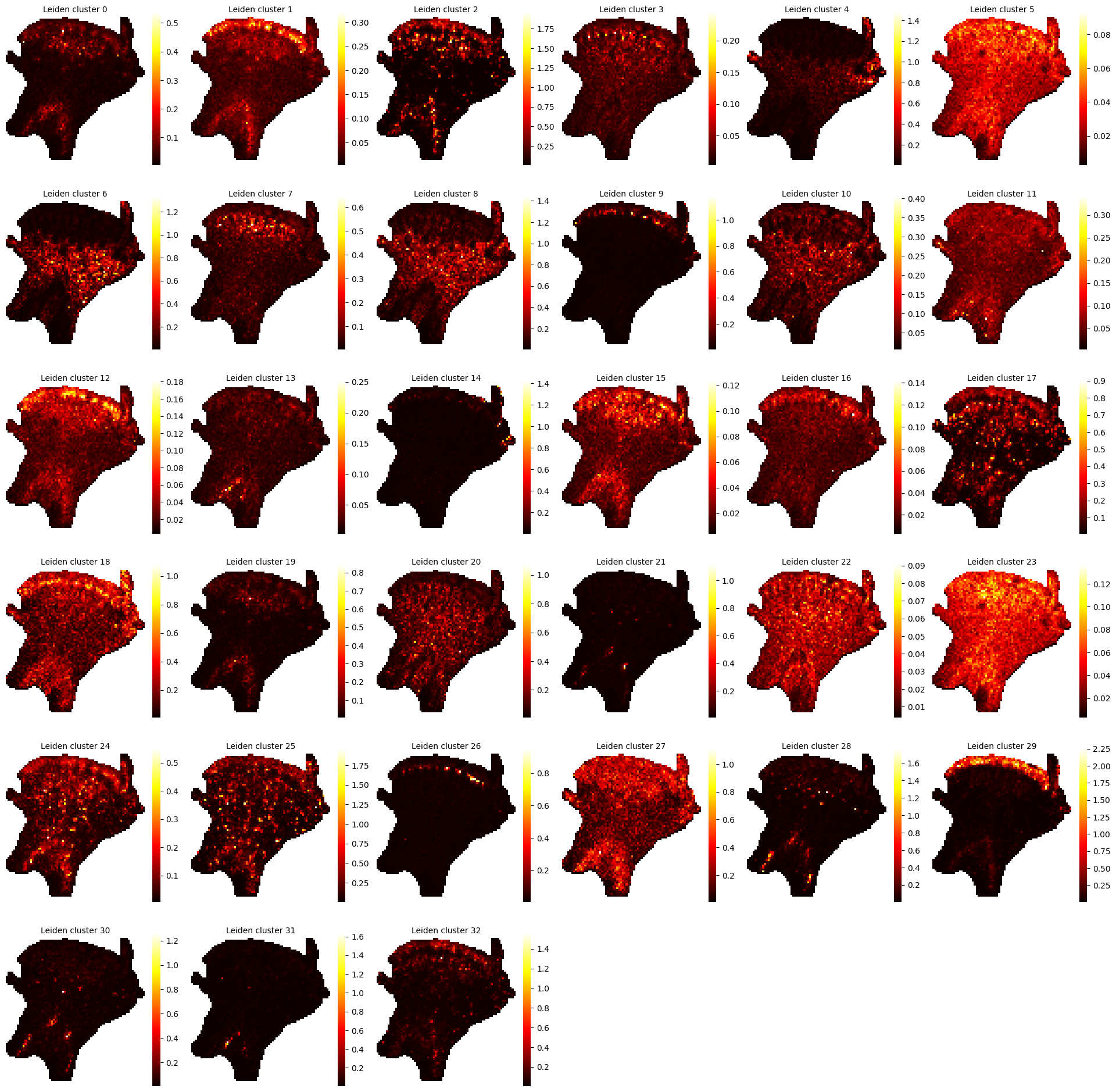

**Supplementary Figure S5.** The cell2location result of mapping cell clusters in single-cell transcriptome S2 onto spatial transcriptome SP3_f. The color shows the inferred cell count in each cluster within each pseudocell.

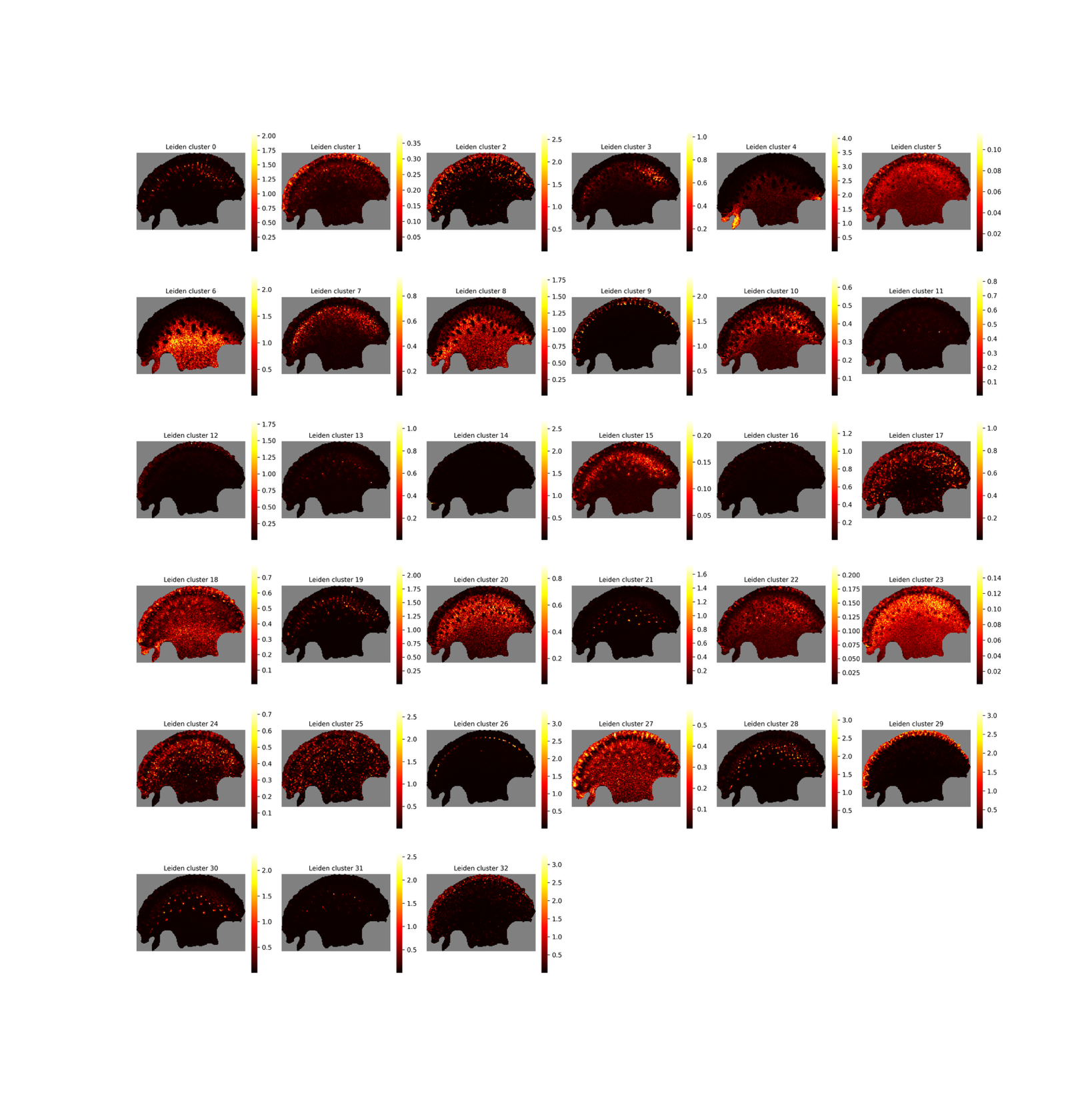

**Supplementary Figure S6.** The cell2location result of mapping cell cluster in single-cell transcriptome S3 onto the spatial transcriptome SP1_d. The color shows the inferred cell count in each cluster within each pseudocell.

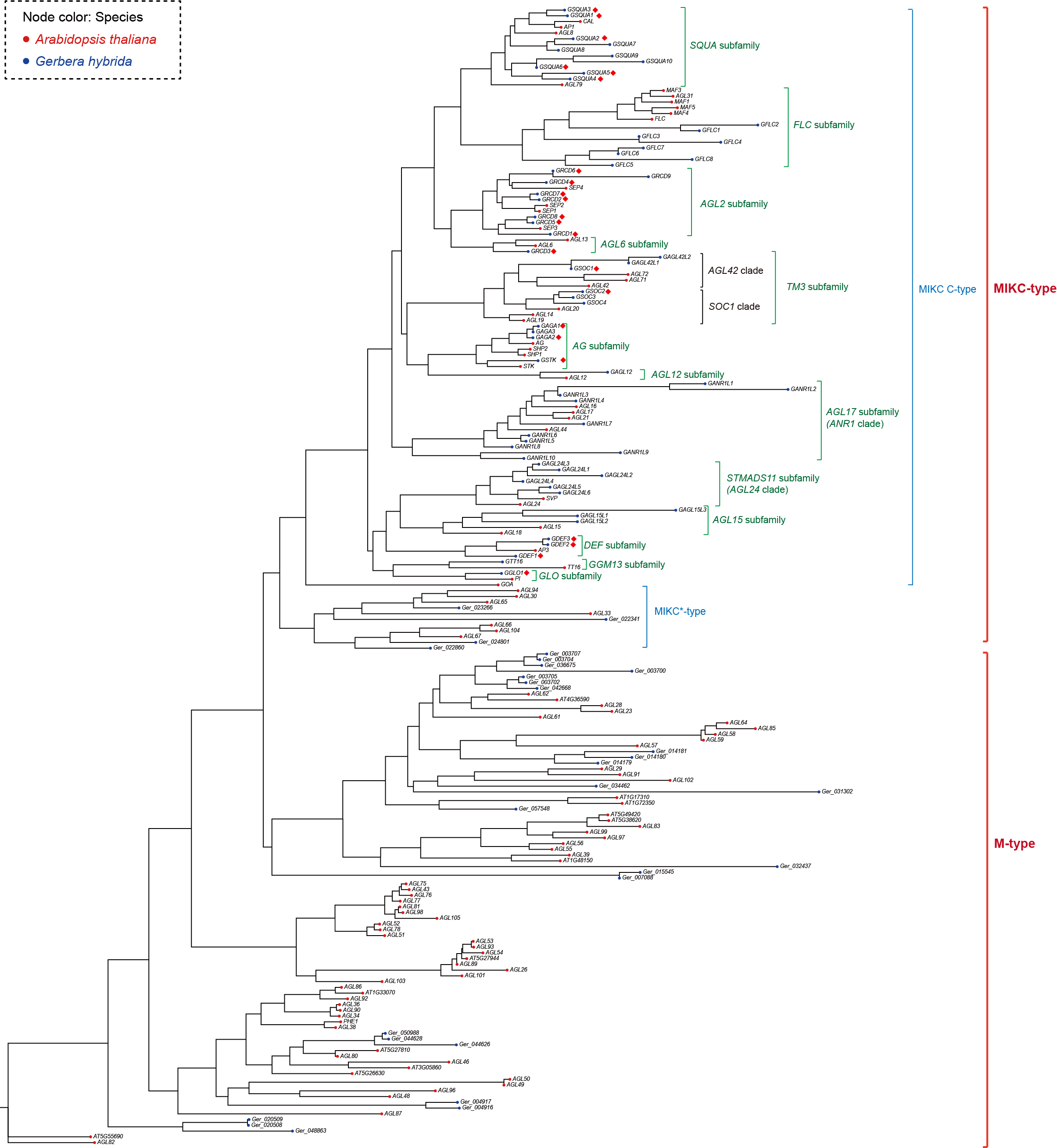

**Supplementary Figure** S7. Phylogenetic tree of MADS-box genes in *A. thaliana* and *G. hybrida*. The color of nodes shows the species of genes. The gene subfamily and some clade information are listed on the right side of gene symbols. We named the new MADS-box genes in a integrative strategy: for genes with an evidently direct orthologous gene, we name them in a like like ‘*G* + orthologous gene symbol’ ; for genes in well studied subfamilies, like *SQUA* and *AGL2* subfamily, we follow the same naming pattern of the previously studied genes; for genes in subfamilies that haven’t been studied in *G. hybrida* before, we name the genes in a pattern like ‘*G* + subfamily or clade name + (L) + number’; for genes in the MIKC*-type and M-type clade, we keep the original gene locus tag in the genome assembly. The genes marked by diamond have been recorded in previous studies.

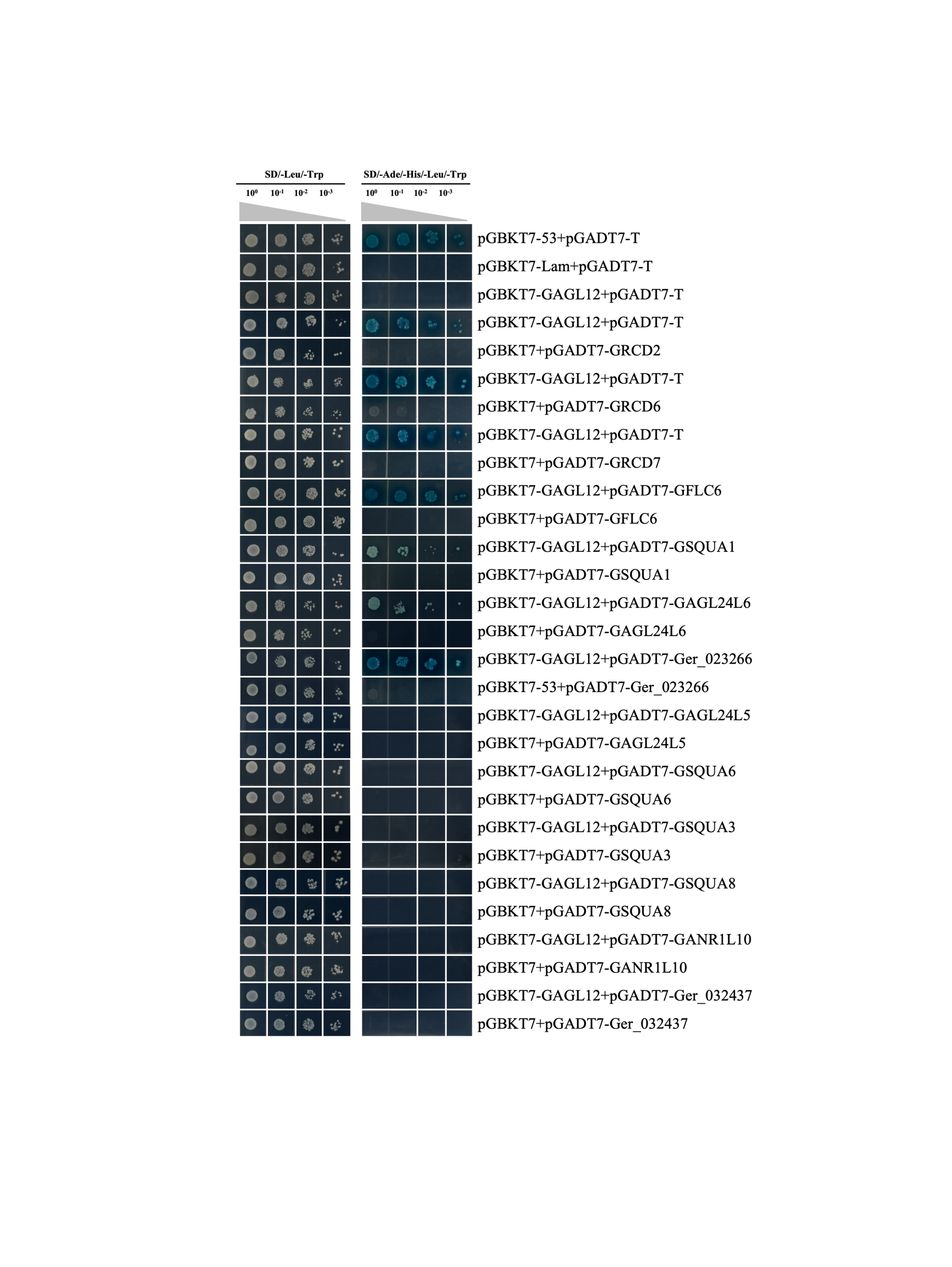

**Supplementary Figure S8.** The yeast-two hybrid result. The co-transformed yeast strains pGBKT7-GAGL12 + pGADT7-GRCD2, pGBKT7-GAGL12 + pGADT7-GRCD6, pGBKT7-GAGL12 + pGADT7-GRCD7, pGBKT7-GAGL12 + pGADT7-CFLC6, pGBKT7-GAGL12 + pGADT7-GSQUA1, pGBKT7-GAGL12 + pGADT7-GAGL24L6 and pGBKT7-GAGL12 + pGADT7-Ger_023266 grew normally on SD/-Ade/-His/-Leu/-Trp/X-α-Gal plates and turned blue, indicating that GAGL12 interacts with GRCD2, GRCD6, GRCD7, GFLC6, GSQUA1, GAGL24L6, and Ger_023266 in yeast. In contrast, the co-transformed strains pGBKT7-GAGL12 + pGADT7-GhSVPc, pGBKT7-GAGL12 + pGADT7-GSQUA6, pGBKT7-GAGL12 + pGADT7-GSQUA3, pGBKT7-GAGL12 + pGADT7-GSQUA8, pGBKT7-GAGL12 + pGADT7-ChCALb, and pGBKT7-GAGL12 + pGADT7-GhAGL26 grew normally on SD/-Leu/-Trp plates but failed to grow on SD/-Ade/-His/-Leu/-Trp/X-α-Gal plates, demonstrating that GAGL12 does not interact with GAGL24L5, GSQUA6, GSQUA3, GSQUA8, GANR1L10, or Ger_032437 in yeast.
